# ARMC1 regulates mitochondrial fatty acid oxidation through the insertase/scramblase MTCH2

**DOI:** 10.64898/2026.08.26.747330

**Authors:** Andrew Castonguay, Dariel Márquez, Andrew Natale, Royce York, Shelly Harel, Jack Cazet, Mia Pulos-Holmes, Angela Xu, Kibeom Kim, Kit Page, Mariia Burdyniuk, Jacob N. Bonner, Yari Sigal, Michael Paddy, Jin Chen, Marijn G. J. Ford, Adam Frost, Daniel Itzhak, Stefka Tyanova, Maxence Le Vasseur, Jodi Nunnari

## Abstract

MTCH2 (mitochondrial carrier homolog 2) is a noncanonical member of the solute carrier family with five transmembrane (TM) helices, localized to the mitochondrial outer membrane. MTCH2’s atypical topology creates a membrane-accessible hydrophilic groove, predicted to be necessary for its protein insertase and lipid scramblase activities. MTCH2 is linked to lipid metabolism and obesity and is required for starvation-induced mitochondrial hyperfusion. Here, we show that MTCH2 is a stable component of a complex containing the Armadillo (ARM) repeat-containing protein, ARMC1, and the DnaJ/Hsp40 chaperone protein, DNAJC11. Protein crosslinking, protein structural modeling, and molecular dynamics simulations demonstrate that the ARMC1 alpha-helical C-terminal domain (CTD) inserts into and stably interacts with the MTCH2 hydrophilic groove and blocks its lipid scramblase activity. We observe that starvation-induced mitochondrial fatty acid oxidation (FAO) is negatively regulated by the ARMC1-MTCH2 interaction. In ARMC1-deficient cells, FAO is stimulated compared to wild-type cells and lipid droplet abundance is significantly reduced. The altered lipid phenotype of ARMC1^-/-^ cells is strictly dependent on MTCH2 and is reversed by ARMC1 expression in a manner dependent on its CTD. Beyond this metabolic axis, we also identify a function for ARMC1 in regulating lysosomal distribution and autophagic flux that is independent of its CTD and interaction with MTCH2. Thus, our data support a model in which the MTCH2-ARMC1 interaction functions as a metabolic switch during starvation to regulate the balance between fat storage and fat burning.

## Introduction

Mitochondria are highly dynamic organelles that continuously adapt their metabolic output to environmental stimuli, stress, and shifting cellular states ^1, 2^. A hallmark of this adaptability is the precise coordination between mitochondrial morphology and metabolic flexibility, the ability of the cell to switch between substrates to meet fluctuating energy demands ^3,4,5^. During starvation conditions, cells undergo autophagy and a major metabolic rewiring, including increased lipid mobilization, fatty acid oxidation (FAO), and the induction of starvation-induced mitochondrial hyperfusion (SIMH), an adaptive response promoting cell survival ^6–9^. This mitochondrial plasticity is supported by a dynamic and intricate remodeling of proteins and lipids, which serve to prioritize specific biochemical pathways.

We have previously demonstrated that the Mitochondrial Carrier Homolog 2 (MTCH2) is a critical regulator of mitochondrial morphology and specific mediator of SIMH in a manner dependent on the bioactive lipid lysophosphatidic acid, suggesting a role for MTCH2 in lipid metabolism ^10^. MTCH2 is a structurally atypical mitochondrial outer membrane (MOM) solute carrier protein belonging to the SLC25 superfamily ^11^, but unlike other SLC25 proteins, it possesses five, rather than the characteristic six transmembrane (TM) helices ^12,13^. The ‘missing’ TM helix creates a lipid-accessible hydrophilic groove that has been proposed to facilitate phospholipid scrambling ^14,15^ and protein insertase activity ^13,16^.

MTCH2 has been reported to be a component of a complex containing the Armadillo (ARM) repeat-containing protein ARMC1, the DnaJ/Hsp40 chaperone protein DNAJC11, and the metaxin protein MTX1 ^12^. ARMC1 is dually localized to mitochondria and the cytosol ^17,18^ and has been reported to partition between a MTCH2-DNAJC11 complex and a mitochondrial distribution complex containing mitochondrial trafficking tethers MIRO1/2, mitochondrial fission regulator (MTFR) proteins, and TMEM11^12^, suggesting tight regulation of MTCH2 molecular function. Within these complexes, the ARMC1 heavy metal-associated–like (HMAL) domain is predicted to interact with MIRO or the cytosolic C-terminal domain of DNAJC11 ^12^. However, ARMC1 recruitment to mitochondria is strictly dependent on its C-terminal domain (CTD), and AlphaFold models predict that the ARMC1 CTD engages the MTCH2 hydrophilic groove ^12,17^.

Here, we report that the MTCH2-ARMC1 interaction regulates lipid homeostasis. Using a combination of proteomics, molecular modeling, molecular dynamics simulations, biochemical assays, and functional metabolic analysis, we show that MTCH2 lipid scramblase activity is dependent on the accessibility of its hydrophilic groove and negatively regulated by the recruitment of the ARMC1 CTD. ARMC1 deletion or truncation of its CTD rewires lipid metabolism in cells by enhancing mitochondrial FAO under starvation conditions, causing depletion of lipid droplets (LDs), in a manner strictly dependent on MTCH2. Beyond this metabolic axis, we also identify a CTD-independent role of ARMC1 in regulating lysosomal distribution and autophagic flux. Collectively, our data highlight a metabolic switch that coordinates lipid metabolism and autophagy to fine-tune cellular metabolic flexibility under starvation.

## Results

### MTCH2 contains a cytosol-accessible hydrophilic groove that recruits ARMC1 CTD amphipathic helix to form a MTCH2-ARMC1-DNAJC11 supramolecular complex

Canonical SLC25 mitochondrial carriers are defined by their highly specialized structure and shared topology, which allows for a unique rocking mechanism that alternates access between the mitochondrial matrix and the intermembrane space to transport metabolites ^11^. However, unlike other solute carrier proteins, MTCH2 localizes to the outer mitochondrial membrane (OMM) and a recent cryo-EM study of chimeric MTCH2 revealed that it contains only five instead of the canonical 6 transmembrane helices of the SLC25 family^13^, confirming the earlier predicted model (Fig. 1A, left). This topology creates a membrane-accessible hydrophilic groove predicted to be required for its lipid scramblase and protein insertase activities ^14,15^. Given its functional significance, we tested MTCH2 membrane topology *in situ* using proteinase K protection on mitochondria isolated from MTCH2^-/-^ cells stably expressing an N- or C-terminal 3xFlag tag. Expression and proper localization of recombinant MTCH2 was validated by immunoblot and IF microscopy, respectively (Supplementary Fig. 1-1A-B). Treatment of mitochondria with proteinase K completely digested the epitopes of both OMM protein TOM70 and MTCH2 N-terminal 3xFlag proteins, thereby confirming its localization to the OMM (Fig. 1B). In contrast, a residual ∼15kDa Flag-containing fragment was protected for C-terminally tagged MTCH2, consistent with the localization of MTCH2 C-terminus within the mitochondrial intermembrane space (Fig.1B, arrow).

**Fig. 1:**
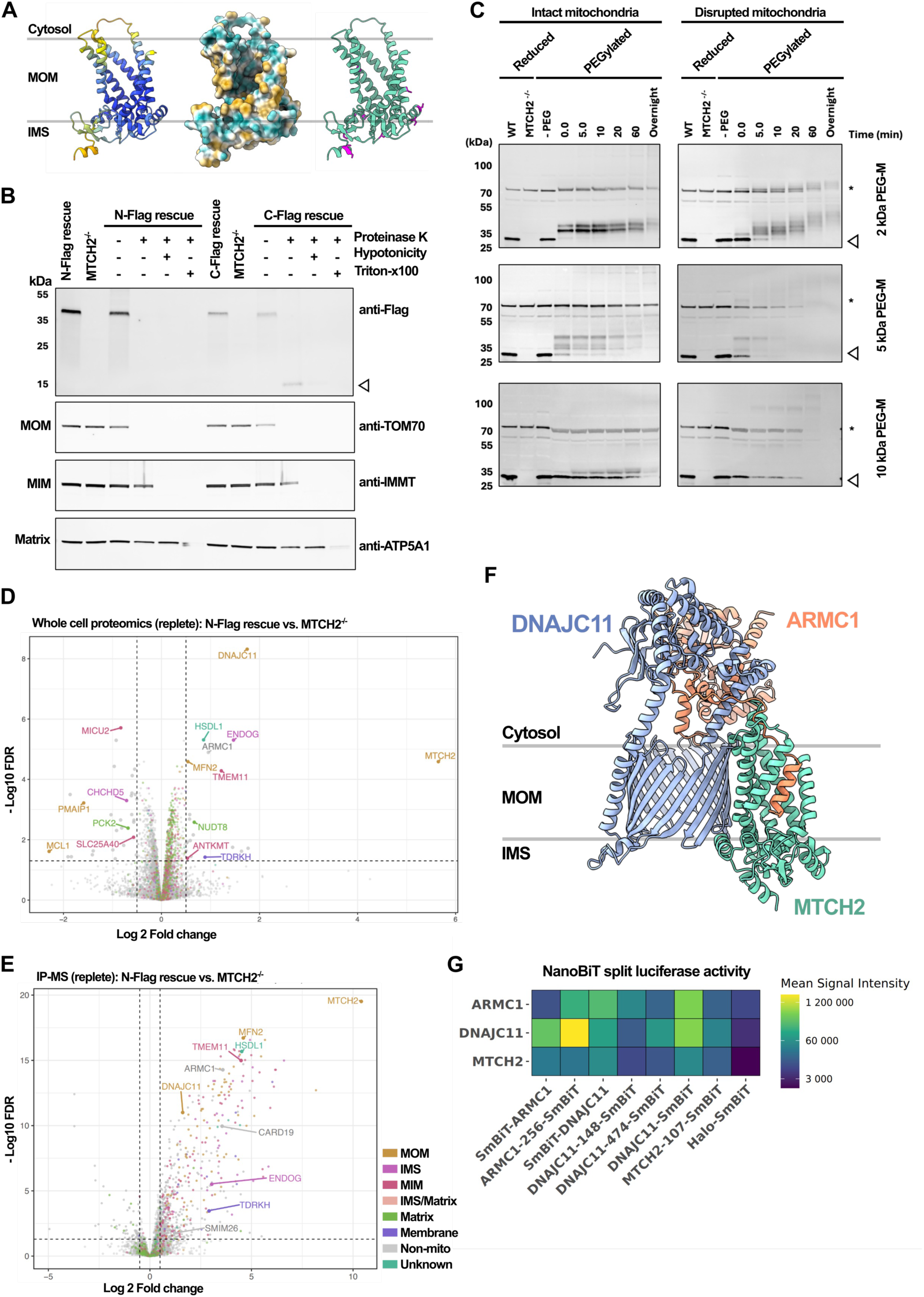
MTCH2 topology and interactome support an MTCH2-ARMC1-DNAJC11 complex. (A) MTCH2 ribbon structure modeled by AlphaFold colored by pLDDT score (left), space-filling model colored by hydrophobicity for positioning of bilayer (center), and MTCH2 AlphaFold ribbon structure highlighting cysteine residues in magenta (right). (B) Proteinase K protection assay of enriched mitochondria from rescue cell lines expressing MTCH2 with N- or C-terminal 3xFlag tag. Sub-compartment assignment was given by comparison of anti-Flag migration pattern in comparison to markers of the mitochondrial outer membrane (TOM70), inner mitochondrial membrane (IMMT), or matrix (ATP5A1). (C) WT mitochondria modified by low (2 kDa, top row), medium (5 kDa, middle), or high (10 kDa, bottom row) molecular weight PEG maleimide. Modification of MTCH2 was observed by immunoblot in comparison to unmodified WT and MTCH2^-/-^ controls. Arrowhead designates unmodified MTCH2 at approximately 33 kDa whereas asterisk denotes nonspecific, cross-reacting species. (D) Volcano plot comparing whole cell proteomes of N-terminally tagged MTCH2 rescue cells relative to MTCH2^-/-^ cells under replete conditions (FDR < 0.05, fold-change cutoff set to 0.5, n = 5 replicates). (E) Enrichment of proteins co-purified with MTCH2 depicted by volcano plot (N-terminal 3xFlag-tagged MTCH2, right, relative to WT, left) with Flag tag used as bait under under replete conditions (FDR < 0.05, fold-change cutoff set to 0.5, n = 5 replicates). Proteins co-dependent with MTCH2 labeled. (D) and (E) Mitochondrial proteins colored by compartment and labeled, non-mitochondrial proteins depicted in grey. (F) Boltz-1/2 molecular modeling of heterotrimeric complex consisting of MTCH2 (teal), ARMC1 (coral), and DNAJC11 (lavender). (G) NanoBiT split luciferase activity assay of cell cultures transiently co-expressing chimeric LgBiT-tagged ARMC1, DNAJC11, or MTCH2 with corresponding SmBiT tagged potential pairs. Total luminescence for each pair was plotted by heatmap (n = 3). Combination with Halo-SmBiT served as negative control for nonspecific interactions.

To further test the topology of endogenous MTCH2, we leveraged the polarized distribution of cysteines in the MTCH2 structure (Fig. 1A, right) via a PEGylation assay as previously described ^19^ (Fig. 1C). We observed an upward shift in MTCH2 mass in intact mitochondria incubated with small molecular weight cysteine-reactive PEG-maleimide molecules (2 or 5kDa) that readily diffuse through VDAC pores (Fig. 1C), but not with larger, membrane impermeable molecules (10kDa) ^19,20^. Disruption of mitochondria by treatment with SDS bypassed the VDAC molecular weight cutoff and allowed modification of IMS-facing cysteines in MTCH2 by high molecular weight PEG-maleimides. Consistent with our protease protection assay, these results suggest that the majority of cysteine residues are facing the IMS. Taken together, our data are in agreement with the recently determined MTCH2 structure, confirming its N-out/C-in membrane topology and the resulting cytosol-accessible hydrophilic groove.^12,14–16^.

MTCH2 was recently identified as part of a complex containing ARMC1, DNAJC11, and MTX1 ^12^. We used unbiased mass spectrometry to compare the proteome of MTCH2^-/-^ cells and MTCH2^-/-^ cells rescued with N-terminal 3xFlag-tagged MTCH2 to determine the interdependence of MTCH2 complex components (Fig. 1D). Since MTCH2 is essential for SIMH, we also assessed interdependence after 5h starvation (HBSS), a condition which induces SIMH (Supplementary Fig. 1-2B). Depletion of MTCH2 resulted in a concomitant reduction in DNAJC11 and ARMC1, but not MTX1 abundance independent of growth condition. Several of the co-depleted proteins, including ARMC1, DNAJC11, ENDOG, and TMEM11 have been previously associated with SAM and MICOS complexes ^21^. Prior reports have uncovered the co-depletion of ARMC1 and DNAJC11 with MTCH2, and public databases cataloged the interdependency of these proteins with MICOS/MICOS-interacting proteins ^12,16,22^. We asked whether the stability of DNAJC11 or MTCH2 was reciprocally interdependent with ARMC1 (Supplementary Fig. 1-3A-D). Interestingly, neither DNAJC11 nor MTCH2 were co-depleted upon loss of ARMC1. However, ARMC1 was essential for the stability of the MTFR proteins as previously reported ^12^ (Supplementary Fig. 1-3 and 1-4).

We characterized the MTCH2 interactome using affinity enrichment mass spectrometry (AE-MS). MTCH2 was immunopurified from crosslinked whole-cell lysate prepared from wild type (WT) cells or MTCH2^-/-^ cells rescued with MTCH2 N-terminal 3xFlag tag and the eluates were analyzed by label-free quantitative mass spectrometry (Fig. 1E and Supplementary Fig. 1-2C). Mitochondrial proteins co-depleted with MTCH2 in MTCH2^-/-^ cells were also enriched in the MTCH2 interactome, including ARMC1, DNAJC11, and TMEM11.

The organization of the proposed MTCH2-containing heterotetramer^12^ was interrogated by querying publicly available crosslinking mass spectrometry (XL-MS) data, which provides residue-to-residue resolution and *in situ* structural insights ^23^. Intra- and intermolecular crosslinks were identified for ARMC1, DNAJC11, MTCH2, and MTX1 (Supplementary Fig. 1-5A). However, MTX1 and ARMC1 formed intermolecular crosslinks with the same residues in DNAJC11, indicative of mutually exclusive interactions. Structural modeling of a MTX1-DNAJC11 heterodimer using AlphaFold2 suggested that MTX1 binds the DNAJC11 C-terminus. In agreement with the XL-MS data, mapping of the molecular distances between crosslinked residues in the heterodimer showed that 6 out of 10 intermolecular crosslinks were within the 40 Å Cα–Cα maximum physical distance constraint (Supplementary Fig. 1-6A-C, G). In contrast, in an AlphaFold2 model of an ARMC1-DNAJC11-MTCH2-MTX1 complex, MTX1 is displaced from the DNAJC11 C-terminus and only 1 out of 10 intermolecular crosslinks was within the maximum physical distance constraint. Intermolecular crosslinks were also formed between MTX1 and MIRO1, MIRO2, and TMEM11, of the second ARMC1-containing complex on the OMM suggesting a degree of dynamic rearrangement (Supplementary Fig. 1-5A).

We focused our analysis on the 50 unique crosslinks identified for MTCH2, DNAJC11, and ARMC1. Forty-six of crosslinks were within the physical distance constraint, strongly supporting our predicted models (Supplementary Fig. 1-5 and Supplementary Table 1) ^23^. Using AlphaFold2 and Boltz-1, we generated structural models of a tripartite complex between ARMC1, DNAJC11, and MTCH2 (Fig. 1F). Conformational differences between the models point to potential substantial flexibility of DNAJC11 and ARMC1, however the orientation of DNAJC11 and MTCH2 relative to the membrane remained consistent and the same sets of residues were identified as high-confidence interfaces (Supplementary Fig. 1-5C-F). One interface is the ARMC1 CTD amphipathic helix within the membrane-spanning hydrophilic groove of MTCH2, which completes the 3-fold pseudo-symmetry of the SLC25 protein family ^11^. Consistent with our models, inter-protein crosslinks were observed between ARMC1 and the C-terminal domain of DNAJC11 and between a flexible loop of the DNAJC11 C-terminus and the MTCH2 hydrophilic groove. However, this crosslink was only satisfied in the AF2, but not Boltz-1 model, likely highlighting the flexible nature of this loop (Supplementary Fig. 1-5C-F). Additional inter-protein crosslinks were found between ARMC1 CTD and MTCH2 hydrophilic groove, in close alignment with our tripartite structural model (Supplementary Fig. 1-5E-F).

We tested the predicted ARMC1-DNAJC11 interaction via yeast two-hybrid, co-expression/purification, and split luciferase assays. Two-hybrid analysis (Supplementary Fig. 1-7A-B), indicated that ARMC1 interacts with the C-terminus of DNAJC11 but not the DNAJ domain, consistent with our predicted models and crosslinking mass spectrometry data (Supplementary Fig. 1-7A-D). Introduction of amino acid substitutions targeting the predicted interface – specifically, reversal of charged residues at positions R532, R547, and R554 of DNAJC11 – disrupted this interaction (Supplementary Fig. 1-7C). Reciprocally, charge reversal substitutions in the ARMC1 HMAL domain (E162 and E163) or the acidic linker PDYLPEDE motif also ablated interaction with the DNAJC11 C-terminus (Supplementary Fig. 1-7C). Our modeling predicted that the Y240 sidechain in the ARMC1 PDYLPEDE motif packs into a hydrophobic pocket formed by residues F531 and H536 of the DNAJC11 C-terminus. Substitution of Y240 with alanine was sufficient to disrupt interaction by yeast two-hybrid (Supplementary Fig. 1-7C).

In addition, expression of full-length ARMC1 in bacteria was dependent on co-expression of the DNAJC11 C-terminus, which produced a stable heterodimer (Supplementary Fig. 1-8A-F). In contrast, an ARMC1 CTD-truncation mutant was stably expressed in the absence of DNAJC11, suggesting that presence of ARMC1 CTD requires complex formation with DNAJC11 for protein stability (Supplementary Fig. 1-8G). Taken together, these results support an interaction of ARMC1 with DNAJC11 C-terminus, characterized by charge-charge interactions with the ARMC1 HMAL domain and its acidic linker and stabilized via an interaction of ARMC1 Y240 within the hydrophobic pocket of the DNAJC11 C-terminus.

We further probed MTCH2-ARMC1-DNAJC11 interaction using a low affinity split luciferase assay ^24^. Luciferase activity was detected in human cells co-expressing recombinant DNAJC11 and ARMC1 split luciferase pairs, confirming the molecular proximity of both proteins (Fig. 1G and Supplementary Fig. 1-9C-E). Moderate signal was also observed in cells co-expressing LgBiT-DNAJC11 and DNAJC11-SmBiT (or with the reciprocal SmBiT/LgBiT pair) and in cells co-expressing ARMC1 LgBiT/SmBiT split luciferase pairs (Fig. 1G and Supplementary Fig. 1-9C-E), suggestive of self-interaction, consistent with XL-MS analyses (Supplementary Fig. 1-5A). Luminescence was detected, albeit at lower levels, in cells co-expressing LgBiT-MTCH2 and SmBiT-containing ARMC1 or DNAJC11 (Fig. 1G and Supplementary Fig. 1-9F). Low luminescence levels could be attributable to suboptimal positioning of the split-luciferase tags and reduced complementation efficiency. No significant differences were observed when recombinant split luciferase pairs were coexpressed and luminescence measured under conditions of starvation (Supplementary Fig. 1-9D). Taken together, these results strongly support the formation of a tripartite complex composed of MTCH2, DNAJC11, and ARMC1.

### Mitochondrial localization of ARMC1 is partially dependent on MTCH2

ARMC1 is dually localized between the cytosol and the OMM, a process reportedly mediated by the interaction of its CTD with TMEM11 and MTCH2 ^12,17^. We asked whether ARMC1 partitioning was dependent on MTCH2 or changed under conditions inducing SIMH. We assessed the distribution of full-length or CTD truncated ARMC1 via immunofluorescence (IF) microscopy utilizing lipid nanoparticle (LNP)-mediated mRNA delivery at near-endogenous levels (Fig. 2A-B and Supplementary Fig. 2-1). IF microscopy revealed that full length ARMC1 was significantly depleted from mitochondria in MTCH2^-/-^ cells (Fig. 2B), whereas the CTD truncation mutant was strictly confined to the cytoplasm, confirming the necessity of the C-terminus for mitochondrial targeting (Supplementary Fig. 2-1). These findings were orthogonally validated by immunoblotting and unbiased proteomics analyses of cytosolic and mitochondrial fractionations (Fig. 2C, Supplementary Fig. 2-2).

**Figure 2:**
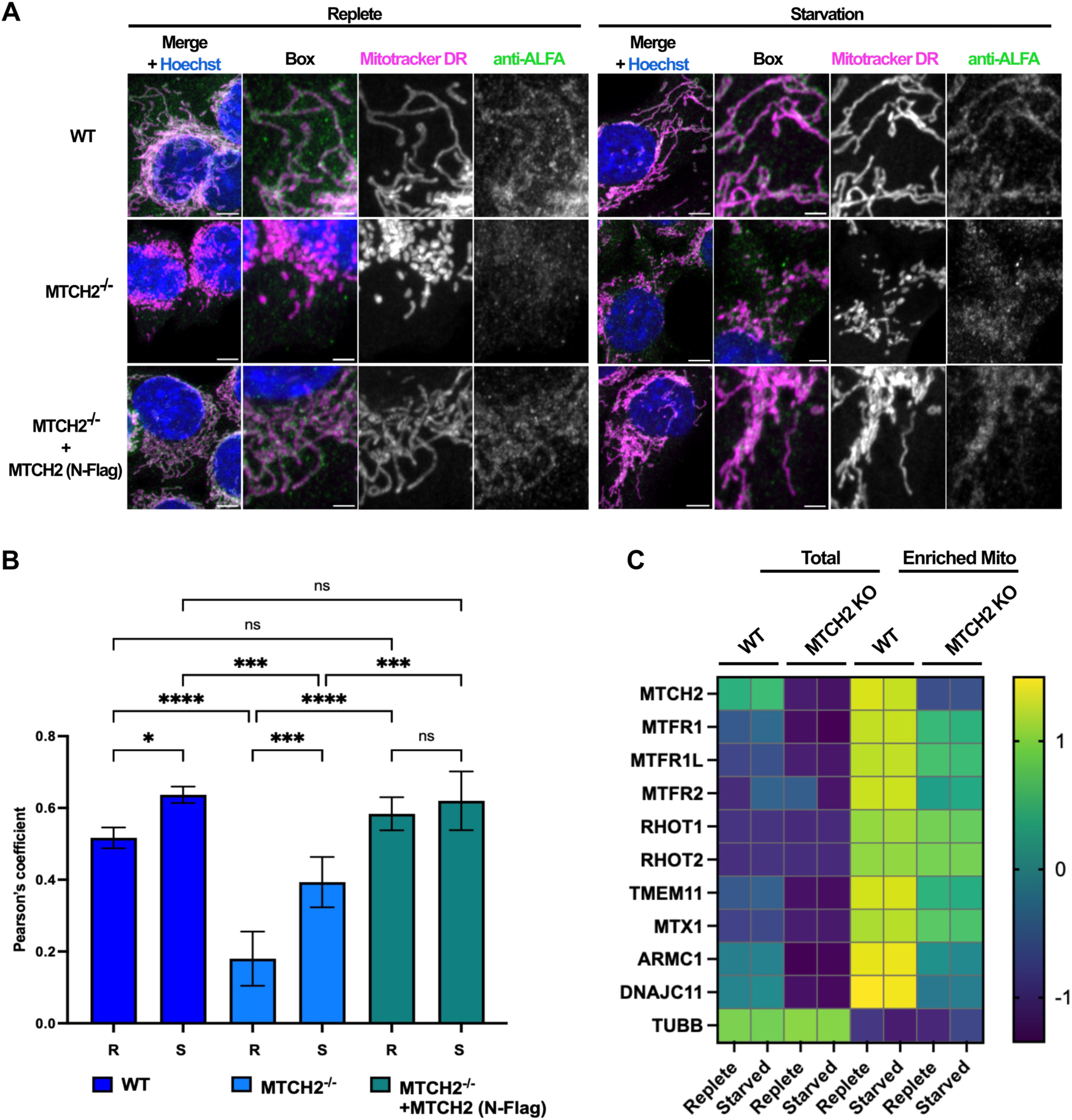
Mitochondrial localization of ARMC1 is partially conditional on MTCH2. Subcellular localization was assessed under nutrient-replete and starved conditions in presence and absence of MTCH2. (A) Representative images of WT, MTCH2^-/-^, and MTCH2^-/-^ + MTCH2 (N-Flag) rescue cells transiently expressing ALFA-ARMC1-FL. The construct was delivered via LNPs, and cells were evaluated under replete and starved conditions following MitoTracker Deep Red and anti-ALFA staining. Main scale bars = 5 µm; inset box scale bar = 2 µm. (B) Quantification of ALFA-ARMC1-FL mitochondrial localization for the conditions described in (A), assessed via Pearson’s correlation coefficient. Statistical significance was determined using a one-way ANOVA followed by Tukey’s test. (C) WT and MTCH2^-/-^ cells were cultured under replete and starved conditions, fractionated, and whole cell and mitochondrial-enriched fractions analyzed by unbiased proteomics (n = 5 replicates/condition). Z-score was calculated and relative protein abundance depicted via heatmap for ARMC1-interacting proteins versus TUBB control.

We also examined whether ARMC1 distribution was altered during starvation, a condition causing MTCH2-dependent mitochondrial hyperfusion ^10^. Image analysis suggested a slight increase in mitochondrial ARMC1 during starvation in MTCH2^-/-^ cells (Fig. 2B). Given that the quantitative IF microscopy can be limited by fixation and permeabilization artifacts affecting epitope accessibility, we orthogonally tested this observation by immunoblot and unbiased proteomics of biochemical fractions and found no significant difference in mitochondrial enrichment during starvation (Fig. 2C, Supplementary Fig. 2-2). Collectively, these data demonstrate that while MTCH2 is a critical determinant of ARMC1 stability and mitochondrial enrichment, it is not strictly required for ARMC1 mitochondrial compartmentalization.

### Proteomics analyses support separation of functions between cytosolic and mitochondrial pools of ARMC1

To assess the roles of cytosolic and mitochondrial ARMC1 pools, we performed unbiased proteomics and redundancy analysis comparing full-length (ARMC1-FL) and C-terminally truncated (ARMC1-trunc) overexpression. While both constructs rescued cytosolic MTFR abundance in ARMC1^-/-^ cells, only ARMC1-FL expression significantly suppressed mitochondrial translation and oxidative phosphorylation signatures (Supplementary Figs. 2-3 to 2-5). This proteomic footprint suggests a spatially segregated dual role, wherein the mitochondrial pool of ARMC1 acts to constrain baseline metabolic capacity. Given the systemic suppression of mitochondrial signatures, we investigated whether ARMC1 serves as a negative regulator of lipid utilization and mitochondrial FAO.

### MTCH2 *in silico* scramblase activity is blocked by engagement of the ARMC1 C-terminus

MTCH2 has demonstrated scramblase activity for neutral phospholipids based on proteoliposome reconstitution experiments ^15^, and *in silico* studies suggest the scrambling pathway traverses MTCH2’s membrane-spanning hydrophilic groove ^14,15^. Building on this work, we used in silico analysis to investigate how MTCH2 interacts with multiple lipid types, and whether the ARMC1-MTCH2 interaction altered scramblase activity. We employed coarse-grained molecular dynamics (CG-MD) to probe lipid behavior at longer timescales while using all-atom molecular dynamics (AA-MD) simulations to examine the details of protein-protein and protein-lipid interactions with greater chemical realism. The protein fold and the architecture of the hydrophilic groove have been validated by a recent cryo-EM structure of MTCH2, demonstrating high consistency with predicted models used in our simulations, except for the terminal and loop regions of lower confidence ^13^.

Our initial AA-MD simulations using the MTCH2 AlphaFold2 model validated that the hydrophilic groove serves as the primary scrambling pathway, with lipids infiltrating from the cytosolic leaflet within nanoseconds (Fig. 3A). The groove accommodates 4-5 phospholipids at once, reoriented so that their headgroups interact with protein residues lining the sides and base of the groove while some fatty acid tails realign with the bilayer midplane. To assess the potential for lipid scrambling, we ran CG-MD Martini3 simulations with six protein copies per system in bilayers roughly approximating the phospholipid diversity of the OMM ^25^ (Supplementary Fig. 3-1A). We observed numerous MTCH2 mediated scrambling events at a rate of ∼1 lipid/μs per protein copy, involving all lipid species at different individual rates (Fig. 3B upper). Alternatively, when the ARMC1 CTD was complexed with MTCH2 no lipid scrambling events were observed, suggesting that ARMC1 can directly regulate MTCH2 scramblase activity via sterically blocking the pathway (Fig. 3B lower).

**Figure 3:**
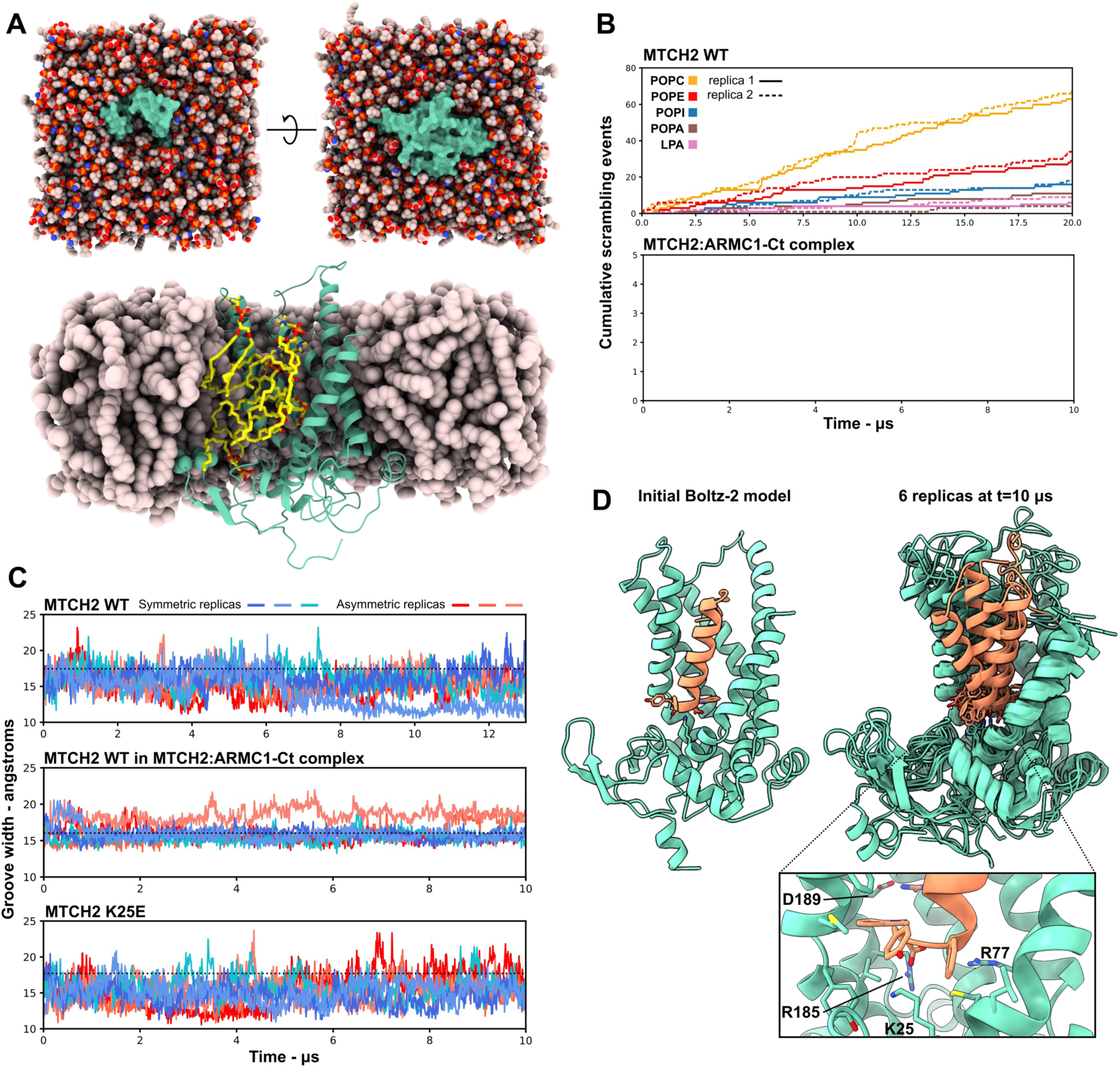
MTCH2 lipid scrambling may be regulated by ARMC1 occupancy of the hydrophilic groove. (A) Visualization of the configuration of a MD simulation of the AF2 model of MTCH2 embedded in a phospholipid bilayer, as viewed from the cytoplasmic face (top left), IMS face (top right) and cut away through the membrane (bottom). MTCH2 is shown as a teal Van der Waals surface (top) or ribbon representation with sidechains of residues Ile43 and Phe44 shown as spheres (bottom). Most lipid heavy atoms are shown as spheres; at bottom, lipids that have entered the MTCH2 hydrophilic groove are shown in stick representations, with carbon atoms colored yellow. (B) CG-MD cumulative total lipid scrambling events (counting both cyto-to-IMS and IMS-to-cyto leaflet transitions) by lipid type over the length of two independently prepared simulation trajectories (solid/dashed lines) for each condition, each containing 6 copies of MTCH2 or MTCH2 WT:ARMC1-Ct complex embedded in a phospholipid bilayer. Each lipid type is indicated by lines of a different color. Scrambling was assessed by using a spectral clustering algorithm to assign each lipid to one of the two leaflets at each saved simulation frame and tracking when lipids changed leaflet assignments (C) The width of the hydrophilic groove over simulation time across all replicas, calculated as the distance between Cα atoms of MTCH2 residues 14 and 224. The black dotted line indicates the distance in the respective initial model. Blue traces denote simulations that started with a symmetric POPA:POPC distribution in each leaflet, while red traces denote simulations that started with all POPA in the outer leaflet (asymmetric). In both cases the overall ratio was 20:80 POPA:POPC. (D) Representation of the Boltz-2 structural model of the MTCH2 WT:ARMC1-Ct complex used to prepare simulations (left) and six aligned replicas of the same after 6 μs AA-MD (right) and a zoomed inset of a single replica (bottom). MTCH2 is shown in light green and ARMC1 in orange; select MTCH2 sidechains are labeled.

Subsequently, in order to resolve atom level details of protein-protein and protein-lipid interactions we prepared long all-atom simulations of MTCH2 WT, MTCH2:ARMC1-CTD, and MTCH2 K25E, a mutation that flips a central charge within the groove and is reported to increase the protein insertase rate^16^, but does not disrupt the overall structure^13^. Each simulation embedded 3 copies of the respective Boltz-2 model in simple phospholipid bilayers (20:80 POPA:POPC bilayers, symmetric or asymmetric leaflet distribution, Supplementary Fig. 3-1B). In all simulations, the overall MTCH2 fold remained consistently stable, with RMSD relaxation of plateauing after 2-4 microseconds and remaining roughly constant thereafter, with adaptations localized to flexible loops and the C-terminus (Supplementary Fig. 3-2A,B). In MTCH2 WT and K25E the hydrophilic grooves narrowed slightly with respect to the input models, but largely remained open and dynamic for the duration of trajectories, while in the MTCH2:ARMC1-CTD complex the groove maintained roughly the same mean width with a much narrower distribution as a result of ARMC1 contacts (Fig. 3C, Supplementary Fig. 3-2C). In simulations of the complex, ARMC1 remained anchored to the base of the groove via a combination of hydrophobic contacts with the final ‘FYW’ motif and salt bridges between the ARMC1 C-terminal carboxylate and variously K25, R77, & R185 of MTCH2 as well as between ARMC1 R278 and MTCH2 D189 (Fig. 3D, Supplementary Fig. 3-2D). We observed two PA-only lipid translocation events for MTCH2 WT and six combined PA/PC events for MTCH2 K25E, while the MTCH2:ARMC1-CTD complex completely blocked lipids from entering the groove or translocating (Supplementary Fig. 3-3A-F). We also observed that distributions of lipid headgroup interactions with protein surfaces in and around the groove were markedly different between MTCH2 WT and MTCH2 K25E, which in particular appears to restrict PA from occupying a site in the deepest part of the groove (Supplementary Fig. 3-4A-D). The apparent increase in lipid translocation rate for K25E parallels the reported increase in protein insertase activity^16^, and suggests both mechanisms depend on similar electrostatic interactions. The CG-to-AA scrambling rate difference for WT MTCH2 (∼1 lipid/μs in CG vs ∼0.05 lipid/μs in AA) is expected based on the smoothed energy landscape of CG potentials. While our CG measured rate is roughly an order of magnitude slower than previously described CG simulations of MTCH2^14,15^, our simulations differ by containing anionic in addition to neutral phospholipids, which we speculate could alter bulk lipid binding and translocation rates through the charged groove. Overall, our results confirm the structural role of the MTCH2 groove in lipid scrambling and identify a potential steric blocking mechanism by which the CTD of ARMC1 regulates this activity.

### The ARMC1 C-terminus regulates mitochondrial FAO and lipid droplet abundance in a MTCH2-dependent manner

MTCH2 is linked to human obesity and has been shown to regulate lipid utilization ^10,26–28^. To investigate whether the MTCH2 hydrophilic groove and the MTCH2-ARMC1 interaction are functionally linked to fatty acid metabolism, we quantified lipid droplet (LD) abundance in MTCH2^-/-^ and ARMC1^-/-^ cells under replete and nutrient starvation, a condition that increases LD abundance^39^. In agreement with prior findings^10,28^, MTCH2^-/-^ cells exhibited markedly higher lipid droplet (LD) abundance under both nutrient-replete and starvation conditions. The levels of LDs found in MTCH2^-/-^ cells under both conditions are comparable to those observed in wild-type cells during starvation (Fig.4A-B). This phenotype was reversed upon reintroducing MTCH2 expression exogenously (Fig. 4A-B). Conversely, ARMC1^-/-^ cells exhibited a substantial reduction in LD content across all nutrient conditions. This phenotype was successfully rescued by ectopic expression of full-length ARMC1, with the most pronounced recovery observed during starvation. However, expression of a C-terminally truncated ARMC1 mutant did not rescue the LD content, phenocopying ARMC1^-/-^ (Fig 4A-B).

**Figure 4:**
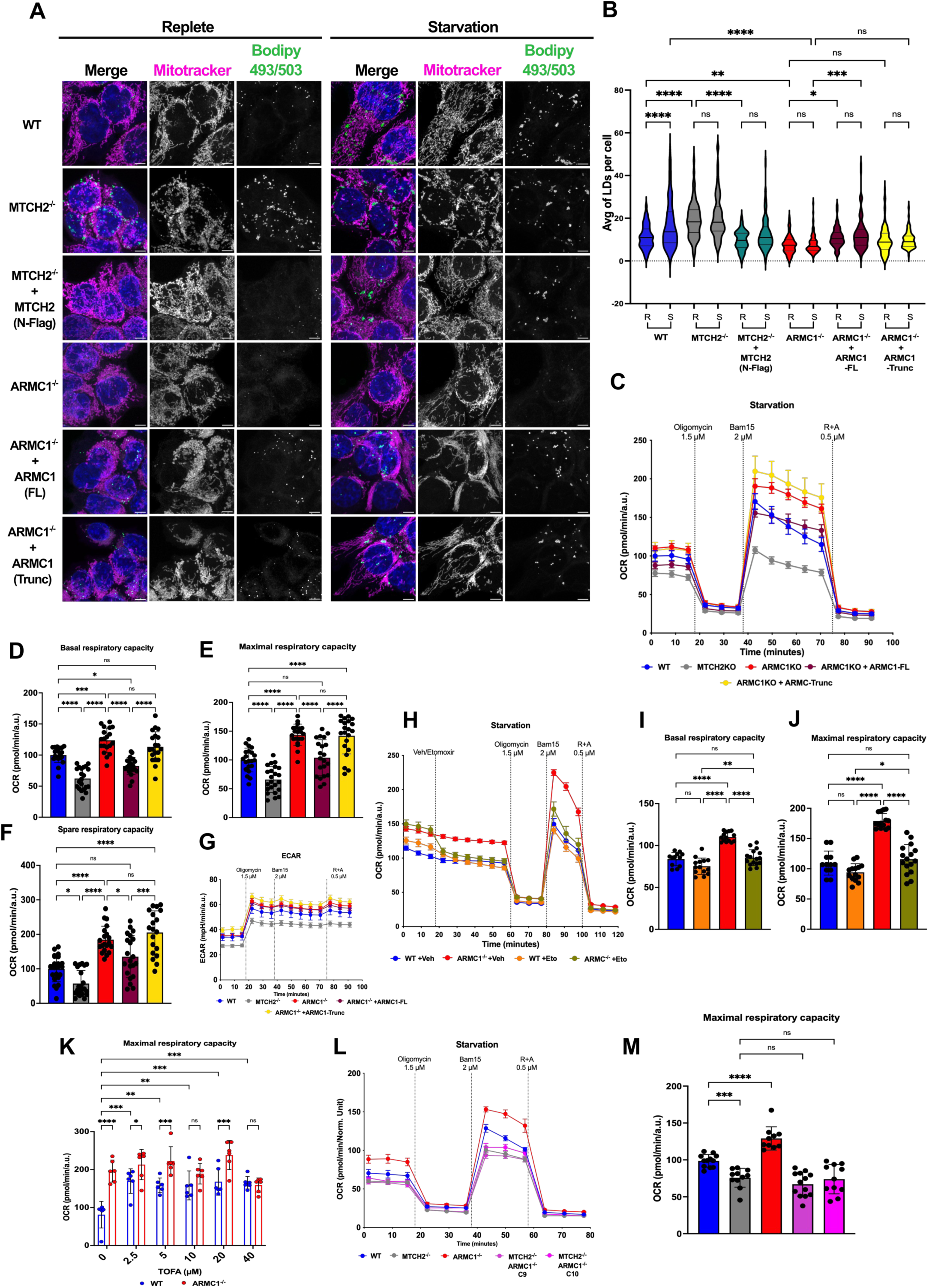
ARMC1 regulates FAO and LD abundance in a manner dependent on its C-terminus and MTCH2. (A) Mitotracker Deep Red, Bodipy 493/503, and Hoechst staining during Replete and starvation conditions of HCT116 WT cells, MTCH2^-/-^ cell line, a MTCH2^-/-^ stable line expressing MTCH2 N-terminus Flag-tagged (N-Flag), ARMC1^-/-^ cell line, an ARMC1^-/-^ stable line expressing ARMC1 full length (FL), and an ARMC1^-/-^ stable line expressing ARMC1 with a 30 amino acid C-terminus truncation (Trunc). Scale bars = 5 µm. (B) Violin plots for the quantification of Lipid Droplets for the cell lines described above. Statistical significance was evaluated by two-way ANOVA followed Tukey’s test. (C) Oxygen consumption rates during starvation for HCT116 WT cells, MTCH2^-/-^, ARMC1^-/-^, ARMC1^-/-^ stable line expressing ARMC1 full length (FL), and an ARMC1^-/-^ stable line expressing ARMC1 with a 30 amino acid truncation in the C-terminus domain (Trunc). (D) Basal Respiratory capacity using the mid-time point after assay initialization. (E) Maximal Respiratory capacity using the mid-time point after Bam15 injection. (F) Spare Respiratory capacity. (G) Extracellular acidification Rate during starvation. All seahorse values were normalized to the WT average and show ±SEM. All data points were combined from at least three independent experiments. Statistical significance was assessed by one-way ANOVA followed by Tukey’s HSD test. (H) Substrate oxidation assay during starvation to measure OCR under etomoxir (4 µM) or vehicle (Media) treatment for HCT116 WT and ARMC1^-/-^ cells. (I) Basal Respiratory capacity for the substrate oxidation assay using the mid-time point after assay initialization. (J) Maximal Respiratory capacity for the substrate oxidation assay under etomoxir using the mid-time point after Bam15 injection. Values were normalized to the WT average. All data points were combined from two independent experiments. Statistical significance was assessed by two-way ANOVA followed by Tukey’s test. (K) Maximal Respiration of a dose response of TOFA assessed by Seahorse during starvation. Values represent the mid-time point after Bam15 injection and are normalized to the WT average. Error bars show ±SEM. Statistical significance was assessed by two-way ANOVA followed by Tukey’s test. All data points were combined from at least two independent experiments. (L) Oxygen consumption rates during starvation for HCT116 WT cells, MTCH2^-/-^, ARMC1^-/-^, and two different clones (C9 and C10) of the double mutant ARMC1^-/-^ + MTCH2^-/-^. (M) Percentage of Maximal Respiratory capacity, values represent the mid-time point after Bam15 injection and are normalized to the WT average of two independent experiments. Error bars show ±SEM. Statistical significance was assessed by one-way ANOVA followed by Tukey’s test. Significance is assessed by *P<0.05, **P<0.01, ***P<0.001, ****P<0.0001, and ns = not significant.

Next, we decided to explore the potential mechanistic link between LD storage and cellular oxygen consumption rate (OCR) using Seahorse XF metabolic profiling assays since LDs serve as one of the major energetic reservoirs. Consistent with earlier findings^10^, MTCH2^-/-^ cells exhibited diminished metabolic efficiency marked by decreased basal, maximal, and spare respiratory capacities and impaired glycolytic flux under both nutrient-replete and starvation conditions (Fig. 4C-F, Supplementary Fig. 4-1 A-B). This simultaneous defect in both oxidative phosphorylation (OXPHOS) and glycolysis underscores the critical, global role of MTCH2 in maintaining cellular energetic homeostasis. Conversely, ARMC1^-/-^ cells displayed enhanced metabolic efficiency marked by a significant increase in basal, maximal, and spare respiratory capacities that only manifested under starvation but not replete conditions (Fig. 4C-F, Supplementary Fig. 4-1 C-D). Interestingly, ECAR levels were unaffected by ARMC1 deletion, suggesting that an increase in OXPHOS rather than glycolytic flux is responsible for the elevated metabolic efficiency (Fig. 4G). Mechanistically, re-expressing full-length but not CTD truncated ARMC1 restored OCR levels, thereby suggesting that ARMC1 CTD is responsible for controlling respiratory capacities during starvation.

We next investigated whether this enhanced respiratory capacity was linked to increased FAO using etomoxir, an irreversible inhibitor of Carnitine Palmitoyl-Transferase (CPT1), the enzyme catalyzing the rate-limiting step of mitochondrial FAO ^29,30^. Etomoxir effectively eliminated the starvation-induced increase in respiratory capacity observed in ARMC1^-/-^ cells, but had no impact on oxygen consumption in WT cells (Fig. 4H-J, Supplementary Fig. 4-1 C-D). Consistent with this, etomoxir enhanced lipid droplet accumulation in starved ARMC1^-/-^ but not wild-type cells (Supplementary Fig. 4-1 E). Overall, these results indicate that ARMC1 has a pivotal role in lipid metabolism by regulating mitochondrial FAO and LD storage during starvation.

Next, we stimulate FAO through pharmacological inhibition of Acetyl-CoA Carboxylase 1 and 2 (ACC1, ACC2) using TOFA (5-(tetradecyloxy)-2-furoic acid)^31^. ACC enzymes catalyze the conversion of acetyl-CoA to malonyl-CoA, the rate-limiting step of de novo fatty acid biosynthesis. Because malonyl-CoA serves as a reversible inhibitor of CPT1, the depletion of malonyl-CoA levels via TOFA treatment increases CPT1-dependent fatty acid transport into the mitochondria, thereby promoting FAO ^32^. Treatment with TOFA in WT cells exhibited a dose-dependent increase in maximal respiratory capacity, while ARMC1^−/−^ cells remained entirely insensitive to the drug. Furthermore, even at peak stimulation, the oxygen consumption rate of WT cells never reached the elevated levels observed in ARMC1^−/−^ cells (Fig. 4K). These findings indicate that the loss of ARMC1 fundamentally rewires lipid metabolism, driving mitochondrial FAO to its maximal physiological limit

We next examined whether MTCH2 is essential for the increase in mitochondrial FAO observed in starved, ARMC1 deficient cells. Under starvation, ARMC1/MTCH2 double-knockout cells did not show elevated maximal respiration relative to wild-type or MTCH2 single-knockout cells (Fig. 4L-M, Supplementary Fig 4-1 F), demonstrating that the enhancement of mitochondrial FAO triggered by ARMC1 loss relies strictly on MTCH2. Collectively, our findings suggest that the amphipathic C-terminal domain of ARMC1 modulates mitochondrial FAO and respiratory capacities by controlling access to the hydrophilic groove of MTCH2 to regulate its scramblase activity.

### ARMC1 controls autophagy independent of its C-terminal domain

Mitochondrial ARMC1 partitions into two distinct macromolecular assemblies: one in which ARMC1 associates with MTCH2 and DNAJC11, and a second comprising ARMC1, TMEM11, MTFR, and the mitochondrial motor adaptor MIRO. Functional analysis of the latter pool indicates that the ARMC1–TMEM11–MTFR axis specifically directs retrograde mitochondrial motility¹². Because TMEM11 also acts in BNIP3-dependent mitophagy, this subcomplex may couple ARMC1’s role in mitochondrial FAO to mitochondrial quality control^33^.

First, to evaluate whether partitioning of ARMC1 with TMEM11-containing complex is required for lipid metabolism rewiring under starvation, we measured OCR in TMEM11 knockdown (TMEM11^KD^) and TMEM11^KD^/ARMC1^-/-^ cells (Supplementary Fig. 5-1 A-B). Maximal respiratory capacity was only marginally impaired in TMEM11^KD^/ARMC1^-/-^ cells but unimpacted in TMEM11^KD^ cells, indicating that the alternate ARMC1 receptor TMEM11 is dispensable for the increased FAO observed in starved ARMC1^-/-^ cells (Fig. 5 A-B). Our findings instead demonstrate that ARMC1 modulates mitochondrial FAO primarily through its interaction with the MTCH2-DNAJC11 complex, wherein the ARMC1 CTD complements the cytosol-accessible hydrophilic groove of MTCH2. Corroborating this model, unbiased proteomic profiling revealed the concurrent depletion of crucial TMEM11-associated partners, specifically ARMC1, MTFR1/1L, and MIRO1/2, in TMEM11^KD^ cells, whereas steady-state levels of MTCH2 and DNAJC11 were unaltered (Supplementary Fig. 5C-D). Together, these observations indicate that ARMC1 is distributed between two structurally and functionally segregated macromolecular assemblies to maintain mitochondrial homeostasis: the TMEM11–MTFR–ARMC1–MIRO complex, which governs anterograde mitochondrial positioning^12^, and the MTCH2–DNAJC11–ARMC1 complex, which dictates lipid utilization via mitochondrial FAO.

**Figure 5:**
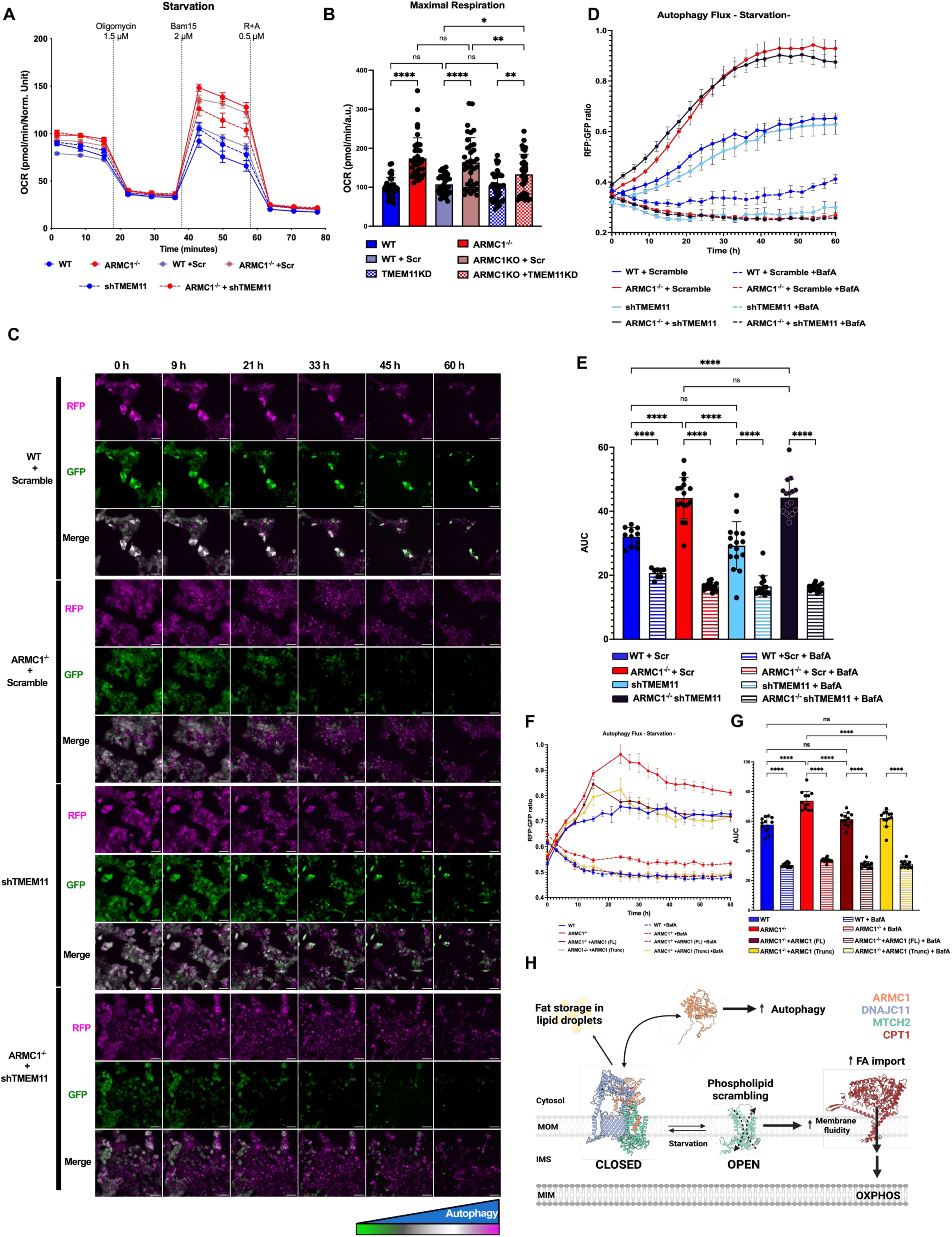
ARMC1 regulates autophagy in a manner independent of its CTD. (A) Oxygen consumption rates during starvation for HCT116 WT cells, ARMC1^-/-^, WT and ARMC1^-/-^ cells expressing a scramble shRNA (Scr), TMEM11 knockdown cells (shTMEM11) and ARMC1^-/-^ + shTMEM11 cells. (B) Maximal Respiratory capacity using the mid-time point after Bam15 injection and normalized against the average of WT. Values are combined from four independent experiments. Error bars represent ±SEM. Statistical significance was assessed by one-way ANOVA followed by Tukey’s test. (C) Representative images of GFP, RFP, and Merge for the detection of starvation-induced autophagy by microscopy for WT + Scramble, ARMC1^-/-^ + Scramble, shTMEM11, and ARMC1^-/-^ + shTMEM11 expressing RFP-LC3-GFP delivered by LNP. Scale bar = 50 µm. (D) Calculation of the RFP:GFP ratio for the autophagy flux mentioned in (C). Error bars mean ±SEM. (E) Area under the curve (AUC) of the cell lines mentioned in (C). Values represent ±SEM. Statistical significance was assessed by two-way ANOVA followed by Tukey’s test. (F) Calculation of the RFP:GFP ratio for the autophagy flux under starvation for WT, ARMC1^-/-^, ARMC1^-/-^ +ARMC1 (FL), ARMC1^-/-^ + ARMC1 (Trunc). Error bars mean ±SEM. (G) Area under the curve (AUC) of the cell lines mentioned in (F). Values represent ±SD. Statistical significance was assessed by two-way ANOVA followed by Tukey test. *P<0.05, **P<0.01, ***P<0.001, ****P<0.0001, and ns = not significant. (H) Schematic summarizing reported interactions by subcellular compartment and cytological response to loss of ARMC1. Created in BioRender. Castonguay, A. (2026) https://BioRender.com/v4hi5kj.

Because prior studies implicated TMEM11 in the regulation of mitophagy by modulating the dimerization of BNIP3 and BNIP3L^33^, a prerequisite for efficient LC3 recruitment and subsequent organelle clearance, we investigated whether autophagic processes were altered in ARMC1^-/-^ cells. Notably, the loss of ARMC1 led to a marked increase in lysosomes and mitochondria overlap, a phenotype fully rescued by reintroducing ARMC1-FL. Expression of ARMC1-Trunc showed similar tendency but not statistical significance (Supplementary Fig. 5-2 A-B). Assessment of autophagic flux during starvation using a GFP-LC3-RFP reporter revealed that ARMC1 depletion significantly accelerates autophagic flux under starvation conditions (Fig. 5C-E, Supplementary 5-3 A). This regulatory effect is independent of the TMEM11-containing complex, as TMEM11 knockdown neither altered baseline autophagic activity nor suppressed the elevated flux in ARMC1^-/-^ cells. Furthermore, expression of both ARMC1-FL and ARMC1-Trunc successfully rescued the elevated autophagic flux, demonstrating that autophagic modulation requires the cytosolic N-terminal domain (NTD) rather than the C-terminal domain (CTD) of ARMC1 (Fig. 5F-G, Supplementary Fig. 5-4A). Together, these data establish a distinct separation of function for ARMC1, wherein the CTD of ARMC1 regulates MTCH2-dependent FAO, while its NTD modulates autophagy independently of TMEM11 (Fig 5H).

## Discussion

The molecular mechanisms coordinating OMM structural plasticity and dynamic substrate utilization during nutrient stress are not fully understood. Here, we establish that the OMM solute carrier homolog MTCH2 forms a stable macromolecular complex with the chaperone protein DNAJC11 and the dual-localized scaffold protein ARMC1. Integrating molecular dynamics simulations with biophysical and genetic analyses, we demonstrate that ARMC1 inserts its amphipathic C-terminal domain (CTD) into the cytosol-accessible, hydrophilic groove of MTCH2. This precise insertion blocks the inherent lipid scramblase and protein insertase activity of MTCH2 by effectively acting as its “missing” sixth transmembrane helix. Because the endogenous copy number of MTCH2 is greater than its interacting partners ^12, 34^, our findings point to an equilibrium shifting between free MTCH2 monomers and structured tripartite complexes, functioning as a tunable rheostat that responds directly to metabolic and environmental cues.

This structural equilibrium manifests as a functional metabolic switch during nutrient deprivation, governing the balance between fat storage in LDs and mitochondrial FAO. Deletion of MTCH2 induces severe oxidative phosphorylation defects, mitochondrial fragmentation, and the downstream accumulation of LDs. Conversely, ablation of ARMC1 or selective truncation of its CTD completely relieves the steric inhibition on the MTCH2 hydrophilic groove, driving an increase in MTCH2-dependent oxygen consumption and depletion of cellular LD pools. Through pharmacological epistasis, we reveal that ARMC1^-/-^ cells achieve a functional ceiling in respiratory capacity, indicating that the absence of ARMC1 locks mitochondrial FAO at its maximal physiological rate in a MTCH2-dependent manner. Previous work has proposed that MTCH2 influences FAO via direct interaction with CPT1, the rate-limiting gatekeeper for mitochondrial FAO ^35^. Inspection of crosslinking mass spectrometry public data and our proteomic profiling argue against a direct MTCH2-CPT1 interaction, suggesting instead that the lipid scramblase activity of uncomplexed free MTCH2 rewires mitochondrial membrane lipid composition–potentially through cardiolipin precursor transport–to intrinsically sensitize or optimize the mitochondrial FAO machinery.

Importantly, our data highlight a remarkable spatial and functional separation of ARMC1 roles mediated by its dual localization. While a portion of the mitochondrial pool of ARMC1 interacts with MTCH2 via its CTD to suppress starvation-induced lipid burning, the ARM and HMAL domains facilitate formation with a distinct macromolecular assembly featuring TMEM11, MTFR1/2/1L, and MIRO1/2 ^12^. Genetic targeting of TMEM11 co-depletes this specific trafficking complex but fails to impact starvation-induced respiration, demonstrating that the TMEM11 complex is dispensable for metabolic substrate switching. Intriguingly, ARMC1 deficiency also triggers a dramatic increase in global autophagic flux and increases mitochondria-lysosome proximity under starvation stress. Because this autophagic acceleration occurs independently of the TMEM11 network and is fully rescued by a CTD-truncated mutant, it is driven strictly by the cytosolic pool of ARMC1. Collectively, this work uncovers a highly coordinated regulatory framework where ARMC1 partitions between distinct sub-cellular pools to simultaneously restrain lipid oxidation. At the mitochondrial surface ARMC1 regulates FAO via its CTD while cytosolic ARMC1 fine-tune autophagic flux via its N-terminal ARM and HMAL domains, establishing a pathway for metabolic flexibility during nutrient deprivation.

## Materials and methods

### Cell culture and reagents

HCT116 parental cell line was obtained from ATCC (CCL-247). MTCH2 KO cell line was generated previously ^10^. All cell lines were cultured in McCoy’s 5A Modified medium (Gibco, 16600082) supplemented with 10% FBS (Gibco, 10-437-028) and maintained in 5% CO_2_ at 37°C. All cell lines were verified for mycoplasma testing using MycoAlert Mycoplasma Detection Kit (Lonza, LT07-318). Starvation was induced by culturing the cells for five hours in Hank’s Balanced Salt Solution, HBSS (Gibco, 24020117).

### Generation of ARMC1 knockout cell lines

To generate ARMC1 knockout (KO) models, a synthetic single-guide RNA (sgRNA) was designed using the Benchling CRISPR Guide RNA Design tool to target exon 2 of the *ARMC1* locus (Target sequence: 5’-AAACAGAAGAGCCATCGTCC-3’). Custom sgRNAs were synthesized by Invitrogen (Thermo Fisher Scientific). Knockout lines were generated in both parental HCT-116 cells and a previously established *MTCH2* KO HCT-116 background to produce single and double KO lines, respectively.

To facilitate gene editing, ribonucleoprotein (RNP) complexes were formed by incubating 105 pmol of Alt-R S.p. Cas9 Nuclease V3 (IDT, 10000735) with 120 pmol of sgRNA (1:1.15 molar ratio) in PBS for 20 minutes at room temperature. The RNP complexes, supplemented with 100 pmol of Alt-R Cas9 Electroporation Enhancer (IDT, 1075916), were delivered into 2 x 10^5^ HCT-116 cells via electroporation using the Lonza 4D-Nucleofector System (SE Cell Line Nucleocuvette Kit, V4XC-1032) under program EN-113.

Following expansion, the editing efficiency of the bulk cell pool was validated. Genomic DNA was extracted, and the target locus was amplified by PCR (primers: forward 5’- CCTCGGGCACTTGTTTTAGGGC -3’ and reverse 5’- CTACTTGGGAGGCTGAGGTCGG -3’). The resulting amplicons were Sanger sequenced (Quintara Biosciences), and the knockout efficiency was quantified using Synthego’s Inference of CRISPR Edits (ICE) tool.

Upon confirmation of pool editing, single cells were isolated via fluorescence-activated cell sorting (FACS). Cells were gated based on forward and side scatter (FSC/SSC) profiles to ensure the selection of a viable, single-cell population and were sorted into 96-well plates. Following clonal expansion, the complete loss of ARMC1 protein expression was confirmed by Western blot analysis to ensure successful gene editing.

### Lentivirus preparation and cell line generation

Recombinant genes were synthesized and cloned (GenScript, Twist Bioscience) into pLVX-eF1 vector in place of mCherry fluorescent reporter and sequence-verified via nanopore whole plasmid sequencing (PlasmidSaurus). Recombinant pLVX cargo vectors were co-transfected into Lenti-X cells with psPAX2 and pMD2.G packaging vectors at a ratio of 4:3:1 (w/w) using TransIT-VirusGEN Transfection Reagent (Mirus Bio, MIR 6705) according to manufacturer’s instructions. At 48 h post-transfection, T25 flask contents were transferred to 50 mL conical tubes and cells pelleted by centrifugation at 300 x g for 5 min at room temperature. Resulting contents were filtered through 0.45 μm PVDF filter (Sigma, SE1M003M00) and lentiviral particles precipitated with 3x volume of Lenti-X Concentrator overnight at 4°C. Following overnight precipitation, lentiviral particles were collected by centrifugation at 4,000 x g for 45 min at 4°C and the resulting lentiviral pellet resuspended in PBS, aliquoted, and stored at -80°C. Concentration of lentiviral particles was estimated by functional titer and recipient KO cell lines infected with an MOI of 0.1. Stably transduced cell lines were selected by treatment with puromycin (2 μg/mL) for > 1 week with at least 3x media exchanges. Expression of transgenes were validated by immunoblot compared to un-transduced controls.

### SDS-PAGE and immunoblotting

Cells were passaged and treated as described above unless otherwise noted. For lysate collection, cell cultures were washed twice with 1 plate volume of ice-cold PBS (Gibco, 10010-023) then lysed on plate with 1/20 volume of 1x RIPA (EMD Millipore, 20-188) supplemented with 1x HALT protease inhibitor cocktail (Thermo Scientific, 78445) and incubated on ice for 15 min. Cell debris was scraped, further homogenized using water bath sonicator (Qsonica Q800R3 with 5 s pulse, 50% amplitude, 1 min total time), and protein concentration measured by BCA assay (Thermo Scientific, 23225) or Pierce 660nm Protein Assay (Thermo Scientific, 22660). Cell lysates were normalized to 1-2 mg/mL, supplemented with 4x Laemmli buffer + BME (Sigma-Aldrich, M6250-100ML), boiled for 5 min, then re-equilibrated to 4°C on ice. Unless otherwise specified, 10 ug of protein was loaded per lane in precast SDS-PAGE gels (Bio-Rad Criterion TGX stain-free); percent acrylamide of gel varied depending on molecular weight of target protein. Protein samples were resolved at 100 V (voltage constant, room temperature) until exit of dye front from gel. Gels were washed 2x with distilled water to remove buffer salts, then even loading and sample quality assessed by stain-free signal acquired by Bio-Rad ChemiDoc gel imager.

Resolved protein species were then transferred to 0.2 micron PVDF membrane by semidry method using Bio-Rad Trans-Blot Turbo Transfer System (Bio-Rad, 1704150) and kit (Bio-Rad, 1704273) according to manufacturer’s instructions. Following transfer, membranes were washed 3x with TBS + 0.1% tween-20, blocked for 5-60 min in EveryBlot Blocking Buffer (Bio-Rad, 12010020), and incubated with primary antibody diluted in blocking buffer overnight at 4°C with gentle agitation. Dilution of primary antibody was chosen based on manufacturer’s criteria (see Supplementary Table 1). Four washes of 5 min each were performed at room temperature with TBS + 0.1% tween-20 following primary antibody incubation and membranes transferred to secondary antibody solution in blocking buffer. AlexaFluor-conjugated goat-anti-mouse IgG and goat-anti-rabbit IgG were used with different attached fluorophore but each to 1:5,000 dilution. Following 30-60 min incubation with secondary antibody at room temperature with gentle agitation, 4 additional washes were performed (same wash buffer and conditions). Membranes were scanned using Azure Sapphire gel imager and immunoblot images processed with ImageJ.

### Fluorescence Microscopy

Cells were seeded in 24-well glass-bottom plates with #1.5 cover glass (0.170±0.005mm) in McCoy’s 5A Modified medium (Gibco, 16600082) supplemented with 10% FBS (Gibco, 10-437-028) and maintained in 5% CO_2_ at 37°C for 48 h. Cells were stained with Mitotracker Deep Red (Invitrogen, M22426) at 100 nM for 20 minutes in 5% CO_2_ at 37°C in McCoy’s 5A Modified medium supplemented with 10% FBS, washed twice with DPBS (Gibco, 14190144), followed by replenishment of fresh media or Hank’s Balanced Salt Solution, HBSS (Gibco, 24020117) for starvation induction. After 5 hours of starvation, cells were fixed for 20 minutes in 4% PFA pre-warmed, washed three times with DPBS (Gibco, 14190144), and maintained at 4°C until imaging acquisition or immunofluorescence staining. Bodipy 493/503 (ThermoFisher, D3922) and Hoechst (ThermoFisher, 62249) staining was performed separately in fixed cells for 20 minutes each at 1 µM and 2 µM, respectively, in DPBS, followed by three washes in DPBS.

For immunofluorescence staining, fixed cells were permeabilized with 0.1% Triton X-100 (Millipore Sigma, 9036-19-5) for 20 minutes at room temperature (RT), washed three times with DPBS, blocked with 5% of bovine serum albumin (BSA) in DPBS (Sigma Millipore, 9048-46-8), and immunolabeled overnight at 4°C in 1% of BSA in DPBS with mouse anti-Flag or rabbit anti-ALFA, as indicated. Samples were washed three times with DPBS, incubated with goat anti-mouse Alexa Fluor 567 in 1% of BSA in DPBS for 1 hour at RT, washed three times with DPBS and maintained at 4°C until imaging acquisition.

With exception of supplementary Figure 2-2, all epi-fluorescence images were collected with a Zeiss LSM 980 microscope utilizing an Airyscan 2 detector. An 63x/1.4 Oil DIC Plan-Apochromat objective (Zeiss part number 420782-9900-799) was used for all imaging. High-resolution Z-stacks were collected with a 0.3 µm step size using the Airyscan 2 detector in Super-Resolution, 8-line multiplexing mode (SR-8Y; spot size equivalent to 8 airyscan channels along y sweep axis; nominal resolution 120 × 160 × 450 nm**3). All Airyscan images were 3D processed in ZEN Blue software using the standard Airyscan processing algorithm to reconstruct the super-resolution image.

Images in supplementary Figure 2-2 were captured using a Zeiss Celldiscover 7 (CD7) high content microscope operating in widefield epi-fluorescence mode from 24 well plates. These plates were seeded and prepared for IF microscopy as described above but with mouse anti-ALFA and rabbit anti-TOM20 antibodies. To capture an entire cell volume that is outside any one focal plane, 16 um deep focal series (58 slices with 0.280 um step size) comprised of 0.276 um × 0.276 um images (6×6 fused tiles) of each well were collected using a Plan-Apochromat 50×/1.2 water immersion objective employing the 0.5x tube lens on the CD7. Z-series Images acquired with the CD7 were scaled and cropped for display using ZEN Blue software version 3.13.

### Imaging Analysis

For Lipid Droplet quantification, images were imported to Arivis Pro 4.4.0 for analysis in 3D to capture particle size across all the Z-stacks. Mitochondria (MitoTracker deep red 644/665 nm) were denoised using the mean intensity method and segmented using the Otsu method. In combination with nuclei (Hoechst 367/497 nm), both channels were used for cell segmentation using a Cellpose model imported into Arivis Pro. Lipid droplets were background-corrected, morphology-filtered to preserve the brightest objects, and were segmented using the Blob Finder tool on Arivis, utilizing a diameter filter of 300 nm. Objects were exported, excluding edges, and filtered by Surface Area (Voxels).

For colocalization analysis between mitochondria and Alfa-ARMC1-FL or ARMC1-Trunc we used Fiji 2 (Schindelin et al 2012). Cells were segmented manually to calculate Pearson’s coefficient by using Coloc 2 plugin. For overlap area between mitochondria and LAMP1, cells were manually segmented and thresholded followed by mask selection. Area between the two different masks was calculated using the “AND” function on Fiji that measures the intersection of distinct channels.

### Proteinase K protection assay

Proteinase K protection assay for assessing MTCH2 topology was performed as described previously ^36^. Briefly, mitochondria were enriched from stable cell lines expressing MTCH2 with either N- or C-terminal 3xFlag tag by differential centrifugation. Aliquots of enriched mitochondrial membranes (50 ug each) were resuspended in homogenization buffer plus or minus mild detergent (1% Triton-x100) or in hypotonic buffer to induce swelling. Resuspended membranes were then treated with proteinase K. Equal amounts of protein resolved by SDS-PAGE and MTCH2 detected via immunoblot for 3xFlag epitope tag. Topology was assigned based on comparison with mitochondrial subcompartment markers TOM70 (MOM), IMMT/MIC60 (MIM), and ATP5A1 (matrix).

### Cysteine accessibility assay

Mitochondria were isolated from HCT116 cells as previously described for U2OS cells ^37^ but with 25-30 strokes of tight-fitting Dounce homogenizer. Following mitochondrial enrichment, protein concentration was quickly measured by BCA assay (Thermo Scientific, 23225), normalized to 1 mg/mL in homogenization buffer (10 mM HEPES, 1 mM EDTA, 210 mM mannitol, 70 mM sucrose, pH 7.4, 1x protease inhibitor cocktail), and 25 ug aliquots maintained on ice. Disruption of mitochondria was performed by addition of SDS to 1% final. PEG-maleimide moieties of increasing molecular weight were used for modification of free thiols at 2,000 Da (Nanocs, PG1-ML-2k), 5,000 Da, (Nanocs, PG1-ML-5k-1), and 10,000 Da (Nanocs, PG1-ML-10k). mPEG-maleimide stock solutions were prepared to 20 mM in homogenization buffer, equilibrated to 4°C, and spiked into mitochondrial aliquots at 2 mM final concentration. At each timepoint within the time course (0 – 60 minute), PEGylation reaction was quenched with 25 mM dithiothreitol (DTT). Note, in contrast to other methods no reducing agents were used prior to incubation with mPEG-maleimide. Quenched samples were prepared and resolved by SDS-PAGE as described below but without supplementation of 2-mercaptoethanol.

### Split luciferase assay

ARMC1, DNAJC11, and MTCH2 interaction was evaluated *in cellulo* using the NanoBiT PPI MCS Starter System (Promega, N2014) based on manufacturer’s instructions but with the following modifications. Briefly, ARMC1, DNAJC11, and MTCH2 were tagged with either the LgBiT and SmBiT split luciferase components ^24^ with position of either appendage guided by AF2 modeling. Recombinant genes encoding the chimeric proteins were codon optimized, synthesized, cloned in pBiT1.1-N-TK-LgBiT replacing LgBiT segment (Twist), and resulting vectors validated by nanopore whole plasmid sequencing (PlasmidSaurus). Chimeric reporter constructs were delivered to WT HCT116 cells using Lipofectamine 3000 Transfection Reagent (Thermo Scientific, L30000015) according to manufacturer’s instructions and total lysates resolved by SDS-PAGE on BIORAD Criterion 4-15% gradient polyacrylamide gels (20 ug/lane). Expression of recombinant proteins was validated by immunoblot with antibodies specific to either the endogenous protein or anti-LgBiT antibody.

Luciferase activity was tested for all binary combinations of recombinant SmBiT-LgBiT pairs. In short, 1e4 WT HCT116 cells were seeded per well in 96-well white-walled, white-bottom plates (Costar, 3917). Twenty-four hours following seeding, constructs were co-delivered (50 ng each, 100 ng vector total/well) via Lipofectamine 3000 pairs and incubated overnight. At 48 h post cell-seeding, media containing transfection reagent was removed, cells were washed, and shifted to either replete or starved conditions as described above. After 5 h of media treatment, cells were washed 3x with phenol free HBSS (Fisher Scientific, 14-025-134), left in final wash, and Luciferase mix added. Luciferase activity was measured using BMG ClarioSTAR microplate reader with incubator module under the following instrument conditions: 2×2 matrix scan of each well (6 mm diameter), enhanced dynamic range mode, total luminescence (no filter) with crosstalk aperture. Measurements were collected at 37°C and 5% CO2 with 15 s double orbital shaking (300 rpm) performed prior to each reading. Samples were prepared in biological triplicate. Average luminescence signal for each interacting pair was plotted and statistical analysis (one-way ANOVA) performed by GraphPad Prism 10 software.

### Bacterial (co)expression and purification

Recombinant protein expression and purification was performed through Viva Biotech Ltd. (Shanghai). Briefly, full length ARMC1 (1-282) or truncated (1-235) was cloned into pET15b with N-terminal 10xHis-StrepII-3C tag, verified by whole plasmid sequencing (PlasmidSaurus), and transformed into BL21 (DE3) cells for protein expression. Recombinant protein was induced by treatment with 1 mM IPTG overnight at 16°C (100 rpm) and monitored by Coomassie stain of whole cell lysates resolved by SDS-PAGE. Expression of truncated but not full length ARMC1 was readily detectable; therefore, expression was also attempted in BL21 (DE3) LOBSTR, BL21 (DE3) Codon Plus, and Rosetta (DE3) cells lines unsuccessfully. Full length ARMC1 with 10xHis-StrepII-3C- tag was cloned into the MCS-1 polylinker of pETDuet-1 for co-expression with DNAJC11 C-terminus (411-559) at MCS-2 and verified by whole plasmid sequencing (PlasmidSaurus). Recombinant protein co-expression was induced and measured as with pET15b constructs. Bacterial cell pellets containing recombinant protein(s) were resuspended in gentle lysis buffer (20 mM Tris HCl pH 7.5, 300 mM NaCl, 5 mM MgCl_2_, 1 mM DTT, 10% glycerol, 1x Roche protease inhibitor cocktail) at 5 mL/g wet cell mass, then spiked with 50 U benzonase/g wet cell mass, and lysed by ATS high pressure homogenization (1x 400 Bar, 2x 800 Bar). Recombinant protein(s) were then tandem (co)purified via Strep and Ni columns. The resulting purified protein was then fractionated by size exclusion chromatography with S75 column and high purity fractions pooled for subsequent analysis by SDS-PAGE and LC-MS for intact mass determination.

### Size Exclusion Chromatography and Multi-Angle Light Scattering (SEC-MALS)

40 µl of 2 mg/ml ARMC1 (1-282) + DNAJC11 (411-559) was injected onto a Superdex 200 10/300 gl column (Cytiva) equilibrated in 20 mM Tris-HCl, pH 7.5, 150 mM NaCl, 1 mM DTT. The column was coupled to a static 18-angle light-scattering detector (DAWN, Wyatt Technology) and a refractive index detector (G7162A, Agilent). Data were collected at a flow rate of 0.3 ml/min. Data analysis was performed using Astra VII. 40 µl 2 mg/ml monomeric BSA (2.0 mg/ml) (Thermo Scientific) was used for quality control and as an isotropic scatterer benchmark.

### Yeast two-hybrid

Interaction between DNAJC11 and ARMC1 was evaluated using the MatchMaker Gold Yeast two-hybrid system (TaKaRa, 630489) according to the manufacturer’s instructions. Briefly, identified segments of DNAJC11 and ARMC1 were selected, yeast codon optimized versions synthesized (GenScript), and cloned into pGADT7-AD and pGBKT7-53 vectors, respectively. Recombinant vectors were verified by nanopore sequencing (PlasmidSaurus), and introduced into yeast host by heat shock method. Recombinant bait (pGBKT7) and prey (pGADT7) containing strains were selected for by the nutritional markers -Trp and - Leu, respectively. Interaction between N- or C-terminal segments of DNAJC11 with ARMC1 was assessed via simple spot assay on selective media containing Aureobasidin A for viability and X-alpha-gal for blue coloration, by cells expressing AUR1-C and MEL1 genes, respectively. A single positively interacting DNAJC11-CTD segment was chosen for further mutagenesis (DNAJC11-Arg413-CTD). Positions of amino acid substitutions in DNAJC11 or ARMC1 were guided by AF2 multimer modeling and the corresponding codon optimized mutant sequences generated through de novo gene synthesis. WT and variant protein combinations of DNAJC11 and ARMC1 were evaluated via MatchMaker Gold system as described above and interaction assessed via 5-fold dilution series. Briefly, 1 day old stationary phase cultures were pelleted, washed 2x with 150 mM sterile NaCl, and cultures normalized to an OD600 of 1.0 in 150 mM NaCl. Five-fold serial dilutions of each combination and kit provided controls were spotted on selective media or YPD and incubated for 2 days in a humidified chamber at 30°C after which plates were photographed using Singer PIXL plate imager.

### Affinity purification

WT and 3x-Flag-MTCH2 expressing HCT116 cells were cultured under either replete or starved conditions (n = 4) and crosslinked *in cellulo* as described previously ^37^ with the following modifications. After quenching, DSP-crosslinked cells were washed 2x with ice cold PBS (Gibco, 10010-023), scraped in 1 mL cold PBS and pelleted by centrifugation at 2,000 x g for 5 min (4°C), then supernatant removed by aspiration. The resulting crosslinked cell pellet was resuspended to 5 ml/g of wet cell mass in ice cold 1x RIPA (EMD Millipore, 20-188) supplemented with 0.1% SDS and 1x protease inhibitor cocktail (Thermo Scientific, 78445), then solubilized by end over end rotation for 30 min at 4°C. Insoluble material was pelleted by at 20,000 x g for 10 min at 4°C, the supernatant transferred to new tubes and flash frozen in liquid nitrogen. Protein concentration was measured by BCA assay (Thermo Scientific, 23225) and normalized to 1 mg/mL in lysis buffer. Affinity capture of 3xFlag-MTCH2 was facilitated by anti-Flag nanobody-conjugated magnetic beads (Engibody, AT1770) according to manufacturer’s protocol with the following modifications. Total amount of bead slurry was extrapolated for number of samples, washed, and prepared according to manufacturer’s instructions). Twenty-five uL of washed beads were added per mg of crosslinked lysate and incubated for 4 hr at 4°C with end over end rotation. MTCH2-interacting proteins were captured by magnetization of beads for 1 min and washed 3x with 0.5 mL lysis buffer followed by an additional 3 washes with 0.5 mL of 0.1 M Tris-HCl, pH 8.5. Washed bead-protein mixtures were resuspended in 0.5 mL of 0.1 M Tris-HCl, pH 8.5, and reduced/alkylated by addition of 10 mM TCEP-HCl pH ∼7 (Sigma-Aldrich, 20491) and 40 mM 2-chloroacetimide (thermo scientific, A15238.30) at 95°C for 5 min then cooled to room temperature. Co-captured proteins were digested on-bead by addition of 25 ul of 0.1M Tris pH 8.5 buffer containing 15 ug/ml of LysC and 15 ug/ml of Trypsin (for a final trypsin and LysC concentration of 5 ug/ml), and incubated overnight in thermomixer at 37 °C and 1,000 rpm. Following overnight incubation, condensation was pelleted by quick spin, then acidified with trifluoroacetic acid to 1% final and beads removed by magnetization. Supernatant containing digested peptides was transferred to new tubes and stored at -80°C.

### Lysate preparation and protein digestion for whole cell proteomics

Cell lysates (n = 5) were prepared as for immunoblotting analyses until the addition of Laemmli buffer then 10 mM TCEP-HCl pH ∼7.0 (Sigma-Aldrich, 20491) and 40 mM 2-chloroacetamide (Thermo Scientific, A15238.30) added for one-step reduction/alkylation reaction at 95°C for 5 min. Protein was precipitated with 4 volumes of ice cold acetone overnight at -20°C. Following overnight incubation, precipitated protein was collected by centrifugation at 20,000 x g for 30 min, supernatant removed, pellet dried at room temperature for 10-15 min, then resuspended to 5 μg/μL in 8 M urea in 100 mM Tris-HCl pH 8.1. Total protein was sequentially digested by LysC (Wako, 125-05061) with 1:50 ratio LysC:protein overnight at 25°C, 1,000 rpm, then diluted 1:5 with 100 mM Tris-HCl pH 8.1, trypsin (Sigma-Aldrich, T6567-5X20UG) added at 1:50 ratio and incubated for 5 hrs at 25°C and 1,000 rpm in thermomixer. Trypsin was quenched by acidification with trifluoroacetic acid to 1% final concentration and stored at -80°C.

### Proteomics sample preprocessing and LC-MS/MS analysis

Samples for proteomics analyses were preprocessed and run in NanoFlow as described previously ^38^ with exception of MTCH2^-/-^ whole cell proteomics which was run in Microflow mode but with identical pre-processing and similar LC gradient. Briefly, filtration plates containing 0.45 um membranes (Millipore Sigma, MSRLN04) were activated with 400 uL of methanol and then equilibrated with 400 uL methanol with 1% TFA (volumes are per well). Following filtration plate equilibration, 250 uL of peptide mixture was applied to each well and the volume brought up to 500 uL with 1% TFA and clarified by centrifugation at 1,000 x g for 1 min. The resulting flow through was then subjected to automated Solid-Phase Extraction (SPE) performed on an Agilent AssayMap Bravo platform equipped with 5 ul C18 reverse-phase cartridges. SPE cartridges were activated with 50% acetonitrile (ACN) and 0.1% TFA, then equilibrated with 0.1% TFA. The filtered samples were then loaded, bound peptides washed with 0.1% TFA, and subsequently eluted in 70% ACN and 0.1% TFA. The desalted peptide eluates were concentrated under vacuum and resuspended in 3% ACN and 0.1% formic acid for a final peptide concentration of 2.5 g/L. Peptides were analyzed by LC-MS/MS with Vanquish Neo UHPLC system (Thermo Fisher Scientific) coupled to an Orbitrap Ascend Tribrid mass spectrometer (Thermo Fisher Scientific). Chromatographic separation was achieved on a Waters CSH C18 column (1.7 μm, 1 × 150 mm) maintained at 55°C. The mobile phase system consisted of solvent A (0.1% formic acid in water) and solvent B (80% ACN, 0.1% formic acid in water). Peptides were separated using a two-step linear gradient: 3–35% B over 56 min, followed by 35–45% B over 4 min, at a constant flow rate of 50 ul/min prior to ionization with MS settings described as in ^38^. All reagents and solvents were LC-MS-grade.

The Orbitrap Ascend mass spectrometer was operated in the positive mode using a heated electrospray ionization source (H-ESI), with a spray voltage of 3,500 V, sheath gas at 32°C, sweep gas at 0°C, ion transfer tube temperature at 325°C, and vaporizer temp at 125°C. Full MS scans were acquired in the Orbitrap at a resolution of 120,000, normalized automatic gain control (AGC) target at 100%, maximum inject time (IT) of 251 ms, RF-lens setting of 60%, and a full scan range of 350 — 1,050 m/z.

Data-independent acquisition (DIA) scans were acquired with higher-energy collisional dissociation (31% normalized collision energy), at an Orbitrap resolution of 15,000, over a scan range of 200 - 1,800 m/z. Seventy overlapping 9.5 m/z windows covering the mass range of 384.5 – 1015 m/z were used, with an AGC target of 1,000%, and a maximum IT of 27 ms.

### Raw Data Search using DIA-NN v1.8

DIA raw files were searched with DIA-NN v1.8 in library-based mode against an in silico–predicted spectral library, which was reannotated during the run against the Homo sapiens UniProt reference proteome (including isoforms) combined with a common-contaminants database. In silico tryptic digestion (cleavage after K/R) allowed a maximum of 1 missed cleavage with N-terminal methionine excision enabled; peptides were restricted to 7–30 residues, precursor m/z to 300–1800, and precursor charge to 1–4.

Cysteine carbamidomethylation (+57.0215 Da) was applied as a fixed modification on cysteine. Two variable modifications were defined — methionine oxidation (UniMod:35, +15.9949 Da) and protein N-terminal acetylation (UniMod:1, +42.0106 Da) — with at most one variable modification permitted per peptide. N-terminal acetylation was further designated a monitored modification.

Mass accuracy was fixed at 10 ppm for MS2 (fragment) and 20 ppm for MS1 (precursor) ions. A two-pass search was used: an empirical spectral library was first generated from the DIA data and all runs were then re-searched against it. Identifications were filtered to 1% FDR at both the precursor and protein-group levels, with protein inference based on proteotypic peptides only.

### Data Analysis of Searched Results

MS intensities filtered by 1% FDR from PG.MaxLFQ in DIA-NN’s report output were log2 transformed and imputed before analysis. To filter for reliably detected and quantified protein groups, those that were not quantified in all replicates in at least one sample group were removed from the dataset.

With the filtered dataset, missing quantifications were assumed to be below the detection threshold of the MS and imputed. Missing values were randomly sampled from an imputation distribution derived from the intensities of each sample replicate. If the sample distribution has a mean of u and a standard deviation of v, the imputation distribution was defined with a mean of u - 1.8v and a standard deviation of 0.03v.

For the analysis of the differential abundance of proteins, the R limma package (v.3.66.0 / Bioconductor 3.21) was used to calculate the log2 fold change, p-values, and Benjamini-Hochberg adjusted q-values.

### Redundancy analysis

To quantify the proportion of proteomic variance explained by genetic constructs and environmental conditions, we performed Redundancy Analysis (RDA) using the vegan package, version 2.7-1 (https://CRAN.R-project.org/package=vegan). Protein abundance data were first z-score scaled and centered. For the first dataset, initial screening indicated that the orientation of the protein tag (C-terminal vs. N-terminal) did not significantly contribute to proteomic variance. Consequently, the genetic component was modeled as a numerical variable representing the presence or absence of the MTCH2 locus, and the RDA was constrained by the formula: Abundance ∼ MTCH2 *Media. For the second dataset, where different rescue constructs exhibited distinct proteomic phenotypes, the genetic component was modeled as a categorical factor. This analysis was constrained by the formula: Abundance ∼ ARMC1 Construct Category * Media, allowing for the evaluation of interaction effects between specific alleles and environmental treatments. The statistical significance of the constraints and individual axes was determined using permutation tests (n = 999). The resulting RDA coordinates were used to identify protein transition vectors and pathways associated with specific phenotypic clusters.

### Structural model building & molecular dynamics simulations

AlphaFold2 monomer models of MTCH2 and ARMC1 were obtained from the AlphaFold structural database^39^, and AlphaFold2-multimer models were generated with the Openfold pipeline^40,41^ using default parameters for MSA construction and multimer inference. Additional structural models were generated using the Boltz-1/2 pipelines with default inference parameters^42,43,44^.

CG-MD simulations were parameterized using the first generation Martini3 force-field for protein, lipids, water, and ions^45^. Martini3 protein topologies were prepared from AF2 models for MTCH2 and the MTCH2:ARMC1-Ct complex (containing ARMC1 residues 259-282) first using the Martinize2 application^46^ then adding additional custom elastic bonds that were empirically tuned to match the hydrophilic groove width distributions and beta hairpin structures observed in preliminary all-atom CHARMM36 simulations for each protein or complex, as these features were not adequately maintained by the default Martinize2 parameterization in test simulations. Initial coordinates for simulation systems were prepared by manually orienting 6 copies of each protein into a pre-equilibrated bilayer model composed of a 42.5:42.5:5:5:5 mixture of POPC:POPE:POPI:POPA:LPA, with 3,000 initial lipids, after which lipids clashing with inserted proteins were deleted. The systems were solvated with standard Martini3 water beads plus ∼1,000 Na+ and ∼600 Cl- ions within ∼290×290×150 Å simulation cells. CG-MD simulations were run using GROMACS 2023.2^47^. Each system was initially minimized for 5,000 steps using a steepest descent algorithm, followed by initializing particle velocities from a Maxwell-Boltzmann distribution at 310 K and 5 ns of equilibration in the NPT ensemble with 50 kJ/mol*nm^2^ harmonic restraints on all protein backbone beads. Next, harmonic restraints were removed for a longer 500 ns equilibration phase, after which production simulation trajectories of 10-20 μs were initiated. Simulations used a 20 fs timestep using a leap-frog integrator, a temperature of 310 K enforced by a stochastic velocity rescaling thermostat with a time constant of 1 ps, and a pressure of 1 atm enforced by a stochastic cell rescaling barostat with a time constant of 20 ps. Short range nonbonded interactions were cut off at 11 Å, long range electrostatics were treated with the reaction field method, and force-field specified bonds were constrained with the LINCS algorithm.

AA-MD simulations used the CHARMM36 force-fields for proteins^48^ and lipids^49^, and the CHARMM-TIP3P water model with standard compatible ion parameters. Protein models were prepared from Boltz-2 models of MTCH2 WT, MTCH2 K25E, and MTCH2:ARMC1-Ct complex by first adding N-terminal acetyl caps to each chain in PyMOL, and then using the GROMACS pdb2gmx application to generate topology files. Initial coordinates for most simulation systems were prepared by manually orienting 3 copies of each protein or complex into two pre-equilibrated bilayer models composed of 640:160 mixtures of POPC:POPA, the first with an equal distribution of lipids between leaflets (symmetric), and the second with all POPA in the cytosolic leaflet (asymmetric), after which lipids clashing with inserted proteins were deleted. Systems were then solvated with TIP3P water and neutralizing K+ ions within ∼160×160×100 Å simulation cells. Hydrogen-mass-repartitioning of 2 amu was applied to all protein and lipid hydrogen atoms. AA-MD simulations were run with GROMACS 2025.3 or 2022.2 (preliminary). Note that shorter, preliminary AA-MD simulations were prepared with one copy of the MTCH2 AF2 model in a smaller bilayer of either pure POPC, or a phospholipid mix designed to mimic the bulk composition of the OMM - these simulations informed later CG and AA models but were not analyzed in detail; settings closely matched later runs. Each system was initially minimized using a steepest descent algorithm until the maximum force reached 1,000 kJ/mol/nm. Velocities were then initialized from a Maxwell-Boltzmann distribution at 150 K, and systems were heated in the NPT ensemble to 325 K over 80 ps while maintaining 1,000 kJ/mol*nm^2^ harmonic restraints on all protein heavy atoms. Restraints were stepped down over the course of 2.5 ns of equilibration in the NPT ensemble, until a final phase of 10 ns with restraints of 30 kJ/mol*nm^2^ on only protein Cα atoms, after which production trajectories with no restraints were initiated. Simulations used a 4 fs timestep using a leap-frog integrator, a temperature of 325 K enforced by a stochastic velocity rescaling thermostat with a time constant of 1 ps, and a pressure of 1 atm enforced by a stochastic cell rescaling barostat with a time constant of 5 ps. Short range nonbonded interactions were cut off at 12 Å, Lennard-Jones interactions were smoothly switched off in the range 10-12 Å, long range electrostatics were treated with the particle mesh Ewald method, and non-water bonds to hydrogen were constrained with the LINCS algorithm. Analysis of simulation trajectories (RMSD, RMSF, lipid headgroup density & headgroup translocation tracking) was performed with custom scripts built on the MDAnalysis python library, version 2.9.0 ^50^.

### Seahorse assay

Respiratory capacity was evaluated using an XF Pro Analyzer (Agilent). XF Pro Cell Culture Microplates (Agilent, 103794-100) were first coated with 30 µL/well of 0.1 mg/mL Poly-D-Lysine (Gibco, A3890401) for 40 min at room temperature (RT), washed twice with sterile PBS (Gibco, 10010-023), and dried for 1 h. HCT116 cells, cultured in McCoy’s 5A Modified medium with 10% FBS (replete medium), were detached using TrypLE Express (Gibco, 12604-013) and counted via ViCell analyzer (Beckman). Cells were then seeded at 20,000 cells/well in 50 µL, allowed to settle at RT for 30 min before the addition of 150 µL replete medium, and then cultured for at least 24 h at 37°C in 5% CO₂. Prior to the assay, XF Pro sensor cartridges (Agilent, 103793-100) were hydrated overnight in Calibrant solution at 37°C in a non-CO₂ incubator. On the day of the assay, cells were first subjected to a 4 h pre-incubation at 37°C in 5% CO₂ in either replete medium or HBSS. Following this, the medium was replaced with the appropriate assay medium for 1 h of equilibration in a non-CO₂ 37°C incubator. Cells in replete condition received Seahorse XF DMEM (Agilent, 103575-100) supplemented with 10 mM glucose, 1 mM pyruvate, and 2 mM glutamine (Agilent, 103577-100, 103578-100, 1035789-100, respectively) while HBSS pre-treated cells received the same medium but without glutamine. Respiratory flux was measured with assay cycles of 3 minutes mixing and 3 minutes measurement. Specific inhibitors were used depending on the assay: for mitochondrial respiration, sequential injections of oligomycin (1.5 µM), Bam15 (2 µM, Sigma-Aldrich, SML1760-25MG), and rotenone/antimycin A (0.5 µM) were used (compounds from Agilent kit, 103592-100); for substrate oxidation assays, Etomoxir (4 µM, Cayman Chemical, 11969) was injected; for FAO inhibition, TOFA (Cayman Chemical, 10005263) was included in the assay medium within 1 h of Seahorse XF DMEM equilibration. After the assay, cells were fixed in 100% methanol overnight, stained with 0.5% (w/v) Crystal Violet (Sigma-Aldrich, C0775), and subsequently destained with 1% (w/v) SDS (Sigma-Aldrich, 436143-100G). Absorbance was measured at 560 nm, and these values were used to normalize OCR data in the Agilent Wave Pro software.

### mRNA generation for autophagy flux reporter (RFP-LC3-GFP)

Plasmid utilized for mRNA generation was previously reported (Morita et al 2018). The DNA template for in vitro transcription was generated via PCR amplification from a plasmid source using Q5® High-Fidelity DNA Polymerase (NEB, M0491L). Amplification was performed using primers pMPH403 (TTTTTTTTTTTTTTTTTTTTTTTTTTTTTTTTTTTTTTTTTTTTTTTTTTTTTTTTTTTTTTTTTTTTTTTTTTTTTTTTTTTTTTTTTTTTTTTTTTTTTTTTTTTTTTTTTTTTTTTTTTTTTGCAATGAAAATAAATGTTTTTTATTAGGCAGAATCC) and pMPH404 (TAATACGACTCACTATAAGGACATTTGCTTCTG). To verify size and purity, PCR amplicons were resolved on a 1% (w/v) agarose gel at 140V for 35 minutes. Target bands were excised and purified using NucleoSpin Gel and PCR Clean-up Columns (Takara, 74060950S) according to the manufacturer’s instructions. The concentration and purity of the resulting linear DNA template were quantified prior to use in the IVT reaction.

Capped and polyadenylated mRNAs were synthesized via in vitro transcription using T7 RNA polymerase (NEB, M0251L). To ensure optimal translation and stability, 5’-capping was achieved co-transcriptionally using CleanCap Reagent AG (TriLink, N-7113), while 3’-polyadenylation was encoded directly via a linearized DNA plasmid template containing a poly-A sequence. To minimize innate immunogenicity, pseudouridine-5’-triphosphate (TriLink, N-1019) was substituted for uridine-5’-triphosphate during synthesis. Following transcription, the DNA template was degraded by a 20-minute digestion with DNase I (Promega, M6101). The resulting mRNA was purified using the Monarch® Spin RNA Cleanup Kit (NEB, T2050L) according to the manufacturer’s instructions.

### Lipid Nanoparticle (LNP) Formulation and Encapsulation

LNPs were formulated using an SM-102-based lipid composition (Cayman Chemical, 35425). The lipid mix was prepared at a molar ratio of 50:10:38.5:1.5 (SM-102:DSPC:cholesterol:DMG-PEG 2,000). Purified mRNA was diluted to a concentration of 90µg/mL in a 50mM sodium acetate (pH 4.0) under RNase-free conditions.

Encapsulation was performed using the XGen iNano™ L+ Rapid Nanomedicine Preparation System at a flow rate ratio (FRR) of 3:1 (aqueous:organic). Following synthesis, crude LNPs were immediately diluted in phosphate-buffered saline (PBS). Buffer exchange and LNP concentration were conducted using Amicon Ultra-15 Centrifugal Filter Units (30 kDa molecular weight cut-off; Sigma-Aldrich, UFC903008), centrifuged at 1,000 RPM for 25 minutes per cycle, for a total of 2 cycles. Final LNP preparations were resuspended in PBS supplemented with 10% sucrose, aliquoted, and stored at -80°C for subsequent applications.

### Autophagy flux measurement

Cells were seeded at a density of 10,000 cells per well in a 96-well glass-bottom plate (Cellvis, P96-1.5H-N) and incubated in McCoy’s 5A Modified medium (Gibco, 16600082) supplemented with 10% FBS (Gibco, 10-437-028) at 37°C in a 5% CO2 for 24 h. The culture medium was then replaced with a fresh medium containing 4 ng/uL of RFP-LC3-GFP mRNA encapsulated in lipid nanoparticles (LNPs). Following a 24-h incubation, cells were washed with PBS and cultured in either nutrient-replete McCoy’s 5A Modified medium or HBSS for starvation. Image acquisition was performed using an ImageXpress HCS.ai system equipped with an automated 40X water immersion objective. Live-cell imaging was conducted over a 60-h time course at 3-h intervals. Image analysis was performed using INCarta software to quantify mean GFP and RFP fluorescence intensities, and the RFP:GFP ratio was subsequently calculated to evaluate autophagic flux.

### Statistical analysis

With the exception of proteomics analyses, statistical analyses were performed using GraphPad Prism. Luciferase assay data (Fig. 1G and Fig. 1-9 supplement) are reported as mean ± SD and statistical significance evaluated via one-way ANOVA and two-way ANOVA as indicated. Average predicted distance for intermolecular crosslinks (Fig. 1-6G supplement) is reported as mean ± SD with statistical significance assessed via Student’s t test. All other experimental results are represented as mean ± SEM of replicates and were analyzed using Student’s t test, one-way ANOVA, or two-way ANOVA followed by a post hoc Tukey honestly significantly different (HSD) test.

## Acknowledgements

The authors thank our senior laboratory manager, Katharine Mellman for her operational support throughout this work. The authors thank all of the hubs and facilities at Altos Labs, including the Bioinformatics, Flow Cytometry, Light Microscopy, Mass Spectrometry, and Cell Models & Cell Engineering hubs for their invaluable technical support, expertise, and infrastructure. The authors acknowledge Tingting Wang and Mengyu Ge at Viva Biotech (Shanghai) Ltd. for SEC-MALS experiments and data analysis, Ziqi Chen and Jiafu Li at Viva Biotech (Shanghai) Ltd. for protein expression and purification, and Mengyu Ge for Altos account management.

---

**Figure 1-1 supplement:**
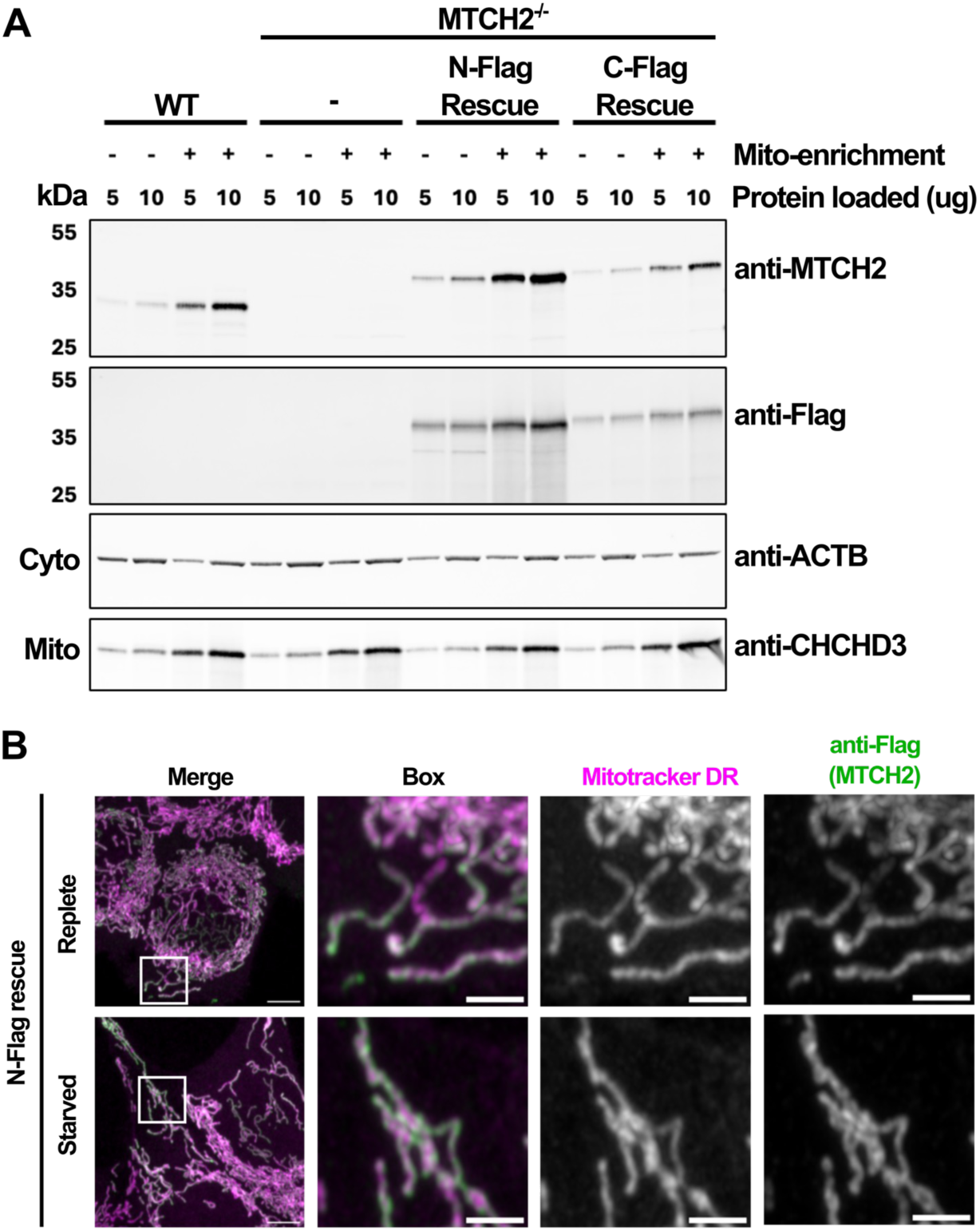
Validation of N- and C-terminally tagged MTCH2 expression in rescue cell lines for topology studies. (A) Validation of rescue cell lines expressing MTCH2 with N- or C-terminal 3xFlag tag by immunoblot of total or mito-enriched fractions in comparison to cytosolic (ACTB) or mitochondrial (CHCHD3) markers. Two different amounts of protein were loaded per lane as designated above each well. (B) Representative IF microscopy images of N-terminally 3xFlag-tagged MTCH2 rescue cells under replete and starved conditions. Subcellular localization was assessed by the colocalization of the Flag epitope tag (green) with MitoTracker Deep Red staining (magenta), scale bar = 5 um, inset box = 2 um.

**Figure 1-2 supplement:**
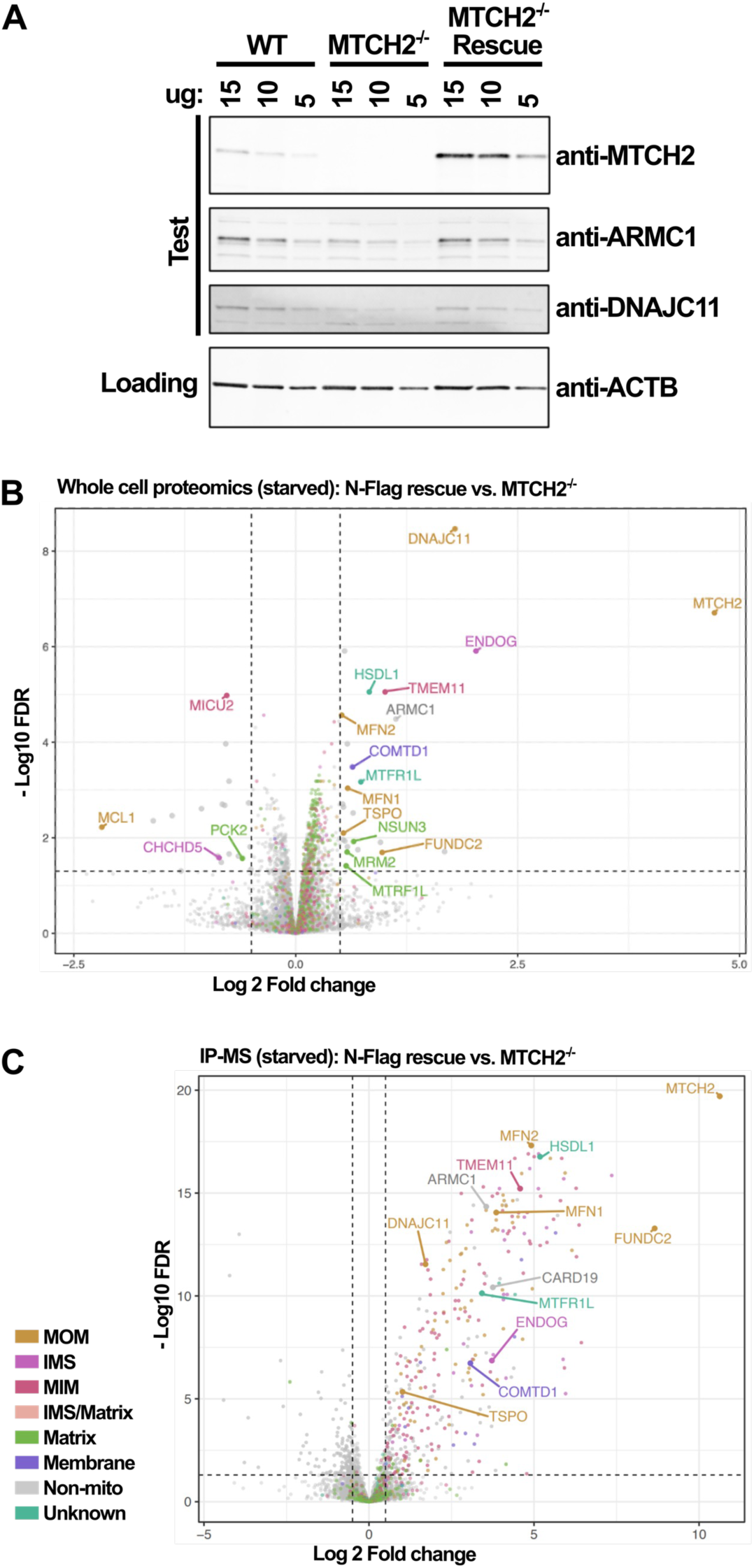
Whole cell and IP proteomics analyses for MTCH2 during starvation. (A) Immunoblot confirming complementation of untagged MTCH2 restores abundance of ARMC1 and DNAJC11 under replete conditions. Three different amounts of protein loaded (as indicated above blots) and resolved on precast 4-15% gradient polyacrylamide gels, visualized by antibodies against the indicated endogenous proteins. (B) Unbiased whole cell proteomics of MTCH2 rescue cell lines with N-terminal 3xFlag (right) compared to MTCH2^-/-^ under starvation, depicted by volcano plot (FDR < 0.05, fold-change cutoff set to 0.5, n = 5 replicates). (C) Enrichment of proteins co-purified with MTCH2 depicted by volcano plot (N-terminal 3xFlag-tagged MTCH2, right, relative to WT, left) with Flag tag used as bait under starved conditions. Proteins color-coded by subcompartment as designated and hits co-dependent with MTCH2 under the same conditions labeled (FDR < 0.05, fold-change cutoff set to 0.5, n = 5 replicates).

**Figure 1-3 supplement:**
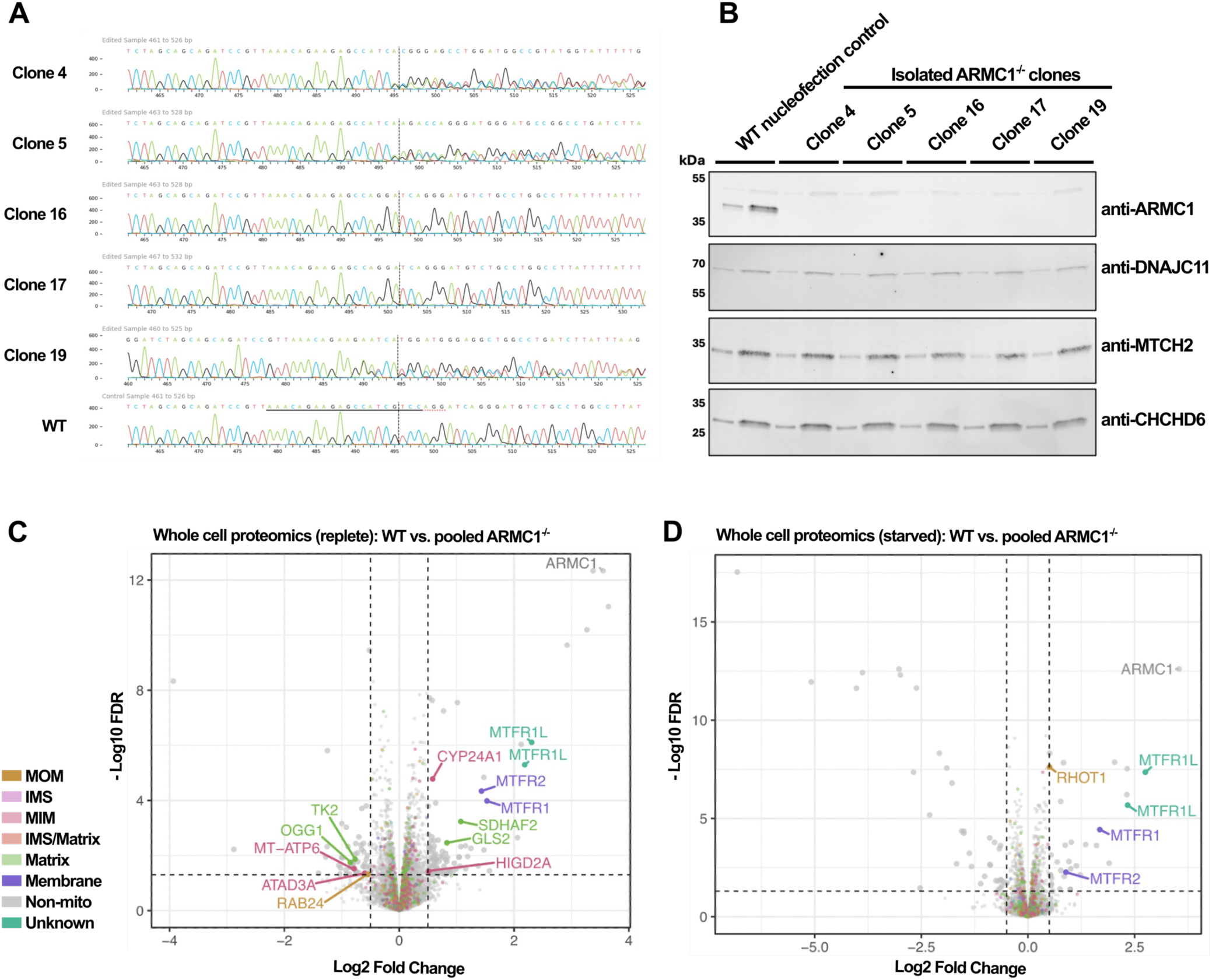
Validation of ARMC1 KO clones and determination of co-dependent proteins. (A) Chromatogram alignment from Sanger Sequencing across CRISPR Cas9 edit site in WT and 5 independently isolated ARMC1 KO clones aligned. (B) Loss of protein expression validated by immunoblot with antibody against endogenous ARMC1 for 5 independent ARMC1 KO clones in comparison to WT. (C) Investigation of proteins co-dependent with loss of ARMC1 by whole cell proteomics under replete and (D) starved conditions. Mitochondrial proteins colored by compartment and labeled, non-mitochondrial proteins depicted in grey (FDR < 0.05, fold-change cutoff set to 0.5, n = 5 replicates).

**Figure 1-4 supplement:**
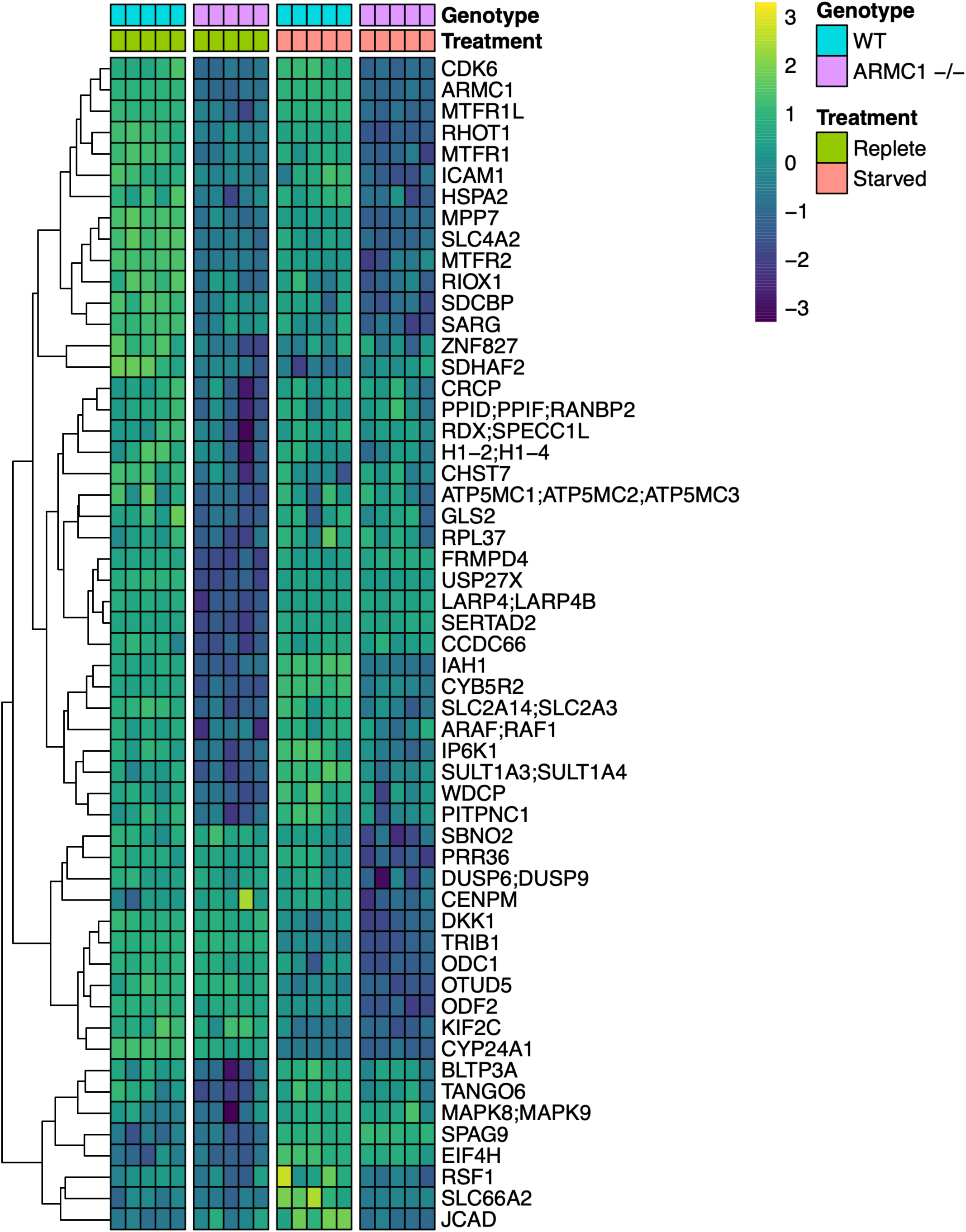
Unbiased proteomics analysis heatmap for differentially enriched proteins in ARMC1 KO. (A) Five independent, validated ARMC1 KO clones were pooled and whole cell proteomics analyses performed in comparison to WT mock nucleofection control under replete and starved conditions. Differentially enriched proteins in either condition defined with fold change (FC) > 0.5 and p-value of < 0.01 were depicted in clustered heatmap for a total of 45 unique hits (41 and 34 differentially enriched proteins identified under replete and starved conditions, respectively).

**Figure 1-5 supplement:**
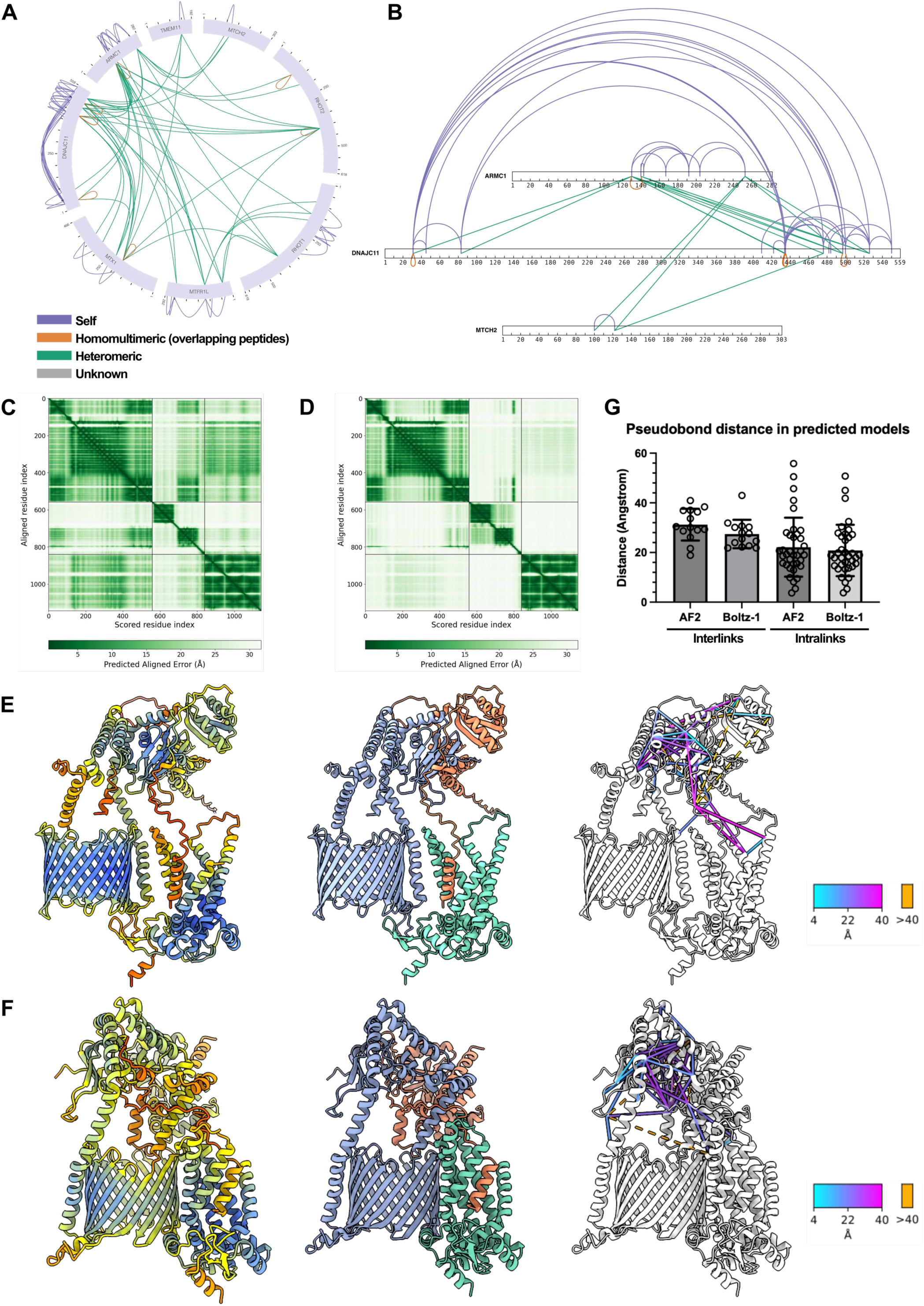
Evaluation of MTCH2-ARMC1-DNAJC11 Boltz-1/2 and AF2 multimer models using published crosslinking mass spectrometry data. (A) Self (purple), homomultimeric (orange), and heteromeric (teal) crosslinks were mapped using publicly available XL-MS data ^23^ for ARMC1, DNAJC11, MTCH2, MTX1, and TMEM11 using xiVIEW^51^. (B) Inter and intramolecular crosslinks mapped for ARMC1, DNAJC11, and MTCH2 by xiVIEW along the length of each protein. (C) and (D) comparison of PAE matrices for AF2 multimer and Boltz-1/2 modeling of MTCH2-ARMC1-DNAJC11 complex. (E and F) Ribbon structures for predicted complex colored by pLDDT score (left), by color (center): MTCH2 in teal, ARMC1 in coral, and DNAJC11 in lavender, and color-stripped with crosslinks mapped to 3D structure (right). Distance in space between crosslinked residues designated with scale bars ranging from 4-40 Angstrom with crosslinks >40 Angstrom depicted in orange. (G) Comparison of inter- and intramolecular crosslink distances predicted by either AF2 or Boltz-1/2 in Angstrom, (each dot = 1 interlink).

**Fig 1-6 supplement:**
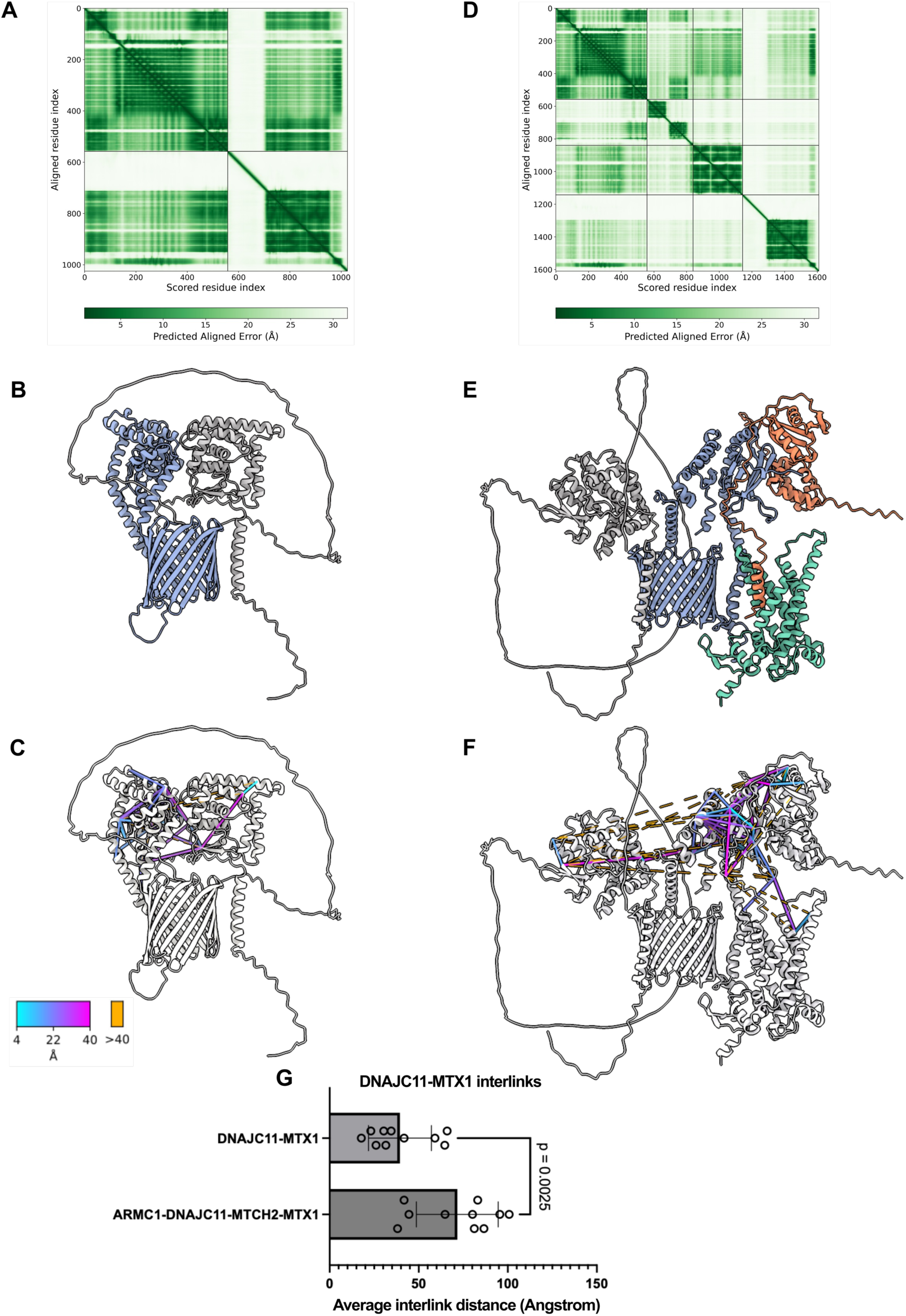
Published crosslinking mass spectrometry data does not support ARMC1-DNAJC11-MTCH2-MTX1 model. Comparison of AF2 multimer model of DNAJC11-MTX1 heterodimer (A-C) with ARMC1-DNAJC11-MTCH2-MTX1 versus (D-F). (A) PAE matrix for DNAJC11-MTX1 supporting 3D structure in (B, DNAJC11 in lavender and MTX1 in light grey). (C) DNAJC11-MTX1 ribbon structure color-stripped incorporating inter- and intra-molecular crosslinks extracted from publicly available XL-MS data ^23^. Distance in space between crosslinked residues designated with scale bars ranging from 4-40 Angstrom with those >40 Angstrom depicted in orange. (D-F) Same as (A) through (C) but for MTX1-containing heterotetramer (ARMC1 in coral, DNAJC11 in lavender, MTCH2 in teal, and MTX1 in light grey). (G) Comparison of DNAJC11-MTX1 interlink distance between MTX1-containing heterodimer and heterotetramer (each dot = 1 interlink), statistical significance assessed via Student’s t-test.

**Figure 1-7 supplement:**
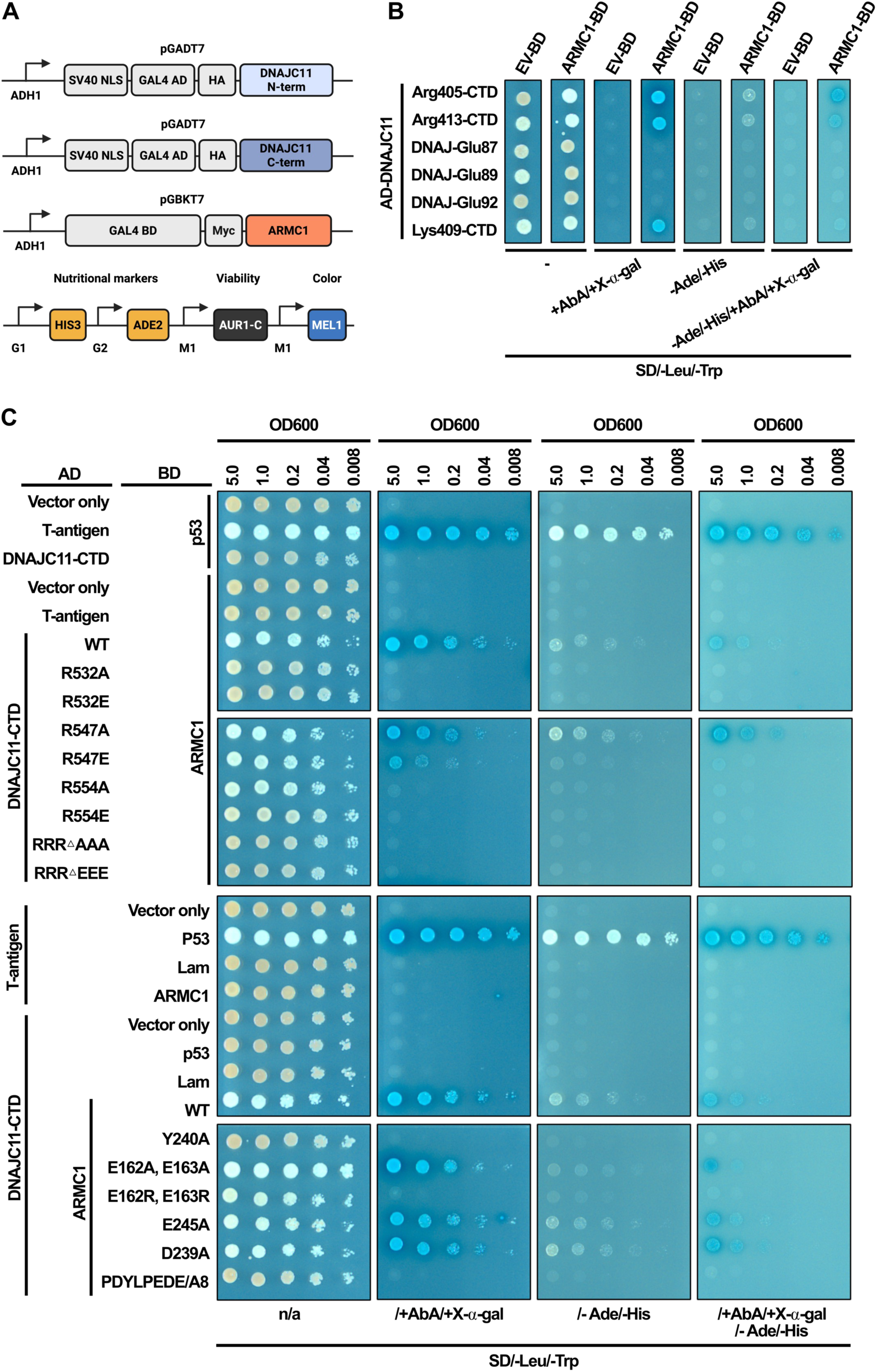
ARMC1 interacts with DNAJC11 C-terminus predominantly via acidic linker and HMAL domain. (A) Diagram for recombinant yeast 2-hybrid constructs of full length ARMC1 and either DNAJC11 DNAJ domain or C-terminus as well as selectable markers for positive interactions (lower panel). Created in BioRender. Castonguay, A. (2026) https://BioRender.com/poxn2dn. (B) Spot assay for yeast two-hybrid assay between recombinant, full-length ARMC1 and DNAJC11 N-terminal DNAJ domain or C-terminus each initiated at 3 different positions. Interaction was visualized on selective media of increasing stringency of interaction vs. nonselective (SD, -Leu, -Trp). (C) Amino acid substitutions targeting interaction between DNAJC11 C-terminus and ARMC1 were introduced testing disruption of positive interaction observed in (B) but via spot assay with 5-fold serial dilutions in comparison to p53:T-antigen positive and empty vector negative controls.

**Figure 1-8 supplement:**
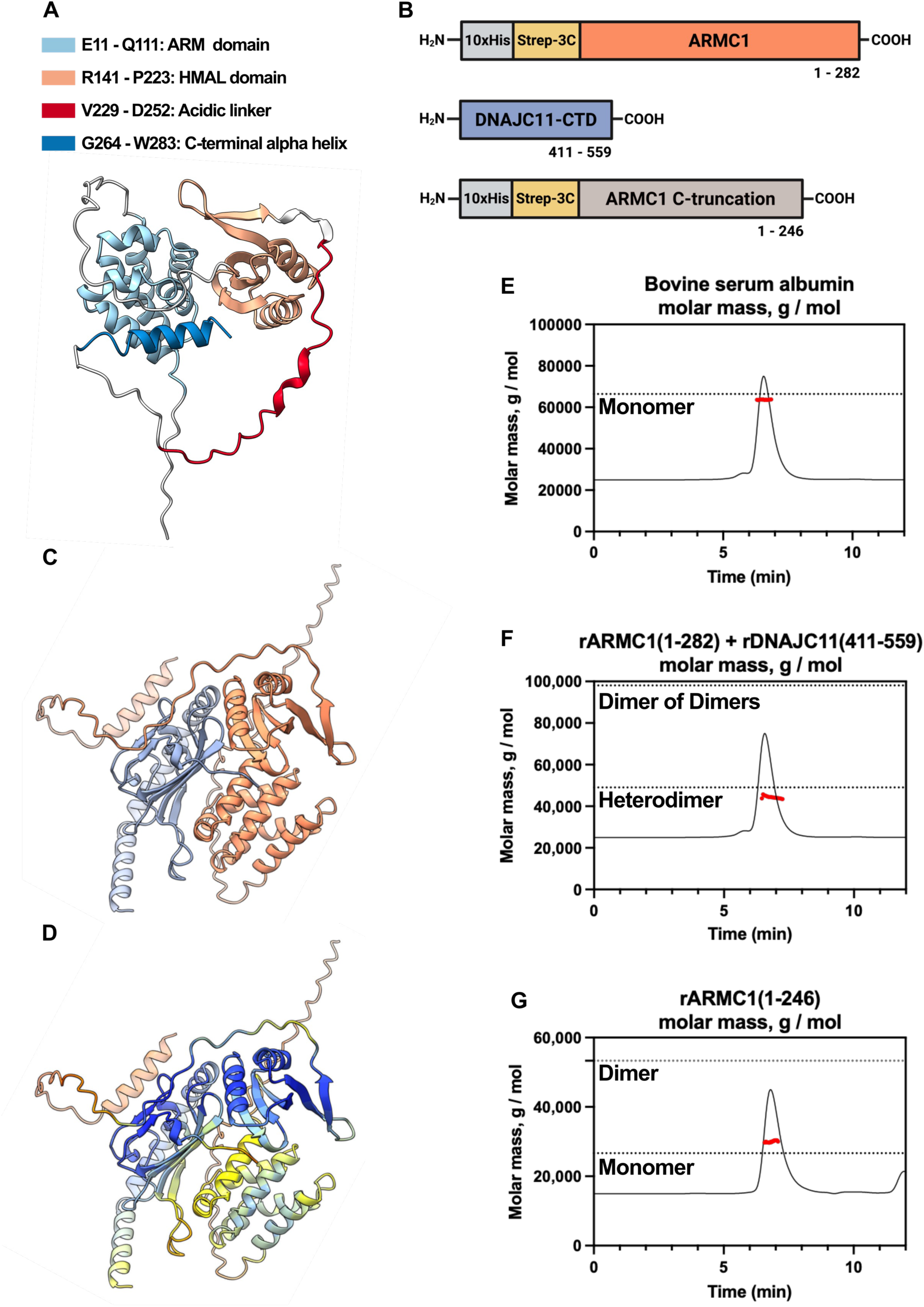
Purification of full length ARMC1 requires co-expression with DNAJC11 C-terminus. (A) AF2 model of ARMC1 colored by domain as designated. (B) Schematic of recombinant constructs for (co)expression and (co)purification from *E. coli*. Empirical molecular weight for 10x-His-Strep-3C-ARMC1(1-282) was 35,538.93 Da and 31,893.92 Da pre- and post-cleavage at 3C site, respectively, and DNAJC11(411-559) was 17,132.74 Da. Created in BioRender. Castonguay, A. (2026) https://BioRender.com/r3hu4a5. Empirical molecular weight of 10x-His-Strep-3C-ARMC1(1-246) was 31,588.62 Da and 27,943.61 Da pre- and post-cleavage, respectively. (C) Interaction of ARMC1-FL with DNAJC11 C-terminus modeled by AF2 multimer using CollabFold (ARMC1 in coral, DNAJC11 in lavender) and (D) colored by pLDDT score. (E - G) Size Exclusion Chromatography with Multi-Angle Light Scattering (SEC-MALS) for BSA control (E), 10x-His-Strep-3C-ARMC1(1-282):DNAJC11(411-559) (F), and 10x-His-Strep-3C-ARMC1(1-246) (G). Empirical and experimental molecular weight is designated with dashed line and red dots, respectively.

**Figure 1-9 supplement:**
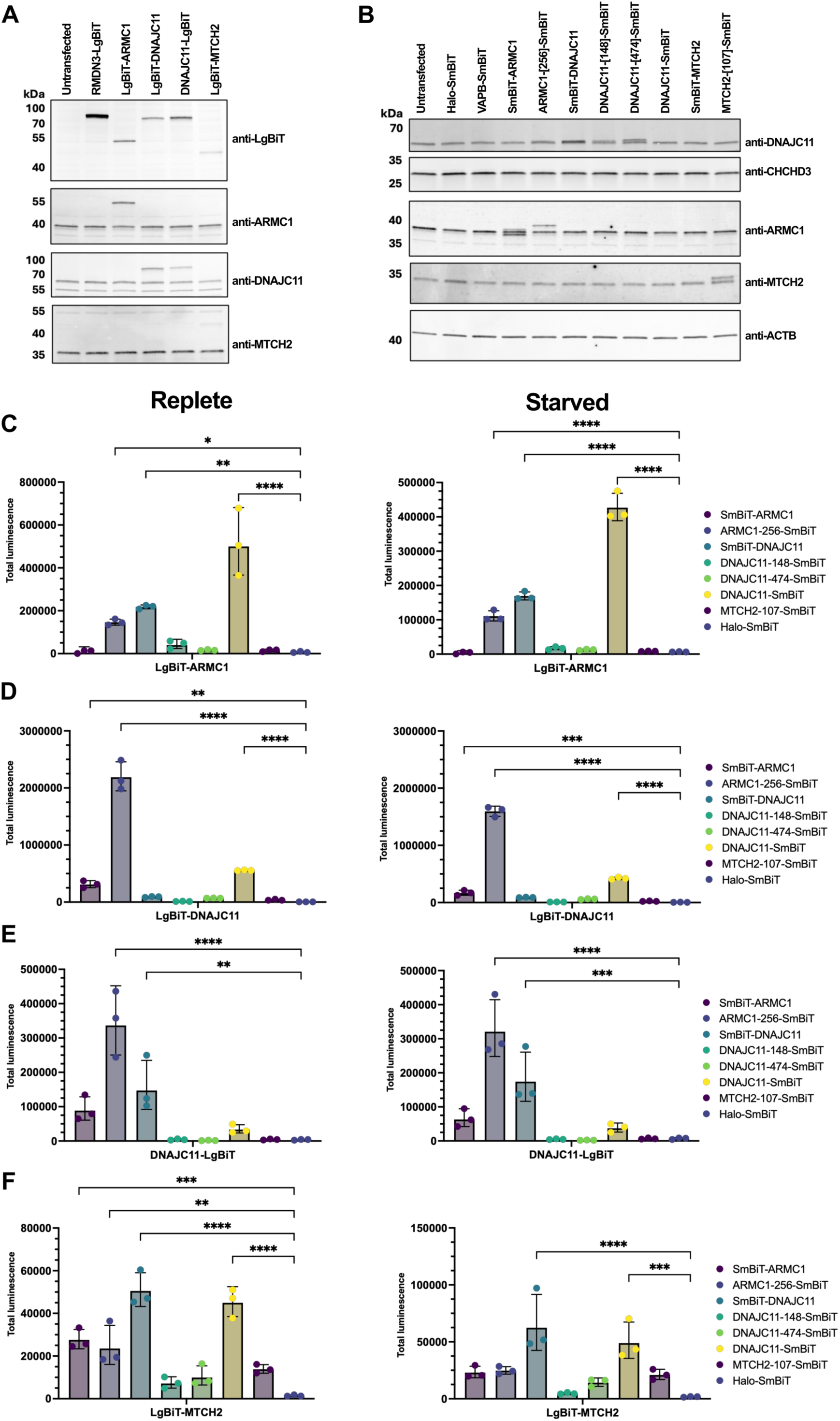
Validation for chimeric reporter expression for NanoBiT assay and test of interactions. (A) Evaluation of transient expression of LgBiT chimeric proteins by immunoblot. Recombinant protein was validated by detection with anti-LgBiT antibody and antibodies against endogenous protein with corresponding size shift in comparison to untransfected control. (B) Expression of SmBiT-tagged recombinant proteins via immunoblot. No specific antibody against the SmBiT tag was available so ectopic expression of SmBit-tagged proteins was verified using antibodies against the endogenous protein. NanoBiT assay of cell cultures under replete (left) and starved (right) conditions transiently co-expressing recombinant LgBiT-tagged ARMC1 (C), DNAJC11 (D, E), or MTCH2 (F) and potential SmBiT containing split luciferase pairs. Total luminescence was measured for each combination and depicted by histogram (n = 3). Combination with Halo-SmBiT served as negative control for nonspecific interactions. Optimization of split luciferase assay was done using RMDN3-LgBiT and VAPB-SmBiT positive control described previously ^52^ in comparison to Halo-SmBiT negative control (gray bars). Statistically significant differences within each set are designated with p-value above each comparison (one-way ANOVA, no P-value = nonsignificant). No statistically significant difference in luciferase activity was observed with LgBiT-MTCH2 expressing cells between replete and starved conditions (2-way ANOVA).

**Figure 2-1 supplement:**
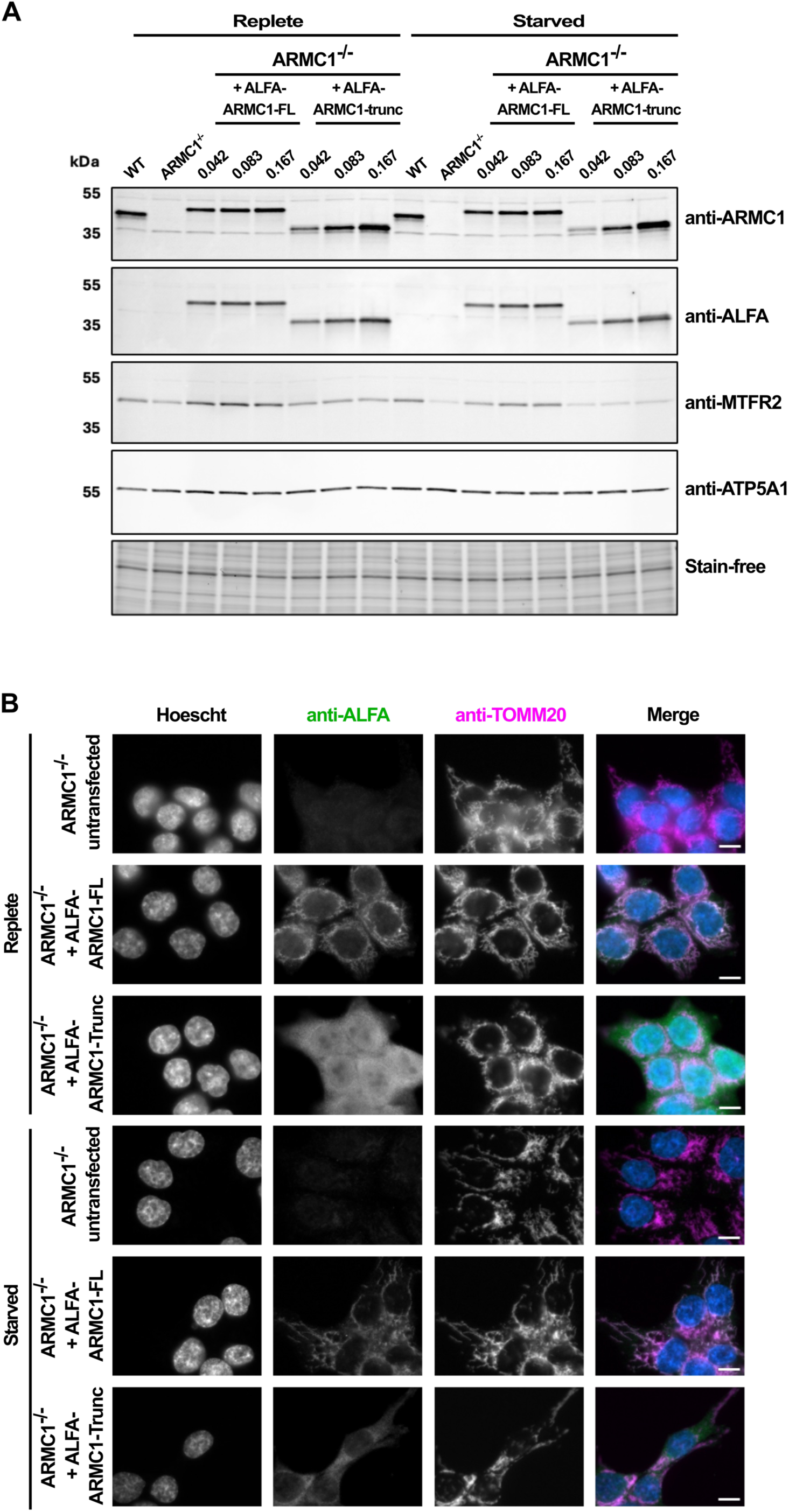
Mitochondrial enrichment of ALFA-ARMC1 is dependent on its CTD. (A) ARMC1^-/-^ cells were transiently transfected with increasing amounts of LNPs delivering ARMC1-FL or ARMC1-trunc. with N-terminal ALFA tag and expression evaluated by SDS-PAGE and immunoblot using protein and tag specific antibodies. Volume designated above each lane corresponds to volume of LNP/10,000 cells. (B) Representative IF microscopy images showing anti-ALFA and anti-TOM20 (mitochondria) staining in ARMC1^-/-^ cells transiently transfected with 0.167 uL of LNP/10,000 cells expressing N-terminally ALFA tagged ARMC1-FL or a C-terminally truncated ARMC1. Scale bar = 10 µm.

**Figure 2-2 supplement:**
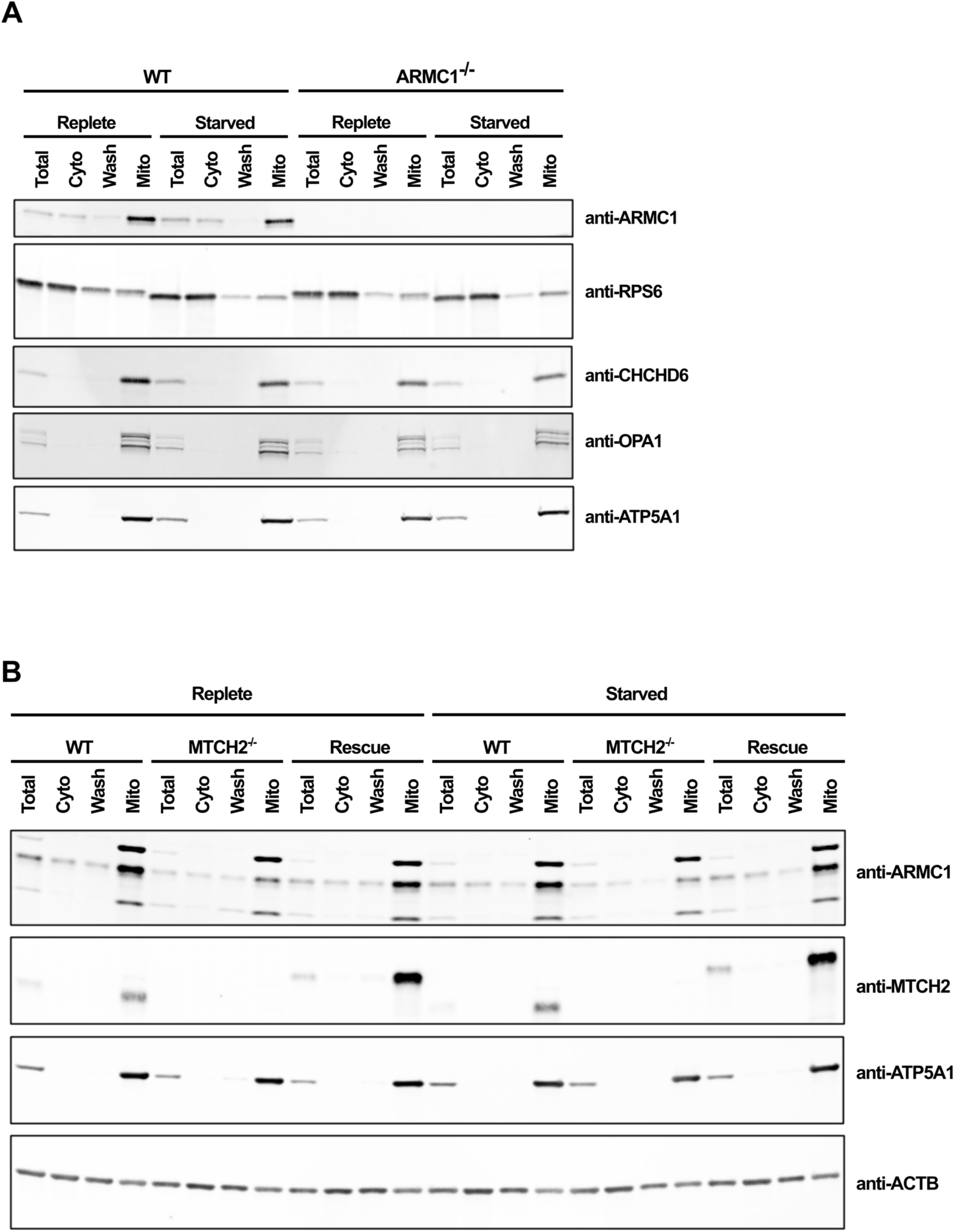
Loss of MTCH2 decreases mitochondrial and cytosolic pools of ARMC1. (A) WT and ARMC1^-/-^ cells were cultured under replete or starved conditions then subcellular fractions resolved by SDS-PAGE. ARMC1 was detected by immunoblotting with antibody against endogenous protein in comparison to compartmental markers (RPS6 = cytosol, CHCHD3 = IMS/IMM, OPA1 = IMM, ATP5A1 = matrix). (B) Same as (A) but for WT, MTCH2^-/-^, and MTCH2 N-terminal rescue cell lines compared to compartment markers (ATP5A1 = mitochondria, ACTB = cytosol).

**Figure 2-3 supplement:**
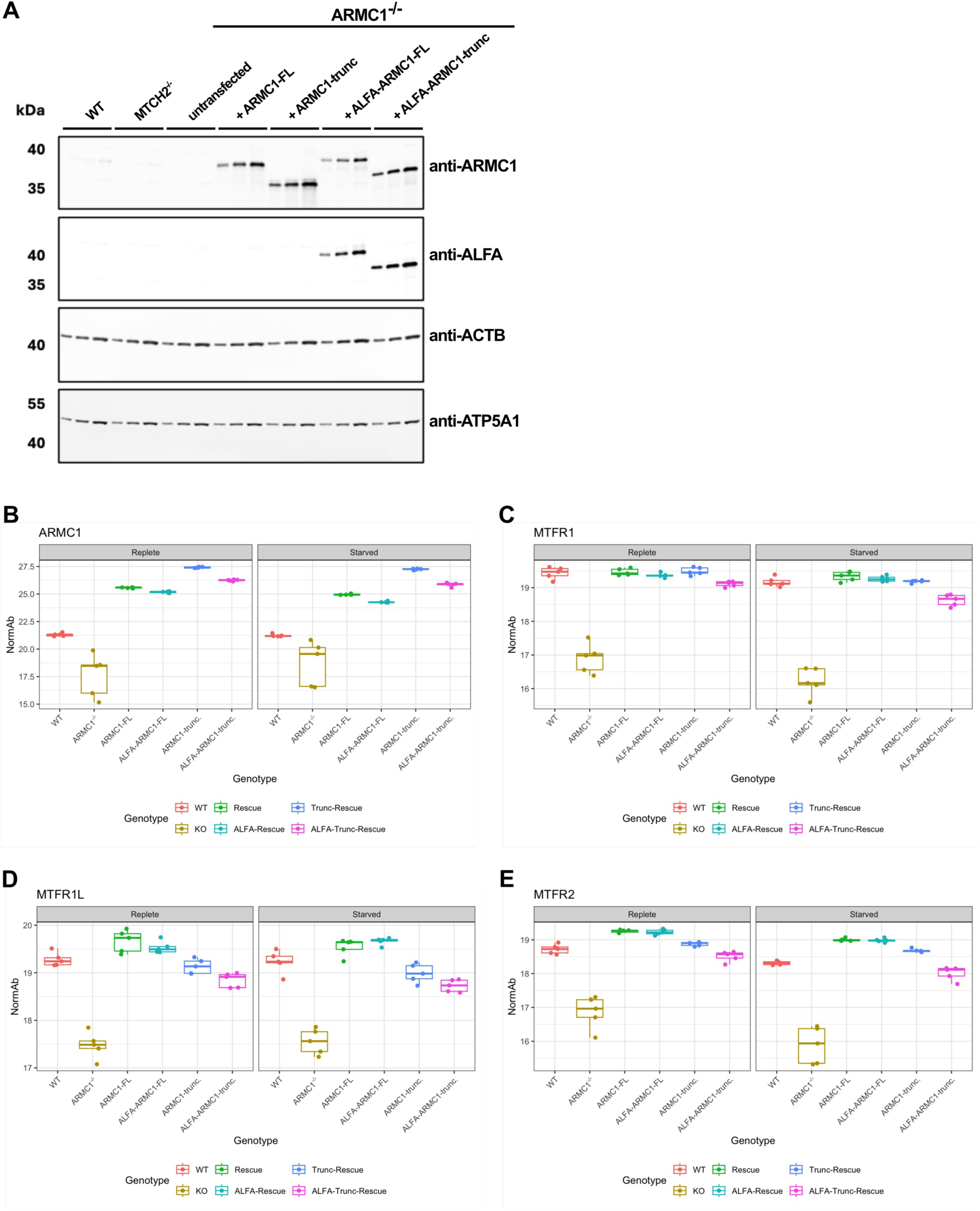
Validation of stable ARMC1-overexpressing cell lines. (A) To rescue ARMC1 expression in ARMC1^-/-^ lines, we constitutively overexpressed full-length ARMC1 as well as a C-terminally truncated variant either untagged or with a N-terminal ALFA tag. Increasing amounts of protein (2, 4, and 8 ug) from whole cell lysates were resolved and immunoblotted with anti-ARMC1 or anti-ALFA antibodies. (B-E) Whole cell lysates collected from WT, ARMC1^-/-^, and cell lines overexpressing full length or truncated ARMC1 with and without N-terminal ALFA tag were submitted for unbiased mass spectrometry. Initial proteomics analyses focused on rescue of ARMC1 and the MTFR proteins (MTFR1, MTFR2, and MTFR1L) previously reported to be co-depleted with loss of ARMC1 depicted by box and whiskers plot, n = 5 replicates/condition.

**Figure 2-4 supplement:**
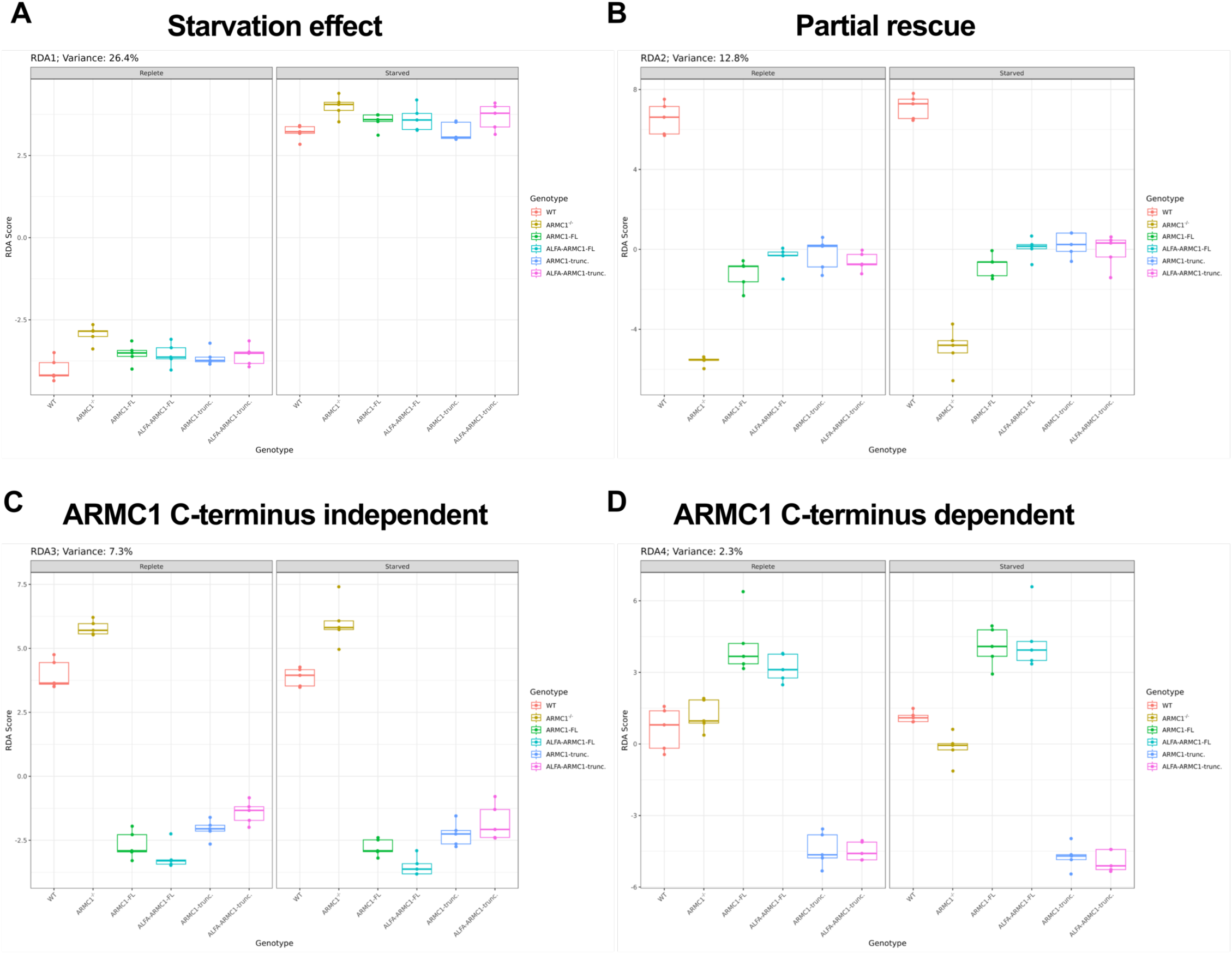
Whole cell proteomics and redundancy analysis in ARMC1^-/-^ cells rescued with full-length or C-terminal truncated ARMC1. (A) Unbiased mass spectrometry was performed on whole cell lysates collected from WT, ARMC1^-/-^, and cell lines overexpressing full length or truncated ARMC1 (with and without N-terminal ALFA tag). Proteomics analyses highlighting change in abundance of ARMC1 displayed by box and whiskers plot, n = 5 replicates/condition. (B-E) Redundancy analysis (RDA) for ARMC1 overexpression dataset highlighting top 4 RDs, ordered by decreasing contribution to variation and designated as (B) media/starvation response, (C) partial rescue of ARMC1^-/-^, (D) ARMC1 C-terminus independent (cytosolic) rescue, and (E) ARMC1 C-terminus dependent (mito-enriched) rescue.

**Figure 2-5 supplement (page 1-2):**
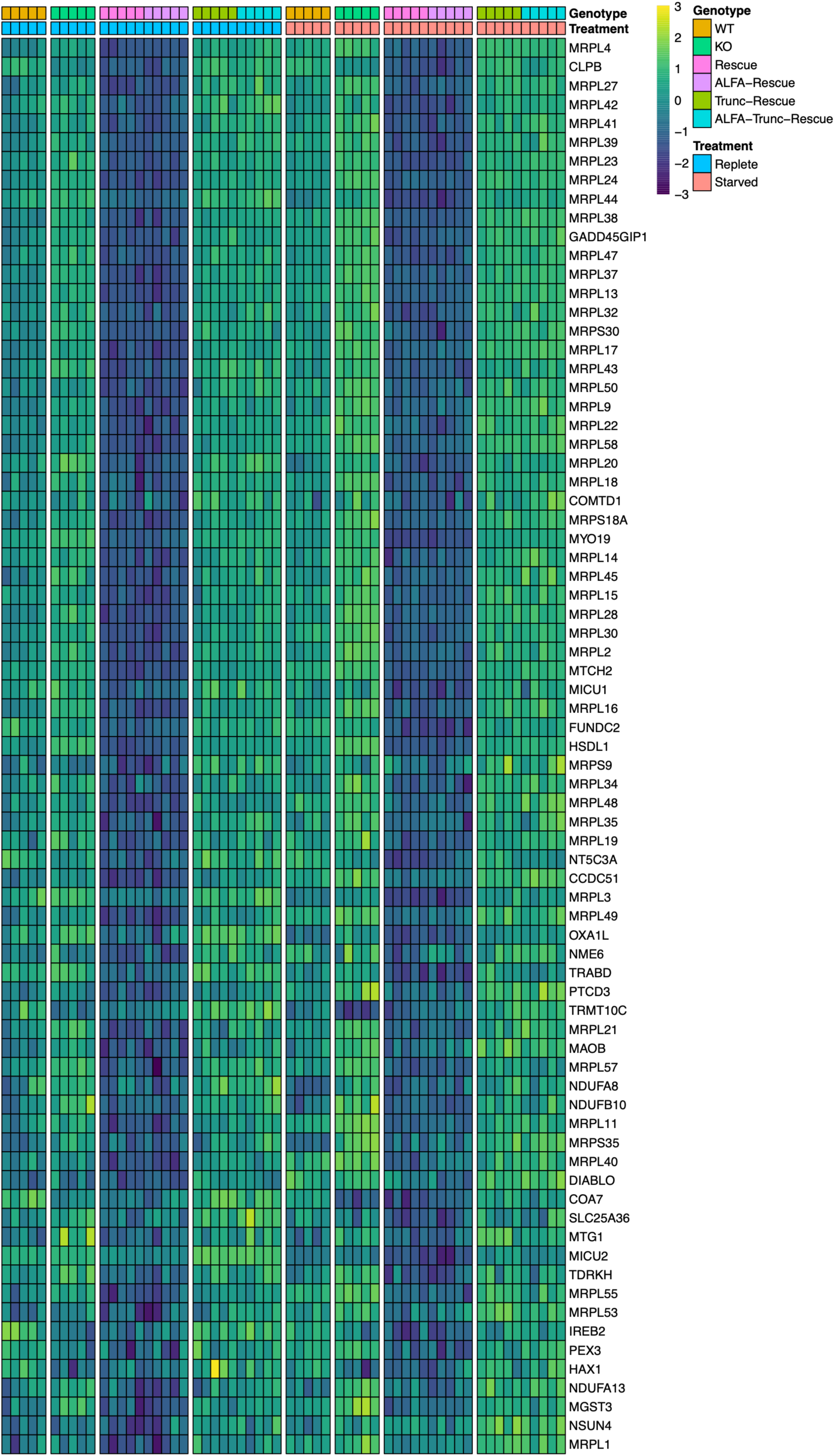
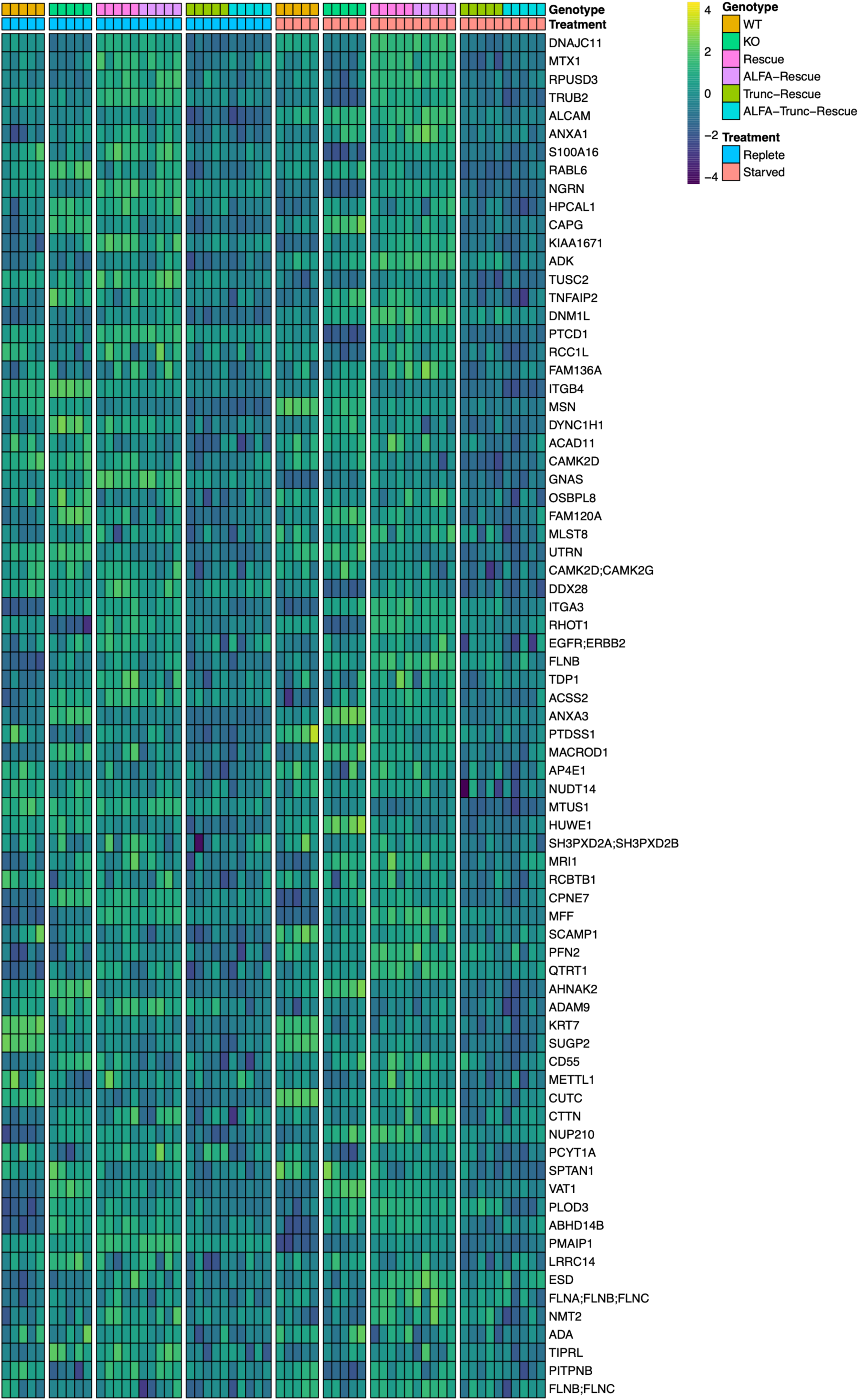
Unbiased proteomics analysis heatmap for proteins positively and engatively contributing to ARMC1 C-terminus dependent RD. (A-B) Top 75 proteins (A) negatively and (B) positively contribute to ARMC1 C-terminus dependent rescue RD separation (from Figure 2-4D supplement) depicted by heatmap.

**Figure 3-1 supplement:**
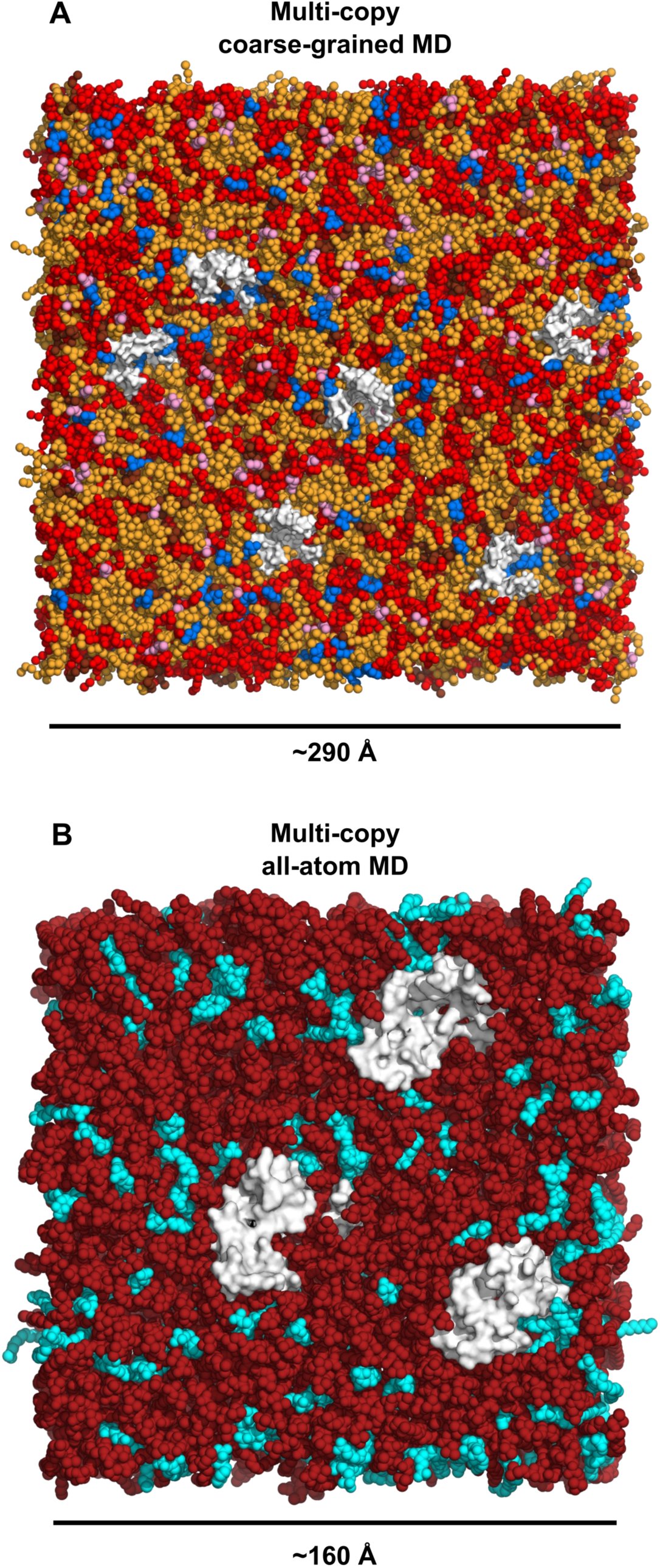
Overview of CG- and AA-MD simulation systems. (A) Visualization of an example configuration of a CG-MD simulation system used to assess lipid scrambling, viewed towards the cytoplasmic face of the membrane; six copies of MTCH2 embedded in the membrane are shown as white Van der Waals surfaces; lipid CG particles are shown as spheres: orange POPC, red POPE, blue POPI, brown POPA, and pink LPA. (B) Visualization of an example configuration of an AA-MD simulation system used to assess lipid scrambling and interactions, viewed towards the cytoplasmic face of the membrane; three copies of MTCH2 embedded in the membrane are shown as white Van der Waals surfaces; lipid heavy atoms are shown as spheres with POPC in dark red and POPA in cyan.

**Figure 3-2 supplement:**
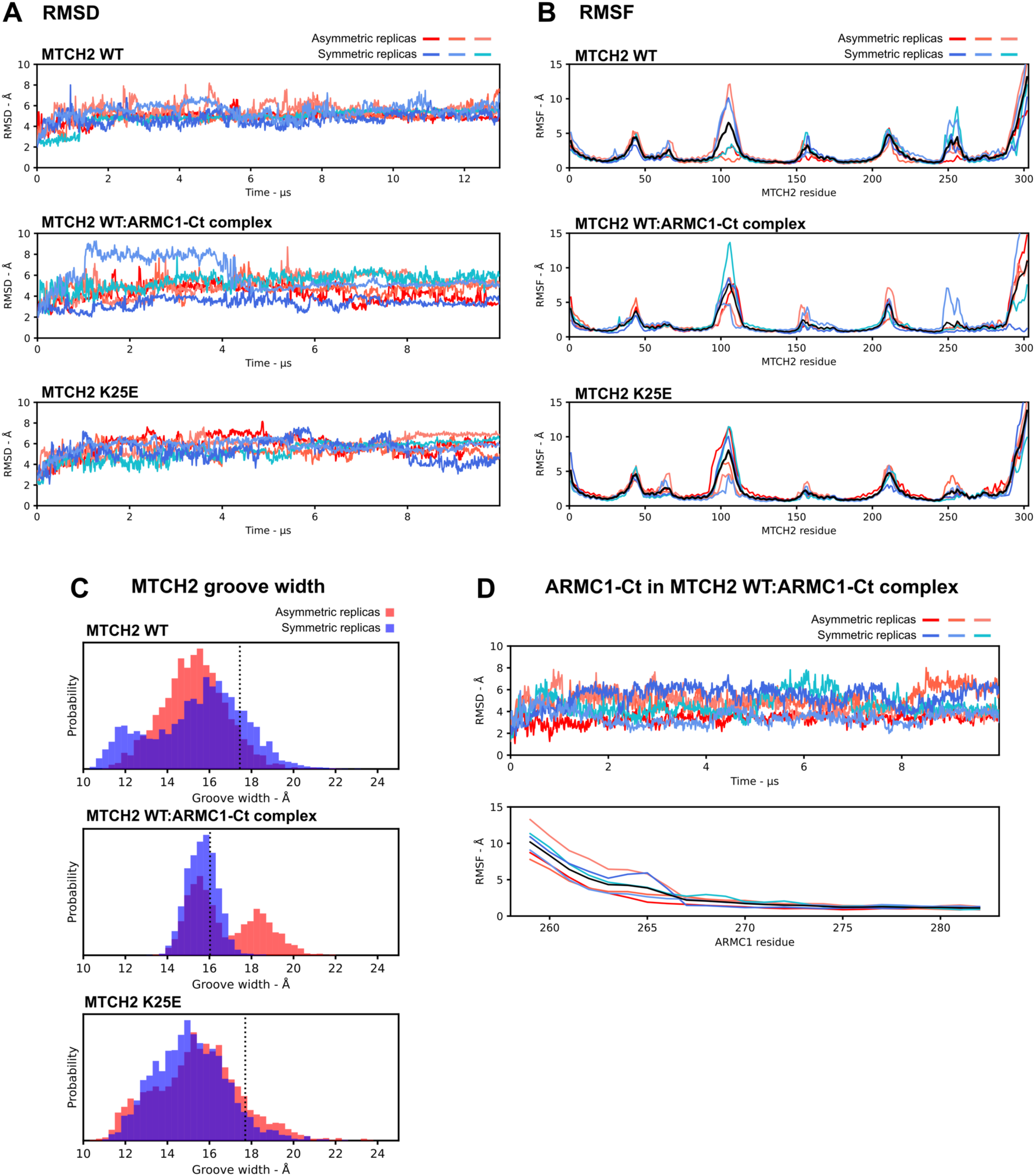
Predicted structural models are stable for microseconds of AA-MD simulation. (A) Cα root-mean-squared-deviation (Cα RMSD) for MTCH2 in various simulation configurations, calculated after all-Cα alignment on MTCH2 for each trajectory/replica to its initial structural model. (B) Cα time-average-root-mean-squared-fluctuation (Cα RMSF) calculations for MTCH2 in various simulation configurations, calculated after all-Cα alignment to initial structural models and excluding conformations in the first 2 μs of each trajectory. The black line indicates the average RMSF over all six replicas. (C) Histograms showing the distribution of distances between Cα atoms of MTCH2 residues 14 and 224, representing the width of the hydrophilic groove. Distributions were calculated excluding the first 2 μs of each trajectory. Red and blue histograms aggregate across all replicas in asymmetric and symmetric bilayers respectively for each condition. Vertical black dotted lines in each panel indicate the distance in the initial structural models. (D) Cα RMSD and Cα RMSF of the ARMC1 C-terminus chain from MTCH2 WT:ARMC1-Ct complex simulations. All values are calculated after alignment of the trajectory to the corresponding MTCH2 chain to which ARMC1 is bound, not the ARMC1 chain itself. RMSD and RMSF traces in panels A, B, and D are colored shades of red for asymmetric bilayer trajectories and shades of blue for symmetric bilayer trajectories.

**Figure 3-3 supplement:**
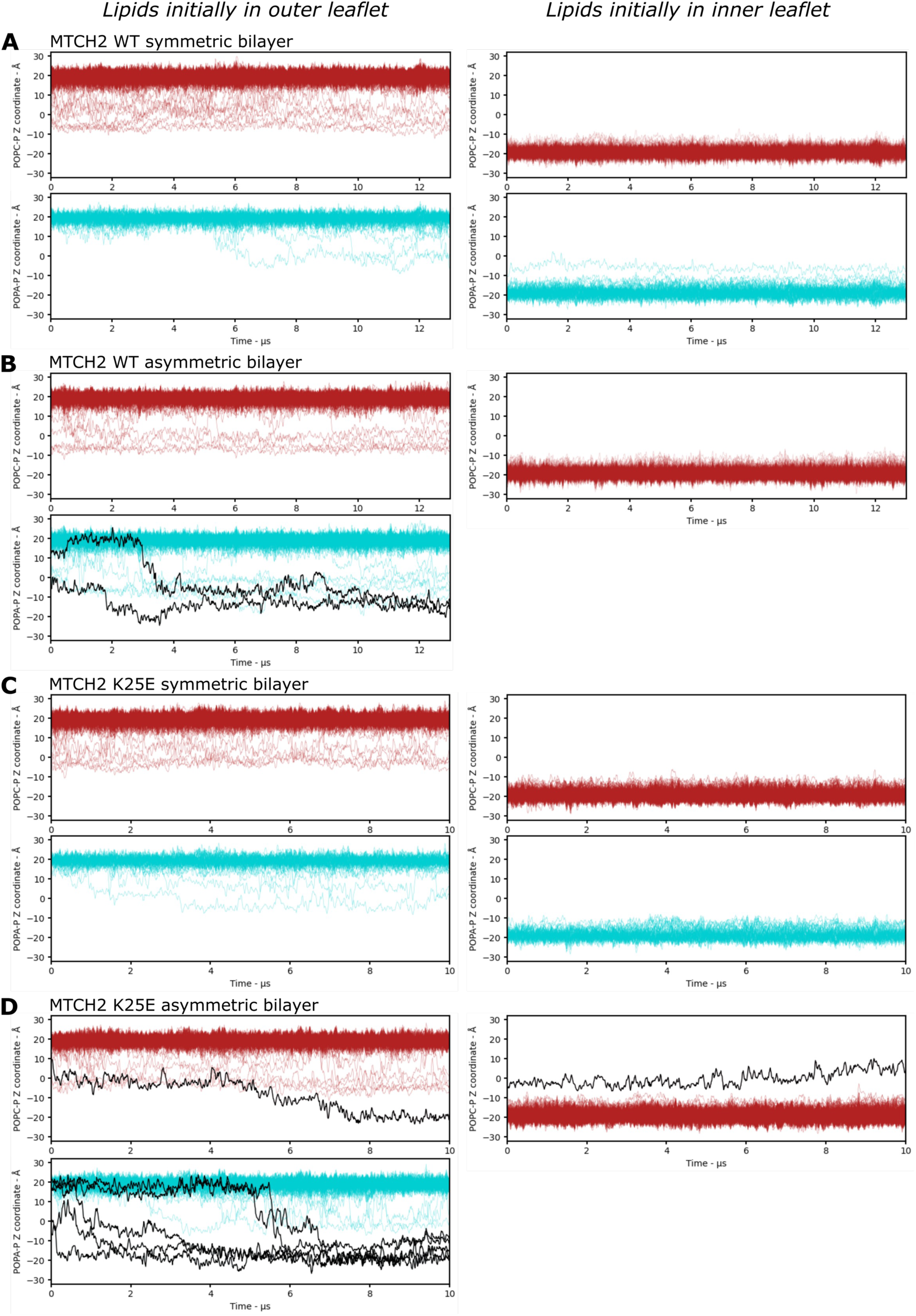
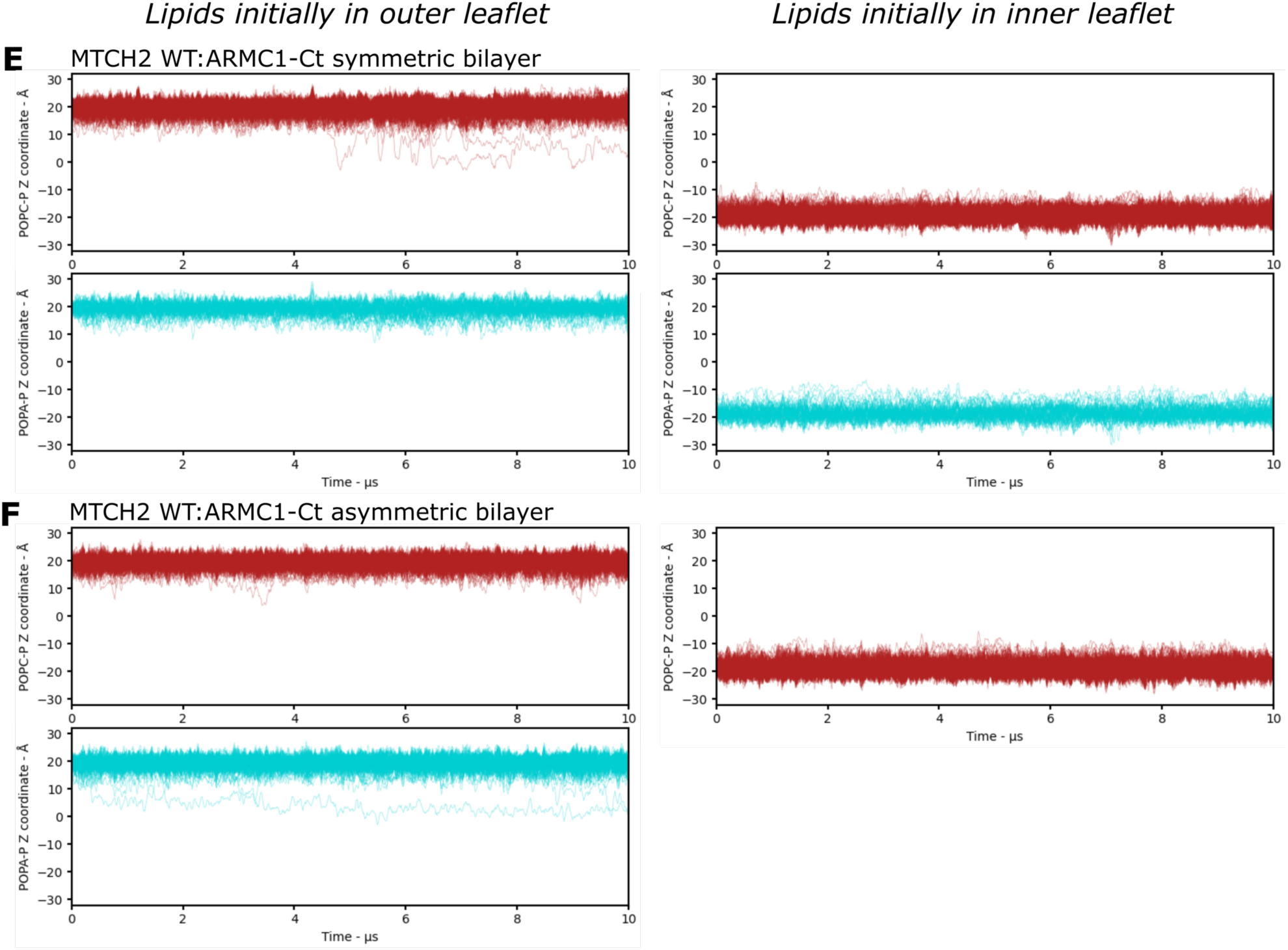
Individual lipid scrambling trajectories across all AA-MD simulation conditions. Each of A, B, C, D, E, & F shows the trajectories of Z-coordinates of every lipid phosphorus atom in each respective AA-MD simulation as indicated. For each simulation trajectory, lipids are broken out into 4 or 3 groups; POPC lipids initially in the cytoplasmic leaflet (upper left panels, dark red lines), POPC lipids initially in the IMS leaflet (upper right panels, dark red lines), POPA lipids initially in the cytoplasmic leaflet (lower left panels, cyan lines), or POPA lipids initially in the IMS leaflet (lower right panels, cyan lines; symmetric bilayers only). All lipids that translocate fully from either 1) the cytoplasmic leaflet to the bulk IMS leaflet via the hydrophilic groove, or 2) from the bulk IMS leaflet to the hydrophilic groove or cytoplasmic leaflet have trajectories highlighted as black lines and were verified by manual inspection of simulation trajectories. The dark bands around either Z=+18 or Z=-18 in each panel are many overlapping trajectories from lipids within the bulk membrane, largely not interacting with protein, while sparser lines in the intermediate region represent lipids interacting with protein, typically within the hydrophilic groove, but also at peripheral sites that shift their Z-coordinates away from those of bulk lipids.

**Figure 3-4 supplement:**
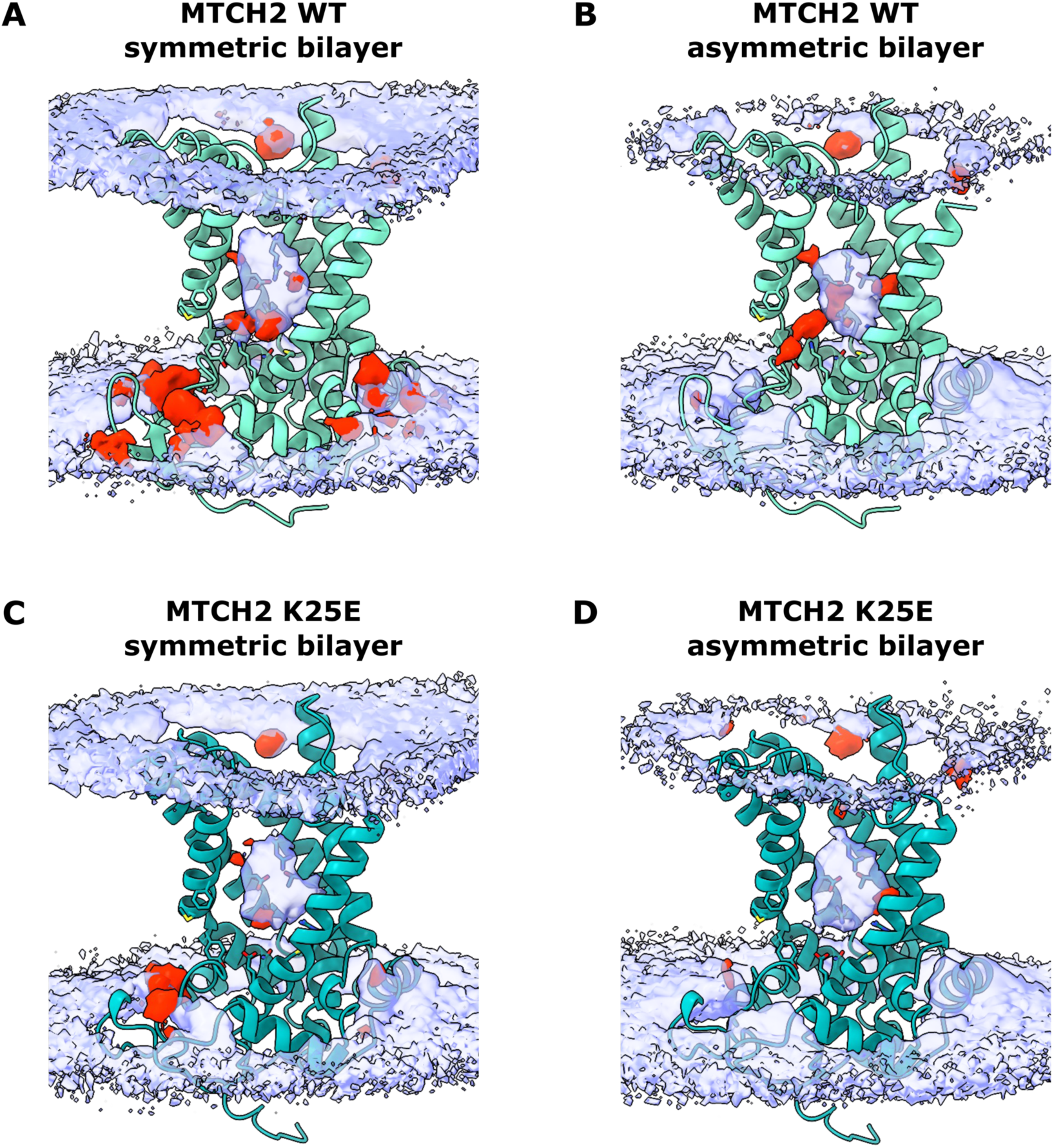
Headgroups binding at discrete MTCH2 sites in and around the hydrophilic groove depends on the arrangement of charged residues. Each of A, B, C, & D shows lipid headgroup atom density maps calculated for each respective AA-MD simulation as indicated. Average protein structures from the simulations are shown in a cartoon representation with MTCH2 WT in green (A & B) & MTCH2 K25E in dark teal (C & D). Side chains of certain lipid contacting residues lining the hydrophilic groove are shown as sticks. Opaque red density maps represent histograms of POPA phosphate groups; transparent lavender density maps represent histograms of POPC choline groups. All maps in all panels for both headgroup types are contoured at an equivalent level representing equal degree of occupancy of the enclosed voxel by the respective atom groups.

**Figure 4-1 supplement:**
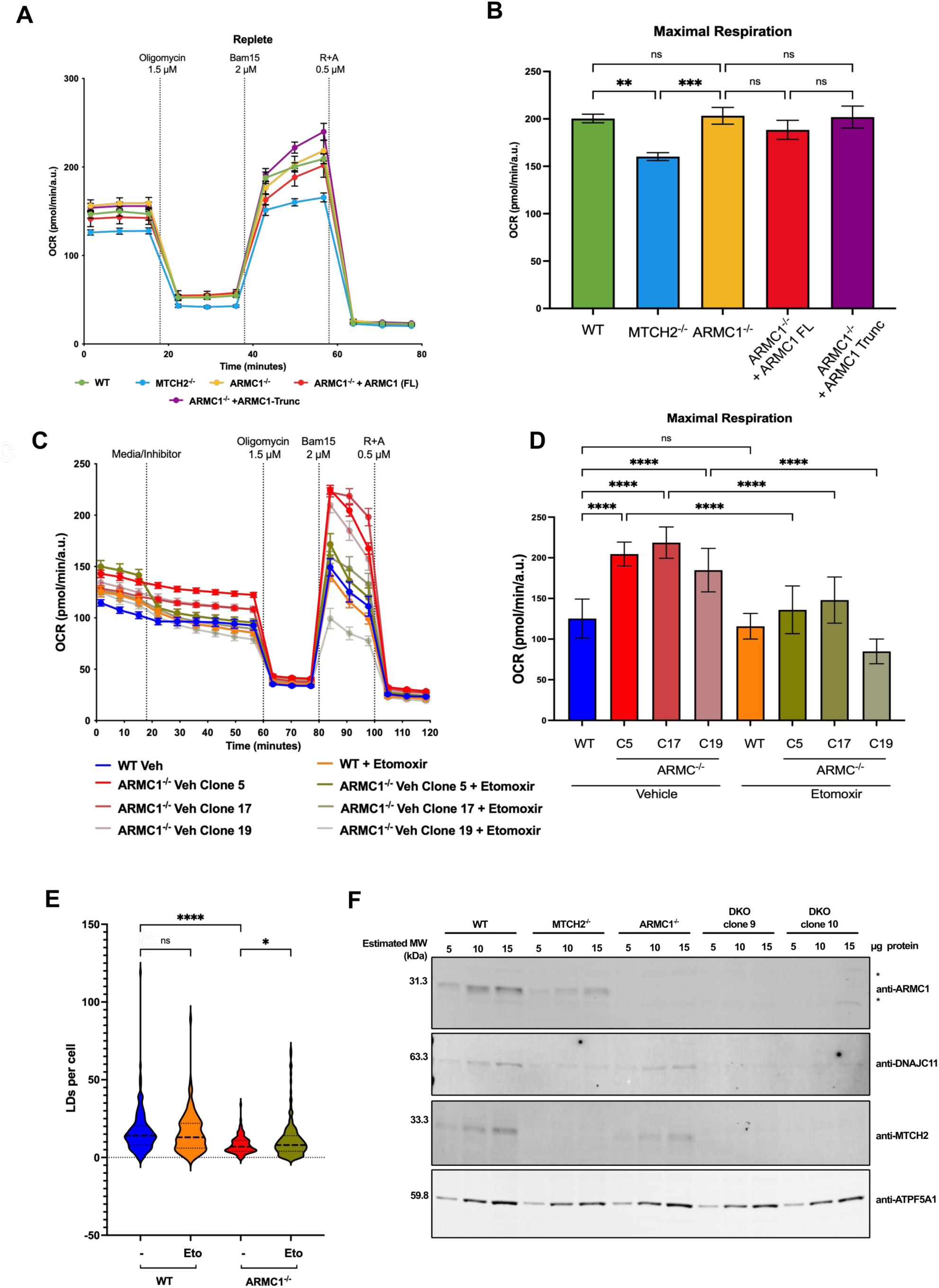
OCR levels under replete conditions and Etomoxir treatment with FAO activator. A) Oxygen consumption rates during replete conditions for HCT116 WT cells, MTCH2^-/-^, ARMC1^-/-^, ARMC1^-/-^ stable line expressing ARMC1 full length (FL), and an ARMC1^-/-^ stable line expressing ARMC1 with a 30 amino acid truncation in the C-terminus domain (Trunc). (B) Maximal respiratory capacity under replete conditions. Error bars mean ±SEM. Statistical significance was evaluated by two-way ANOVA followed by Tukey’s test. (C) Oxygen consumption rate during starvation under etomoxir (4 µM) or vehicle (Media) treatment for HCT116 WT cells and three different clones (Clone 5, 17, and 19) of ARMC1^-/-^ cells. (D) Maximal respiratory capacity from the conditions mentioned in (C). Error bars mean ±SEM. Statistical significance was evaluated by two-way ANOVA followed by Tukey’s test. (E) Violin plot for Lipid droplet (LDs) quantification under etomoxir treatment (4 µM) during starved conditions for WT and ARMC1^-/-^ cells. Statistical significance was evaluated by two-way ANOVA followed by Tukey’s test. Immunoblot for ARMC1 to evaluate the level of knockout in double mutants ARMC1^-/-^ / MTCH2^-/-^. *Non-specific bands. Significance is assessed by *P<0.05, **P<0.01, ***P<0.001, ****P<0.0001, and ns = not significant.

**Figure 5-1 supplement:**
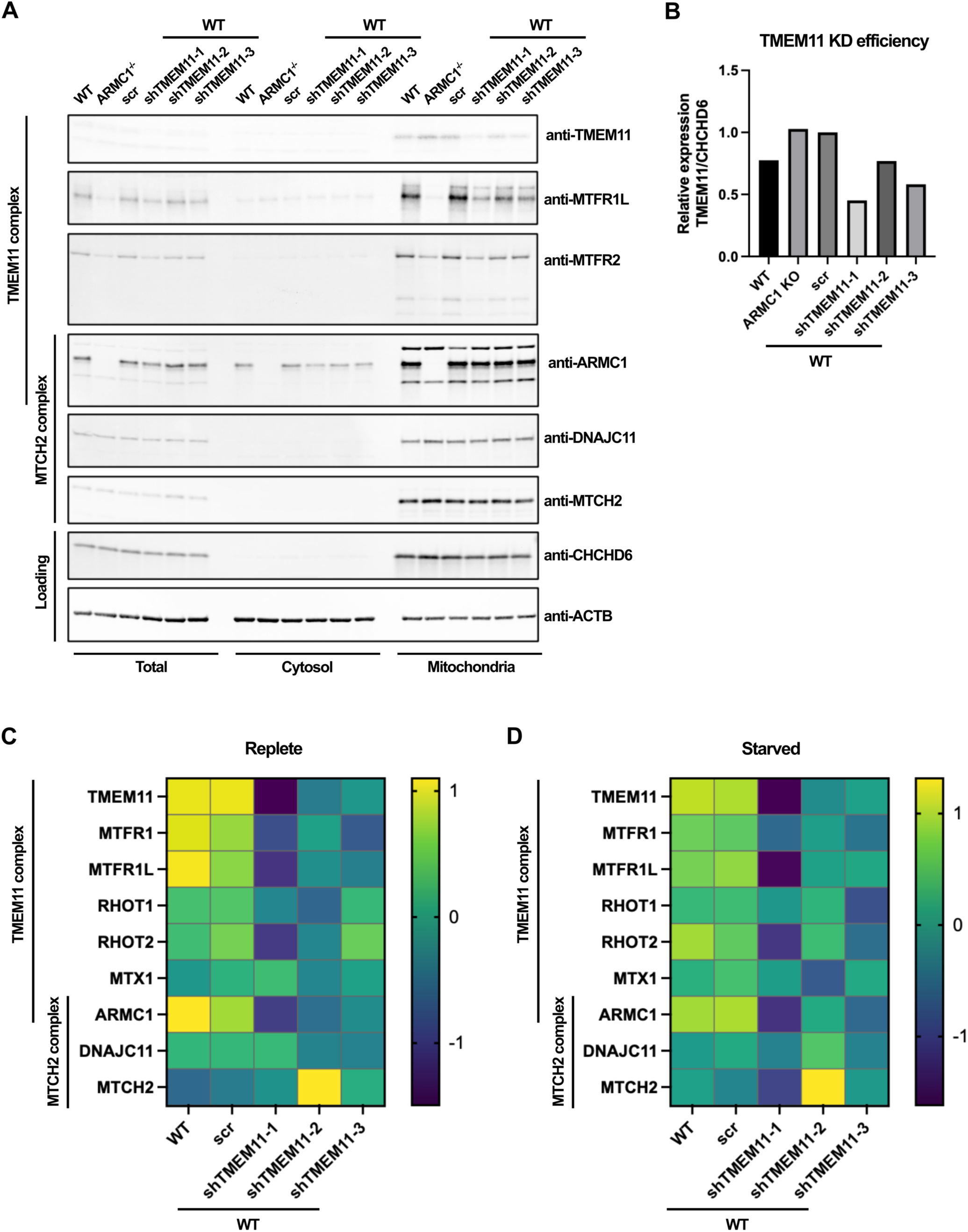
Measurement of TMEM11 KD efficiency and test of co-depletion of ARMC1-interacting proteins. (A) Knockdown efficiency of 3 different shRNAs targeting TMEM11 (top panel) and codependency of ARMC1-interacting proteins was evaluated by immunoblot of subcellular fractions. (B) Abundance of TMEM11 in mitochondrial fraction was quantified relative to CHCHD6. (C-D) Whole cell proteomics was performed on TMEM11 KD cell lines compared to WT and un-transduced controls to validate TMEM11 KD efficiency and further investigate co-depletion of ARMC1-interacting proteins with change in protein abundance depicted via heatmap of Z-score (n = 5 replicates/condition).

**Figure 5-2 supplement:**
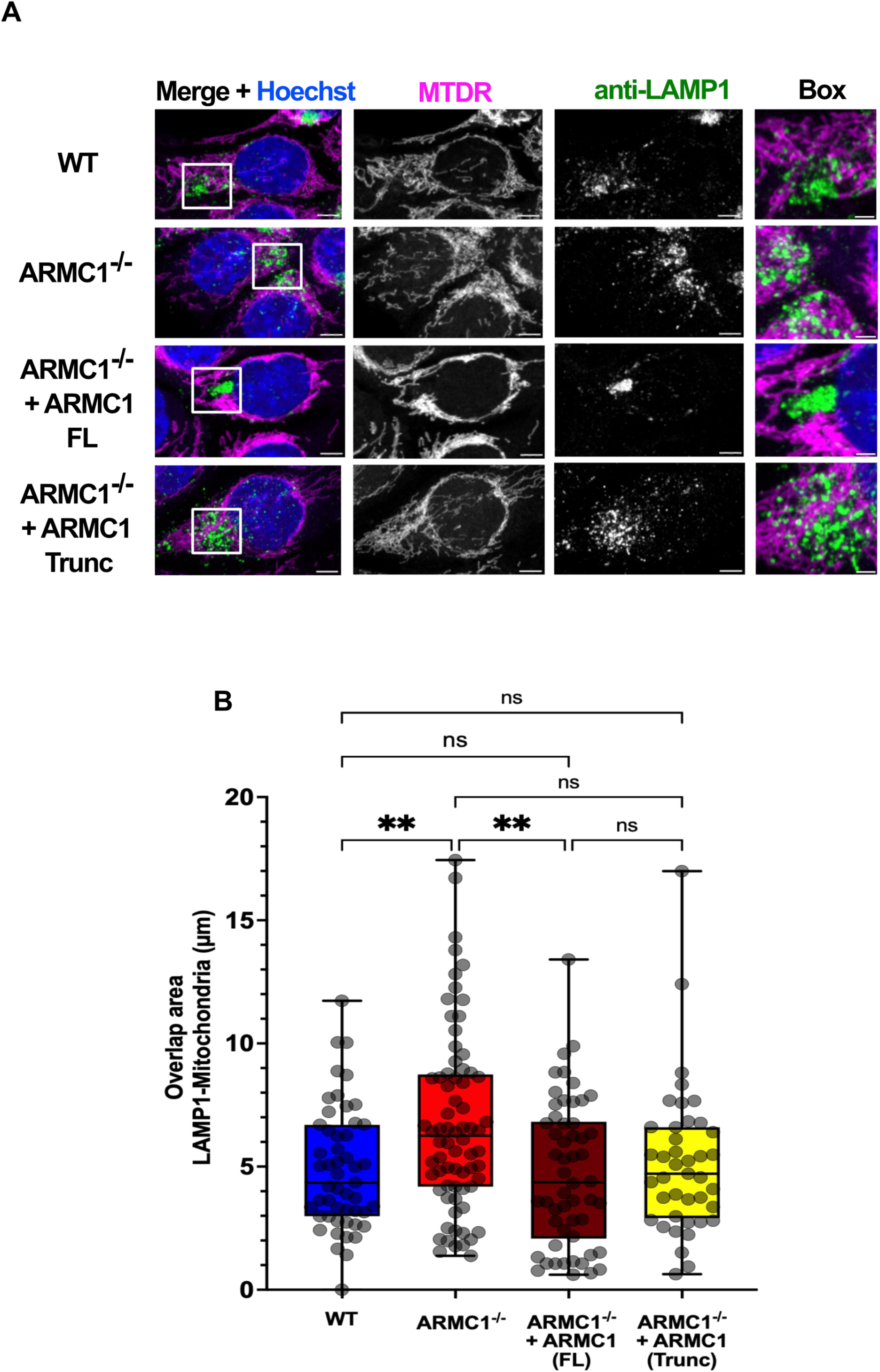
Subcellular localization of lysosomes assessed by LMAP1 immunofluorescence labeling. (A) Immunofluorescence for anti-LAMP1 and staining of Hoechst and Mitotracker Deep Red for WT, ARMC1^-/-^, ARMC1^-/-^ + ARMC1 (FL), and ARMC1^-/-^ + ARMC1 (Trunc) cells under starved conditions. Scale bars = 5 µm. The boxes showed a scale bar = 2 µm. (B) Quantification of the overlap area between mitochondria and LAMP1 for the cell lines mentioned in (A). Error bars mean ±SD. Statistical significance was assessed by one-way ANOVA followed by Tukey’s test. *P<0.05, **P<0.01, ***P<0.001, ****P<0.0001, and ns = not significant.

**Figure 5-3 supplement:**
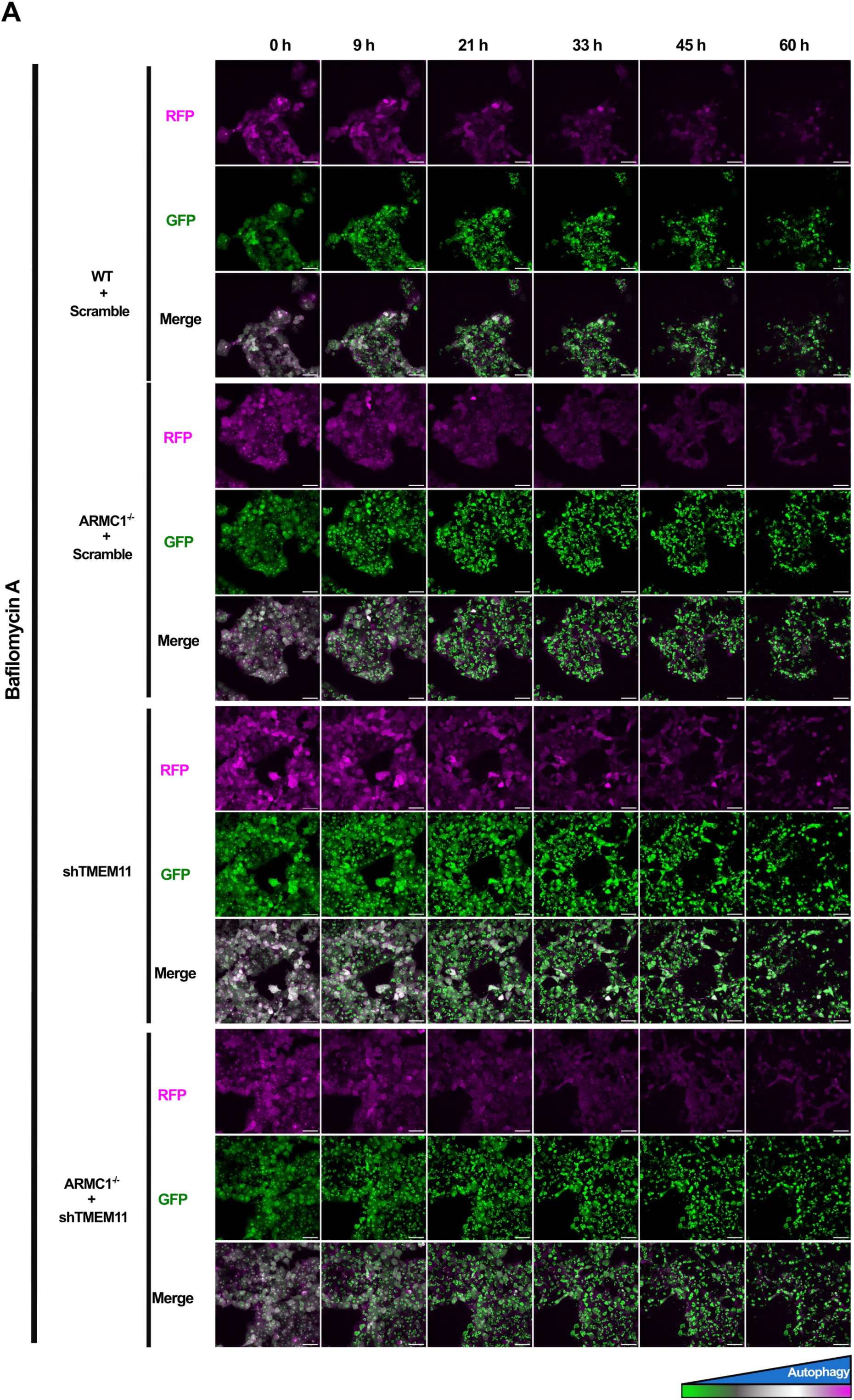
Autophagy flux measurement during nutrient-starved conditions with bafilomycin A1 treatment. (A) Inhibition of autophagy by Bafilomycin (100 nM). Representative images of GFP, RFP, and Merge for the detection of starvation-induced autophagy by microscopy for WT + Scramble, ARMC1^-/-^ + Scramble, shTMEM11, and ARMC1^-/-^ + shTMEM11 expressing RFP-LC3-GFP delivered by LNP. Scale bar = 50 µm.

**Figure 5-4 supplement:**
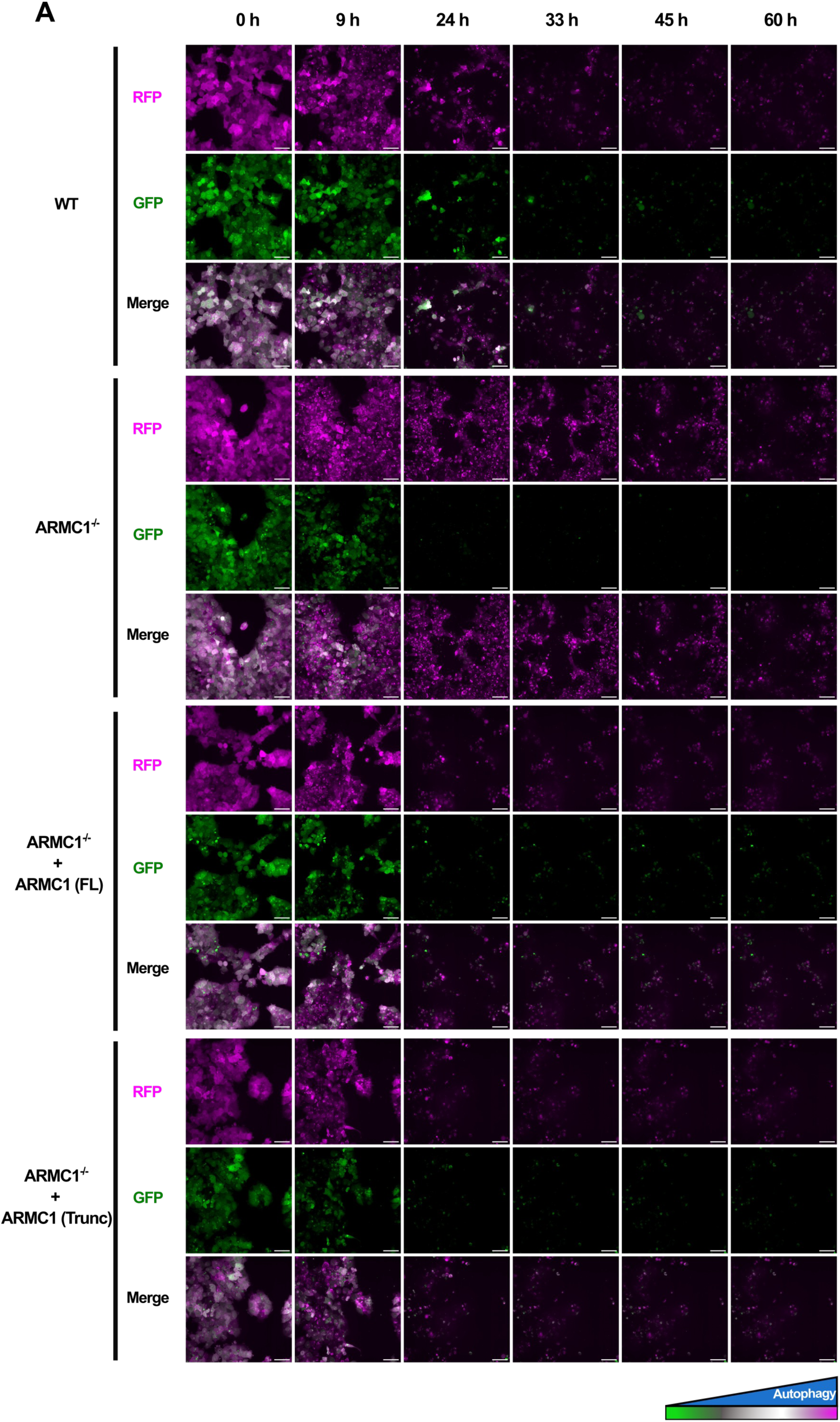
Autophagy flux measurement for ARMC1^-/-^ + ARMC1 (FL) and ARMC1^-/-^ + ARMC1 (Trunc) under starvation conditions. (A) Representative images of GFP, RFP, and Merge for the detection of starvation-induced autophagy by microscopy for WT, ARMC1^-/-^, ARMC1^-/-^ + ARMC1 (FL), ARMC1^-/-^ + ARMC1 (Trunc) expressing RFP-LC3-GFP delivered by LNP. Scale bar = 50 µm.

**Figure 5-5 supplement:**
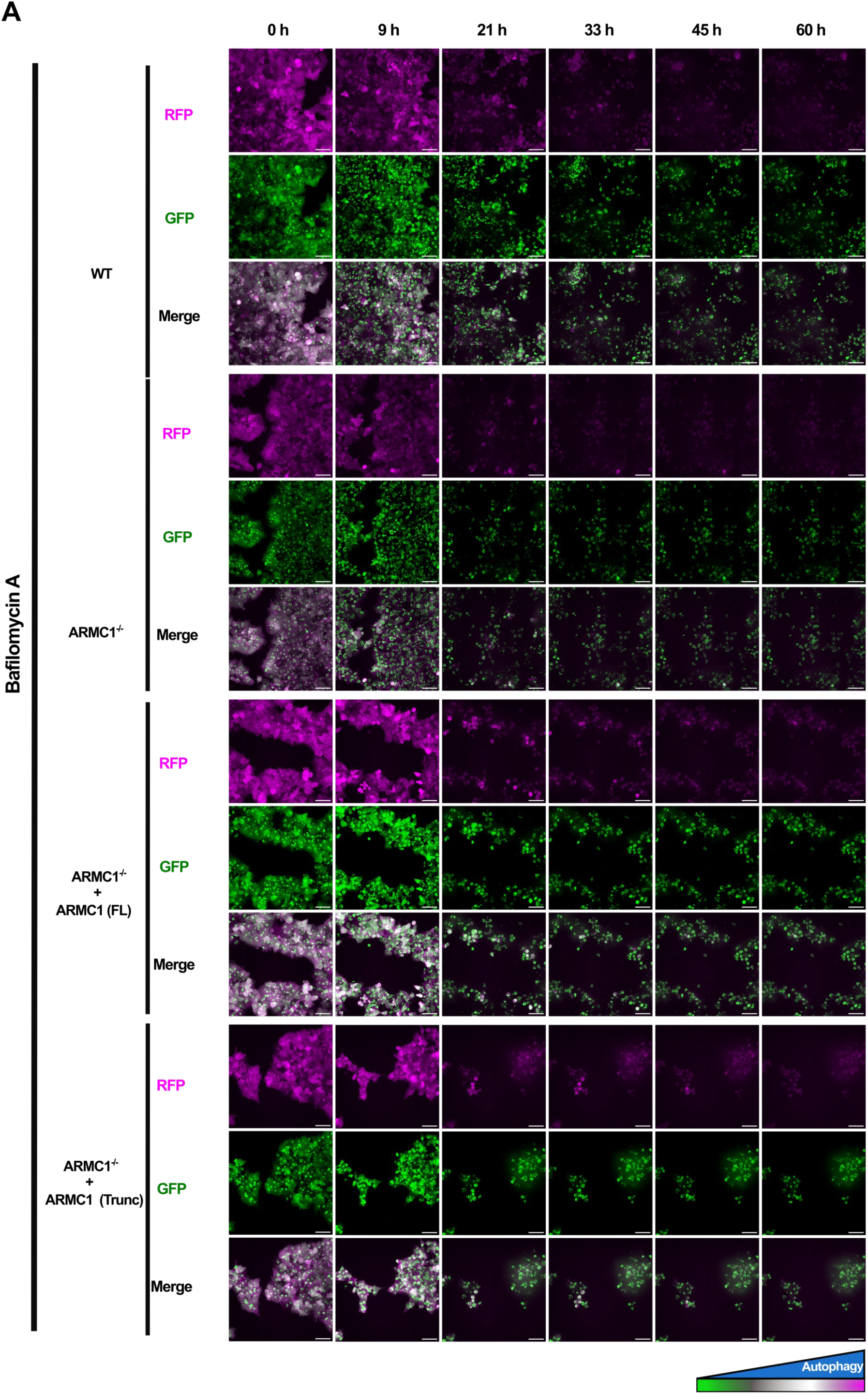
Autophagy flux measurement for ARMC1^-/-^ + ARMC1 (FL) and ARMC1^-/-^ + ARMC1 (Trunc) under starvation conditions with bafilomycin A1. (A) Inhibition of autophagy by Bafilomycin (100 nM). Representative images of GFP, RFP, and Merge for the detection of starvation-induced autophagy by microscopy for WT, ARMC1^-/-^, ARMC1^-/-^ + ARMC1 (FL), ARMC1^-/-^ + ARMC1 (Trunc) expressing RFP-LC3-GFP delivered by LNP. Scale bar = 50 µm.

## References

1. Gao M, An DN, Parks JM, Skolnick J. AF2Complex predicts direct physical interactions in multimeric proteins with deep learning. Nat Commun. 2022;13(1):1744. doi:10.1038/s41467-022-29394-2

2. Bennett CF, Latorre-Muro P, Puigserver P. Mechanisms of mitochondrial respiratory adaptation. Nat Rev Mol Cell Biol. 2022;23(12):817–835. doi:10.1038/s41580-022-00506-6

3. Spinelli JB, Haigis MC. The multifaceted contributions of mitochondria to cellular metabolism. Nat Cell Biol. 2018;20(7):745–754. doi:10.1038/s41556-018-0124-1

4. Gambardella J, Lombardi A, Santulli G. Metabolic Flexibility of Mitochondria Plays a Key Role in Balancing Glucose and Fatty Acid Metabolism in the Diabetic Heart. Diabetes. 2020;69(10):2054–2057. doi:10.2337/dbi20-0024

5. Rahimi Y, Camporez JPG, Petersen MC, et al. Genetic activation of pyruvate dehydrogenase alters oxidative substrate selection to induce skeletal muscle insulin resistance. Proc Natl Acad Sci. 2014;111(46):16508–16513. doi:10.1073/pnas.1419104111

6. Abdullah MO, Zeng RX, Margerum CL, et al. Mitochondrial hyperfusion via metabolic sensing of regulatory amino acids. Cell Rep. 2022;40(7):111198. doi:10.1016/j.celrep.2022.111198

7. Gomes LC, Benedetto GD, Scorrano L. During autophagy mitochondria elongate, are spared from degradation and sustain cell viability. Nat Cell Biol. 2011;13(5):589–598. doi:10.1038/ncb2220

8. Rambold AS, Cohen S, Lippincott-Schwartz J. Fatty Acid Trafficking in Starved Cells: Regulation by Lipid Droplet Lipolysis, Autophagy, and Mitochondrial Fusion Dynamics. Dev Cell. 2015;32(6):678–692. doi:10.1016/j.devcel.2015.01.029

9. Rambold AS, Kostelecky B, Elia N, Lippincott-Schwartz J. Tubular network formation protects mitochondria from autophagosomal degradation during nutrient starvation. Proc National Acad Sci. 2011;108(25):10190–10195. doi:10.1073/pnas.1107402108

10. Labbé K, Mookerjee S, Vasseur ML, Gibbs E, Lerner C, Nunnari J. The modified mitochondrial outer membrane carrier MTCH2 links mitochondrial fusion to lipogenesis. J Cell Biol. 2021;220(11):e202103122. doi:10.1083/jcb.202103122

11. Ruprecht JJ, Kunji ERS. The SLC25 Mitochondrial Carrier Family: Structure and Mechanism. Trends Biochem Sci. 2020;45(3):244–258. doi:10.1016/j.tibs.2019.11.001

12. McKenna MJ, Kraus F, Coelho JPL, et al. ARMC1 partitions between distinct complexes and assembles MIRO with MTFR to control mitochondrial distribution. Sci Adv. 2025;11(15):eadu5091. doi:10.1126/sciadv.adu5091

13. Stevens TA, Luo Z, Lee C, et al. Structural evolution of the MTCH family of mitochondrial insertases. Published online 2026. doi:10.64898/2026.02.17.705849

14. Bartoš L, Menon AK, Vácha R. Insertases scramble lipids: Molecular simulations of MTCH2. Structure. 2024;32(4):505–510.e4. doi:10.1016/j.str.2024.01.012

15. Li D, Rocha-Roa C, Schilling MA, Reinisch KM, Vanni S. Lipid scrambling is a general feature of protein insertases. Proc Natl Acad Sci. 2024;121(17):e2319476121. doi:10.1073/pnas.2319476121

16. Guna A, Stevens TA, Inglis AJ, et al. MTCH2 is a mitochondrial outer membrane protein insertase. Science. 2022;378(6617):317–322. doi:10.1126/science.add1856

17. Wagner F, Kunz TC, Chowdhury SR, et al. Armadillo repeat-containing protein 1 is a dual localization protein associated with mitochondrial intermembrane space bridging complex. Plos One. 2019;14(10):e0218303. doi:10.1371/journal.pone.0218303

18. Xu W, Kimelman D. Mechanistic insights from structural studies of β-catenin and its binding partners. J Cell Sci. 2007;120(19):3337–3344. doi:10.1242/jcs.013771

19. Mattie S, Riemer J, Wideman JG, McBride HM. A new mitofusin topology places the redox-regulated C terminus in the mitochondrial intermembrane space. J Cell Biol. 2018;217(2):507–515. doi:10.1083/jcb.201611194

20. Aksoyoglu MA, Podgornik R, Bezrukov SM, Gurnev PA, Muthukumar M, Parsegian VA. Size-dependent forced PEG partitioning into channels: VDAC, OmpC, and α-hemolysin. Proc Natl Acad Sci. 2016;113(32):9003–9008. doi:10.1073/pnas.1602716113

21. Guarani V, McNeill EM, Paulo JA, et al. QIL1 is a novel mitochondrial protein required for MICOS complex stability and cristae morphology. Elife. 2015;4:e06265. doi:10.7554/elife.06265

22. Arafeh R, Shibue T, Dempster JM, Hahn WC, Vazquez F. The present and future of the Cancer Dependency Map. Nat Rev Cancer. 2025;25(1):59–73. doi:10.1038/s41568-024-00763-x

23. Zhu Y, Akkaya KC, Ruta J, et al. Cross-link assisted spatial proteomics to map sub-organelle proteomes and membrane protein topologies. Nat Commun. 2024;15(1):3290. doi:10.1038/s41467-024-47569-x

24. Dixon AS, Schwinn MK, Hall MP, et al. NanoLuc Complementation Reporter Optimized for Accurate Measurement of Protein Interactions in Cells. ACS Chem Biol. 2016;11(2):400–408. doi:10.1021/acschembio.5b00753

25. Decker ST, Funai K. Mitochondrial membrane lipids in the regulation of bioenergetic flux. Cell Metab. 2024;36(9):1963–1978. doi:10.1016/j.cmet.2024.07.024

26. Buzaglo-Azriel L, Kuperman Y, Tsoory M, et al. Loss of Muscle MTCH2 Increases Whole-Body Energy Utilization and Protects from Diet-Induced Obesity. Cell Rep. 2016;14(7):1602–1610. doi:10.1016/j.celrep.2016.01.046

27. Fischer JA, Monroe TO, Pesce LL, et al. Opposing effects of genetic variation in MTCH2 for obesity versus heart failure. Hum Mol Genet. 2022;32(1):15–29. doi:10.1093/hmg/ddac176

28. Chourasia S, Petucci C, Shoffler C, et al. MTCH2 controls energy demand and expenditure to fuel anabolism during adipogenesis. EMBO J. Published online 2025:1–32. doi:10.1038/s44318-024-00335-7

29. Moon SH, Liu X, Jenkins CM, Dilthey BG, Patti GJ, Gross RW. Etomoxir-carnitine, a novel pharmaco-metabolite of etomoxir, inhibits phospholipases A2 and mitochondrial respiration. J Lipid Res. 2024;65(9):100611. doi:10.1016/j.jlr.2024.100611

30 . Spurway TD, Pogson CI, Sherratt HSA, Agius L. Etomoxir, sodium 2-[6-(4-chlorophenoxy)hexyl]oxirane-2-carboxylate, inhibits triacylglycerol depletion in hepatocytes and lipolysis in adipocytes. FEBS Lett. 1997;404(1):111–114. doi:10.1016/s0014-5793(97)00103-8

31. McCune SA, Harris RA. Mechanism responsible for 5-(tetradecyloxy)-2-furoic acid inhibition of hepatic lipogenesis. J Biol Chem. 1979;254(20):10095–10101. doi:10.1016/s0021-9258(19)86677-2

32. Ngo J, Choi DW, Stanley IA, et al. Mitochondrial morphology controls fatty acid utilization by changing CPT1 sensitivity to malonyl-CoA. EMBO J. 2023;42(11):e111901. doi:10.15252/embj.2022111901

33. Gok MO, Connor OM, Wang X, et al. The outer mitochondrial membrane protein TMEM11 demarcates spatially restricted BNIP3/BNIP3L-mediated mitophagy. J Cell Biol. 2023;222(4):e202204021. doi:10.1083/jcb.202204021

34. Cho NH, Cheveralls KC, Brunner AD, et al. OpenCell: Endogenous tagging for the cartography of human cellular organization. Science. 2022;375(6585):eabi6983. doi:10.1126/science.abi6983

35. Wu C, Wang T, Ghosh A, et al. MTCH2 modulates CPT1 activity to regulate lipid metabolism of adipocytes. Nat Commun. 2025;16(1):8831. doi:10.1038/s41467-025-63880-7

36. Vasseur ML, Friedman J, Jost M, et al. Genome-wide CRISPRi screening identifies OCIAD1 as a prohibitin client and regulatory determinant of mitochondrial Complex III assembly in human cells. Elife. 2021;10:e67624. doi:10.7554/elife.67624

37. Vasseur ML, Friedman J, Jost M, et al. Genome-wide CRISPRi screening identifies OCIAD1 as a prohibitin client and regulatory determinant of mitochondrial Complex III assembly in human cells. Elife. 2021;10:e67624. doi:10.7554/elife.67624

38. Ramírez-Pardo I, Campanario S, Chavda B, et al. Chaperone-mediated autophagy sustains muscle stem cell regenerative functions but declines with age. Nat Metab. Published online 2025:1–18. doi:10.1038/s42255-025-01411-w

39. Fleming J, Magana P, Nair S, et al. AlphaFold Protein Structure Database and 3D-Beacons: New Data and Capabilities. J Mol Biol. 2025;437(15):168967. doi:10.1016/j.jmb.2025.168967

40. Ahdritz G, Bouatta N, Floristean C, et al. OpenFold: retraining AlphaFold2 yields new insights into its learning mechanisms and capacity for generalization. Nat Methods. 2024;21(8):1514–1524. doi:10.1038/s41592-024-02272-z

41. Evans R, O’Neill M, Pritzel A, et al. Protein complex prediction with AlphaFold-Multimer. Biorxiv. Published online 2022:2021.10.04.463034. doi:10.1101/2021.10.04.463034

42. Passaro S, Corso G, Wohlwend J, et al. Boltz-2: Towards Accurate and Efficient Binding Affinity Prediction. bioRxiv. Published online 2025:2025.06.14.659707. doi:10.1101/2025.06.14.659707

43. Wohlwend J, Corso G, Passaro S, et al. Boltz-1 Democratizing Biomolecular Interaction Modeling. bioRxiv. Published online 2025:2024.11.19.624167. doi:10.1101/2024.11.19.624167

44. Mirdita M, Schütze K, Moriwaki Y, Heo L, Ovchinnikov S, Steinegger M. ColabFold: making protein folding accessible to all. Nat Methods. 2022;19(6):679–682. doi:10.1038/s41592-022-01488-1

45. Souza PCT, Alessandri R, Barnoud J, et al. Martini 3: a general purpose force field for coarse-grained molecular dynamics. Nat Methods. 2021;18(4):382–388. doi:10.1038/s41592-021-01098-3

46. Kroon PC, Grünewald F, Barnoud J, et al. Martinize2 and Vermouth provide a unified framework for molecular topology generation. eLife. 2025;12:RP90627. doi:10.7554/elife.90627

47. Abraham MJ, Murtola T, Schulz R, et al. GROMACS: High performance molecular simulations through multi-level parallelism from laptops to supercomputers. SoftwareX. 2015;1:19–25. doi:10.1016/j.softx.2015.06.001

48. Huang J, Rauscher S, Nawrocki G, et al. CHARMM36m: an improved force field for folded and intrinsically disordered proteins. Nat Methods. 2017;14(1):71–73. doi:10.1038/nmeth.4067

49. Klauda JB, Venable RM, Freites JA, et al. Update of the CHARMM All-Atom Additive Force Field for Lipids: Validation on Six Lipid Types. J Phys Chem B. 2010;114(23):7830-7843. doi:10.1021/jp101759q

50. Gowers R, Linke M, Barnoud J, et al. MDAnalysis: A Python Package for the Rapid Analysis of Molecular Dynamics Simulations. Proc 15th Python Sci Conf. Published online 2016:98-105. doi:10.25080/majora-629e541a-00e

51. Combe CW, Graham M, Kolbowski L, Fischer L, Rappsilber J. xiVIEW: Visualisation of Crosslinking Mass Spectrometry Data. J Mol Biol. 2024;436(17):168656. doi:10.1016/j.jmb.2024.168656

52. Shiiba I, Ito N, Oshio H, et al. ER-mitochondria contacts mediate lipid radical transfer via RMDN3/PTPIP51 phosphorylation to reduce mitochondrial oxidative stress. Nat Commun. 2025;16(1):1508. doi:10.1038/s41467-025-56666-4

39. Nguyen, T. B. et al. DGAT1-Dependent Lipid Droplet Biogenesis Protects Mitochondrial Function during Starvation-Induced Autophagy. Dev. Cell 42, 9–21.e5 (2017). doi:10.1016/j.devcel.2017.06.003

